# Full-length transcriptome maps of reef-building coral illuminate the molecular basis of calcification, symbiosis, and circadian genes

**DOI:** 10.1101/2022.03.23.485437

**Authors:** Tingyu Han, Xin Liao, Yunchi Zhu, Yunqing Liu, Na Lu, Yixin Li, Zhuojun Guo, J.-Y. Chen, Chunpeng He, Zuhong Lu

## Abstract

**Background:** Reef-building corals are critical species for sustaining coral reefs and are highly threatened by global climate change. However, relevant transcriptomic data largely rely on short-read sequencing, which severely limits the understanding of coral molecular mechanisms and leaves many important biological questions unresolved.

**Results:** We sequenced the full-length transcriptomes of four common and frequently dominant reef-building corals, including two Robusta clade species, *Pocillopora damicornis* and *Pocillopora verrucosa*, and two Complexa clade species, *Acropora muricata* and *Montipora foliosa,* using the PacBio Sequel II platform. We obtained information on gene functions, structures and expression profiles. Among them, a comparative analysis of biomineralization-related genes provided insights into the molecular basis of coral skeletal density. The gene expression profiles of the symbiote Symbiodiniaceae were also isolated and annotated from the holobiont sequence data; these profiles showed more highly convergent traits related to gene structure and expression level than those of coral hosts. Interestingly, we observed that intracellular algal cells share some evolutionary convergence between intracellular symbiosis in corals and intracellular digestion in amphioxus. Finally, a phylogenetic analysis of key circadian clock genes among 27 evolutionarily representative species indicated that there are four key members in early metazoans, including *cry* genes; *Clock* or *Npas2*; *cyc* or *Arntl*; and *tim*, while *per*, as the fifth member, occurs in Bilateria.

**Conclusions:** Our work overcomes the incompleteness of short-read sequencing and illuminates the molecular basis of calcification, symbiosis, and circadian genes, thus providing a foundation for further work on the manipulation of skeleton production or symbiosis to promote the survival of these important organisms.

## Background

Coral reefs are among the most productive and biodiverse ecosystems on earth [1], and they provide survival habitats for approximately 30% of marine life [2–5]. Approximately 500 million people worldwide depend on these reefs, demonstrating their great ecological and economic value [6–10]. The scleractinian corals that mainly produce coral reefs constitute one of the most primitive metazoan branches [2]. They show the hydrozoan body plan typical of Phylum *Cnidaria* [9] and retain many ancient gene features of ancestral metazoans [11], providing an important genetic background for studying the evolutionary origin of metazoans and bilaterians [12, 13]. Currently, our understanding of the molecular biology of reef-building corals largely derives from omics sequencing, and these data have provided crucial information on coral calcification [14, 15], symbiosis [16–21], heat stress [22–24], acid stress [25, 26], cnidocytes [27, 28], collagen secretion [29], immunity [28, 30–32], budding [33, 34] and circadian clocks [35–37]. Despite these advances, many open questions remain.

The greatest contribution of scleractinian corals to marine biota and ecosystems is the deposition of reefs via biomineralization in calicoblasts [28], which form skeletons. The reef-building coral skeleton is built through the continuous deposition of aragonite [38], which is formed by a mineral fraction consisting of calcium carbonate and an organic matrix molecule fraction that includes carbohydrates, lipids and proteins [14]. Current reports indicate that the main components used for calcification include calcium ATPase, carbonic anhydrase (CA), bicarbonate transporter (i.e., solute carrier 4 [SLC4] and solute carrier 26 [SLC26]) and core skeleton organic matrix proteins (SOMPs) (e.g., acid-rich protein, uncharacterized skeletal organic matrix proteins [USOMPs], galaxins and alpha IV collagen) [14, 15, 38–49]. However, due to the lack of full-length sequences for these genes, in-depth biological studies such as gene function reconstruction or gene editing cannot be performed. Moreover, the expression levels of these genes are rarely reported, limiting precise studies of their regulation under environmental stress. In addition, the relationship between coral skeletal density and the expression levels of calcification genes has been rarely discussed.

Phagocytosis targeting primitive algae was a prerequisite for the development of food webs involving multicellular animals, as well as the origin of mitochondria and chloroplasts [50–53]. Additionally, paleontological [51], geochemical [50, 51] and molecular clock [50–52] evidence suggests that various forms of eukaryotic predation occurred during the Neoproterozoic Era (1000-541 million years ago), from the proliferation of marine algae to the origin of multicellular animal clades. At that time, increasing algal abundance created food webs with more efficient nutrient and energy transfer, driving shifts in ecosystems towards larger and increasingly complex organisms [50]. This effect is recorded in the appearance of sponges and the subsequent radiation of eumetazoans in the Ediacaran period [50]. Currently, most hosts of intracellular symbionts belong to groups with polyp body plans, such as hydra, soft corals, anemones and reef-building corals, and Mollusca, such as giant clam [16-21, 27, 54-56]. Among these groups, reef-building corals have particularly widespread and extensive endosymbiosis (with Symbiodiniaceae). Beyond intracellular symbionts, another interesting form occurred in amphioxus (*Branchiostoma belcheri*), a primitive deuterostome invertebrate that degrades various small-scale algae by phagocytic intracellular digestion within phagosomes [57]. Thus, there are two major forms of interaction with algae, intracellular symbiosis and intracellular digestion, which differ in terms of how algae are acquired and whether intracellular algae are ‘reared’ or ‘destroyed’. Intracellular symbiosis is generally thought to depend on molecular recognition for the acquisition of algae [58], while intracellular digestion involves direct preying by cell cilia [57]. Symbiodiniaceae, similar to other dinoflagellates, must produce low doses of dinotoxins as a defence against predation or as metabolic byproducts; hence, host cells must carry out certain reactions to reduce algal toxicity in an intracellular symbiotic system, similar to intracellular digestion [59–61]. An incompatible dynamic equilibrium relationship with intracellular algae may lead to host cell death, and reducing immune rejection is a method for alleviating immune conflict and benefiting from these algae during intracellular symbioses or digestion [54]. Because both mechanisms involve interaction with intracellular algae and evolved from the early stage of algal phagocytosis, it is expected that there should be some overlap between them; however, the extent of this overlap is unclear. Such connections should be revealed by omics comparisons across host species.

Reef-building corals show obvious circadian rhythms; interestingly, the circadian clock system and related genes have not been deeply investigated, although they play important roles in sustaining healthy coral growth [35–37]. In a marine sponge, *Amphimedon queenslandica*, which is a representative of the oldest extant animal phyletic lineage, some molecular components of circadian clock are shared with eumetazoans, reflecting a core clock present in the last common animal ancestor [62, 63]. At the basic molecular level, the operations of the circadian clock system and biological rhythm behaviour are regulated through a conserved negative transcription-translation feedback loop, which is controlled by five key gene families [64]. Cryptochromes (CRYs) are a class of flavoproteins that can detect blue light [65]. Generally, circadian locomotor output cycles kaput (CLOCK) and its homologous protein Neuronal PAS domain-containing protein 2 (NPAS2), as well as Cycle (CYC) and its homologous protein Brain and muscle ARNT-like (BMAL), act as positive regulatory factors, while Period (PER) and Timeless (TIM) act as negative regulatory factors [66]. However, the specific connections among these families vary across clades. In *Drosophila*, CRY regulates the circadian clock in a light-dependent manner [67], whereas in mice, CRY1 and CRY2 act as light-independent inhibitors of the CYC-CLOCK component [68]. However, in some invertebrates, such as monarch butterflies, CRYs exhibit both *Drosophila*-like and mammal-like functions, providing evidence of an ancestral clock gene regulation state [69, 70]. Three *cry* genes and fifteen other homologous bilaterian circadian clock genes have been found in the *Acropora digitifera* coral genome [35], and 24 circadian genes in the *Acropora millepora* coral transcriptome have orthologues in bilaterian species, including 6 insect and 18 mammalian genes [36, 37]. The composition and related phylogenetic pattern of circadian genes among the actinozoan lineage remain unclear; more omics data is needed to elucidate them.

On a global scale, *Pocillopora damicornis* (*P. damicornis*), *Acropora muricata* (*A. muricata*) and *Montipora foliosa* (*M. foliosa*) are common dominant reef-building corals in the Indo-Pacific region, and *Pocillopora verrucosa* (*P. verrucosa*) is a typical reef-front stony coral that protects fringing reefs [71–74]. These four corals play pivotal roles in the Indo-Pacific coral reef ecological system. Recently, increasing amounts of sequencing data derived from these reef-building corals have been deposited in public databases (see Table 1 and Additional file 2: Table S1). However, all currently available sequences have been generated using short-read (50-300 bp) sequencing approaches, leaving incomplete information and sequence splicing errors [75–77]. Critically, although some high-quality sequences have been published for both corals themselves [23, 24, 30–32, 78–93] and their associated microorganisms [33, 94–113], the reliance on short reads has prevented the precise delimitation of the gene expression profiles of reef-building corals and their endosymbiotic Symbiodiniaceae. Recent advances in long-read sequencing technology (e.g., PacBio Sequel II) have made it possible to obtain large amounts of full-length transcript data from many organisms and tissues [114–116]. In principle, such data should permit the characterization of all expressed transcripts as complete, contiguous mRNA sequences from the transcription start site to the transcription end site; in turn, this enables the more accurate and efficient analysis of the full spectrum of gene expression profile information, including data on gene expression, alternative splicing, gene fusion, expression regulation, coding sequences (CDSs), and protein structure [117–120].

**Table 1.** Sequence data of four reef-building corals in the NCBI database.

| NO. | BioProject | Data Volume,<br>Gb | Data<br>Volume,<br>Mbytes | Library<br>Strategy | Platform | Model |
| --- | --- | --- | --- | --- | --- | --- |
| *P. damicornis* |  |  |  |  |  |  |
| 1 | PRJNA227785 | 27 | 17496 | RNA-Seq | Illumina | Genome<br>Analyzer IIx |
| 2 | PRJNA299443 | 24 | 10069 | RNA-Seq | Illumina | HiSeq 4000 |
| 3 | PRJNA327142 | 10 | 3750 | RNA-Seq | Illumina | HiSeq 4000 |
| 4 | PRJNA433950 | 60 | 23545 | RNA-Seq | Illumina | HiSeq X Ten |
| 5 | PRJNA435620 | 67 | 26334 | RNA-Seq | Illumina | HiSeq X Ten |
| 6 | PRJNA306839 | 18 | 8574 | RNA-Seq | Illumina | MiSeq |
| 7 | PRJEB26650 | 45 | 21254 | RNA-Seq | Illumina | HiSeq 2500 |
| 8 | **PRJNA544778** | **1348** | **0.41T** | **RNA-Seq** | **PacBio**<br>**Illumina** | **Sequel II**<br>**HiSeq X Ten** |
| 9 | PRJNA545379 | 67 | 27278 | RNA-Seq | Illumina | HiSeq 2000 |
| 10 | PRJNA611041 | 258 | 90416 | RNA-Seq | Illumina | HiSeq 1500 |
| *P. verrucosa* |  |  |  |  |  |  |
| 1 | PRJNA552592 | 15 | 7782 | RNA-Seq | Illumina | HiSeq 2500 |
| 2 | PRJNA551401 | 108 | 47540 | RNA-Seq<br>WGS | Illumina | HiSeq4000<br>HiSeq2500 |
| 3 | **PRJNA544778** | **1348** | **0.41T** | **RNA-Seq** | **PacBio**<br>**Illumina** | **Sequel II**<br>**HiSeq X Ten** |
| *A. muricata* |  |  |  |  |  |  |
| 1 | **PRJNA544778** | **1348** | **0.41T** | **RNA-Seq** | **PacBio**<br>**Illumina** | **Sequel II**<br>**HiSeq X Ten** |
| 2 | PRJDB8519 | 1785 | 1.09T | WGS | Illumina | HiSeq 2500 |
| 3 | PRJDB5633 | 110 | 63481 | WGS | Illumina | HiSeq 2500 |
| *M. foliosa* |  |  |  |  |  |  |
| 1 | **PRJNA544778** | **1348** | **0.41T** | **RNA-Seq** | **PacBio**<br>**Illumina** | **Sequel II**<br>**HiSeq X Ten** |

Only datasets of ≥ 10 Gb coral sequences (i.e., excluding Symbiodiniaceae sequences) are included. The bolded lines indicate data generated in this paper.

To address these questions, we applied a full-length transcriptomic isoform sequencing (Iso-Seq) strategy using the PacBio Sequel II sequencing platform and quantitative gene expression analysis using the Illumina HiSeq X Ten sequencing platform to produce transcriptome maps for the four aforementioned important reef-building corals and their endosymbiotic Symbiodiniaceae. Based on the gene expression profiles, we performed a deeper analysis of several important physiological features of reef-building corals, including biomineralization-related gene groups, circadian gene families, and coral and Symbiodiniaceae factors related to the endosymbiotic interaction. These results enhance our understanding of the molecular biology and ecology of reef-building corals, which may aid in the recovery of marine ecosystems, and provide insights into the evolution of the circadian circuitry and holobiosis.

## Results

### Full-length transcriptome sequencing and data processing

Based on the standard processes of the PacBio Sequel II sequencing platform, the full-length raw transcriptomic sequencing data (polymerase reads) of four coral holobionts obtained included 25.34 Gb in *P. damicornis*, 27.8 Gb in *P. verrucosa*, 22.32 Gb in *A. muricata* and 21.44 Gb in *M. foliosa* (Table 2; Additional file 3: Table S2). These data were filtered to ensure quality and reliability, including removing reads of less than 50 bp in length and adapter sequences (for details and statistics, see Table 2; Additional file 1: Fig. S1, 2; Additional file 3: Table S2). The gene expression profiles of corals and their symbiotic Symbiodiniaceae (whose transcripts will be analysed separately in a later subsection) were separated based on an alignment to previously published sequences. In terms of the corals themselves, there were 20609 transcripts and 14167 unigenes (N50 = 2954 bp) in *P. damicornis*, 24174 transcripts and 12822 unigenes (N50 = 2313 bp) in *P. verrucosa*, 31242 transcripts and 13800 unigenes (N50 = 2126 bp) in *A. muricata*, and 25460 transcripts and 10905 unigenes (N50 = 1678 bp) in *M. foliosa* (Table 2; Additional file 3: Table S2). Unigene lengths were concentrated in the range of 1-3 kbp, with few unigenes of less than 1 kbp, verifying that the low-quality sequencing data had been filtered out (Additional file 1: Fig. S1b; Additional file 3: Table S2.8). A previous study has reported that there were 297221 assembled transcripts (N50 = 1831 bp) and 209337 unigenes (N50 = 1435 bp) in *P. damicornis* using a short-read sequencing approach [80]. However, it is obvious that PacBio sequencing can obtain fewer redundant sequences and higher quality sequences. Then, to evaluate the level of redundancy in the data, we examined the unigene-to-transcript ratio. The percentage of unigenes with a one-to-one unigene-to-transcript ratio was 73.71% (*P. damicornis*), 64.75% (*P. verrucosa*), 57.15% (*A. muricata*) and 57.26% (*M. foliosa*), and the percentage of unigenes with a one-to-two unigene-to-transcript ratio was 17.44% (*P. damicornis*), 17.52% (*P. verrucosa*), 18.41% (*A. muricata*) and 17.18% (*M. foliosa*) (detailed in Additional file 3: Table S2.9; Additional file 1: Fig. S1c). The sum of the two ratios was more than 74% in all samples, indicating that data redundancy was largely reduced. Thus, we obtained high-quality full-length transcriptome sequencing data for four coral holobionts that is suitable for subsequent analysis.

**Table 2.**
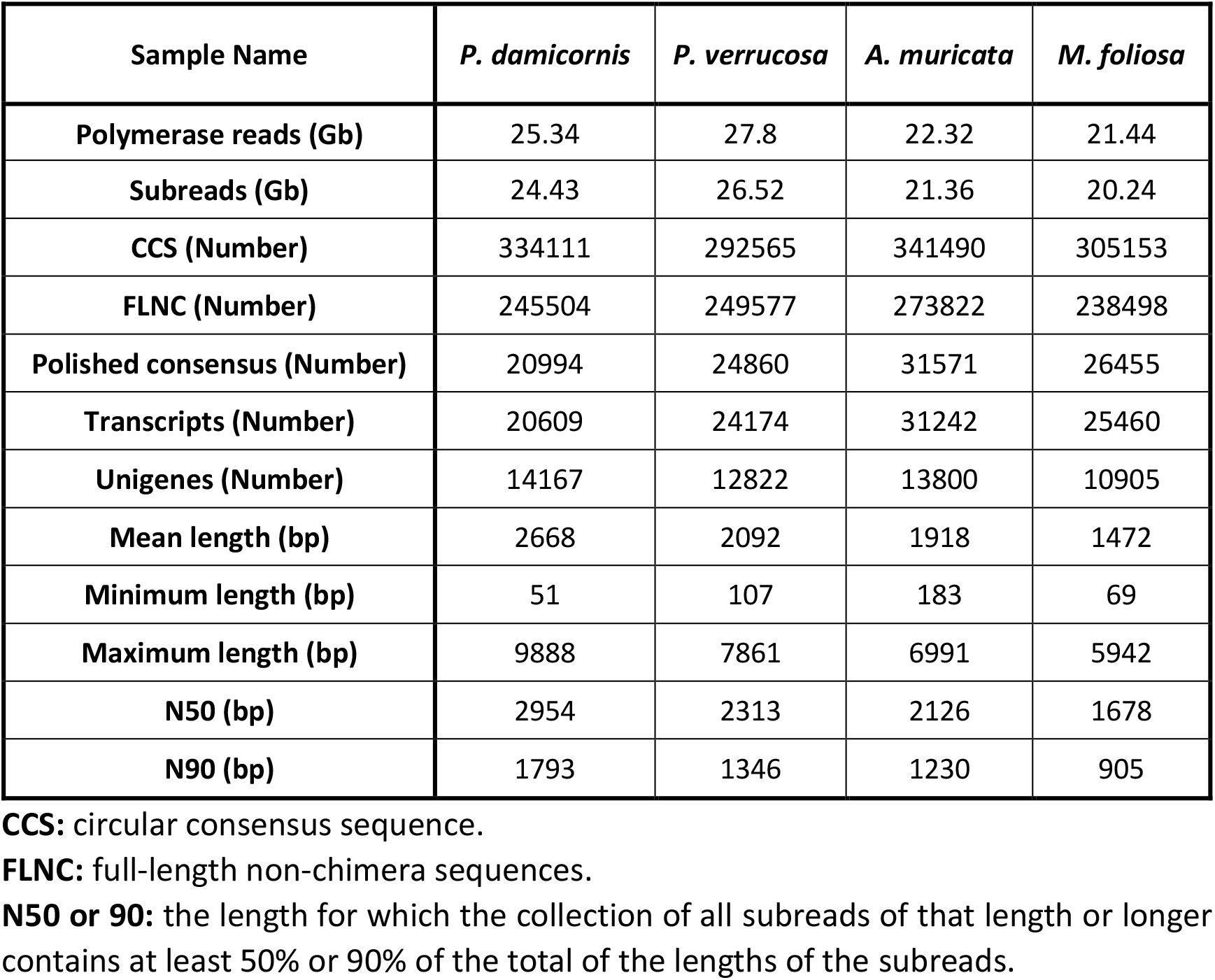
Statistics of PacBio Iso-seq data.

### Gene function annotation and structure analysis

To obtain comprehensive gene function information, seven authoritative databases (NR [121], NT, Pfam [122], KOG [123], Swiss-Prot [124], KEGG [125], and GO [126]) were utilized to annotate the unigenes, and related statistics are shown in Fig. 1 (Additional file 4: Table S3; Additional file 1: Fig. S3-11). In the four corals, 93.88% (*P. damicornis*), 89.32% (*P. verrucosa*), 95.10% (*A. muricata*) and 79.64% (*M. foliosa*) of the unigenes were annotated in at least one database (Fig. 1a; Additional file 4: Table S3.1). The percentage of annotated unigenes in *A. muricata*, *P. damicornis* and *P. verrucosa* was approximately 90%, suggesting that most of the unigenes in these species are orthologues of genes with functional annotations available. In contrast, the percentage of unigenes annotated in *M. foliosa* was only 79.64% because sequencing data from *Montipora* are rare, and the adaptive specialized evolution of these species means that their genomes show especially large differences from those of other reef-building corals with available omics data [92, 127], limiting their in-depth annotation. It was also obvious that the percentage of annotated unigenes in *A. muricata* was higher than that in the other three corals (Fig. 1b) because *A. digitifera* [128] and *A. millepora* [129], with reported genome data, both belong to the same genus as *A. muricata*, providing a good reference genome background.

**Fig. 1.**
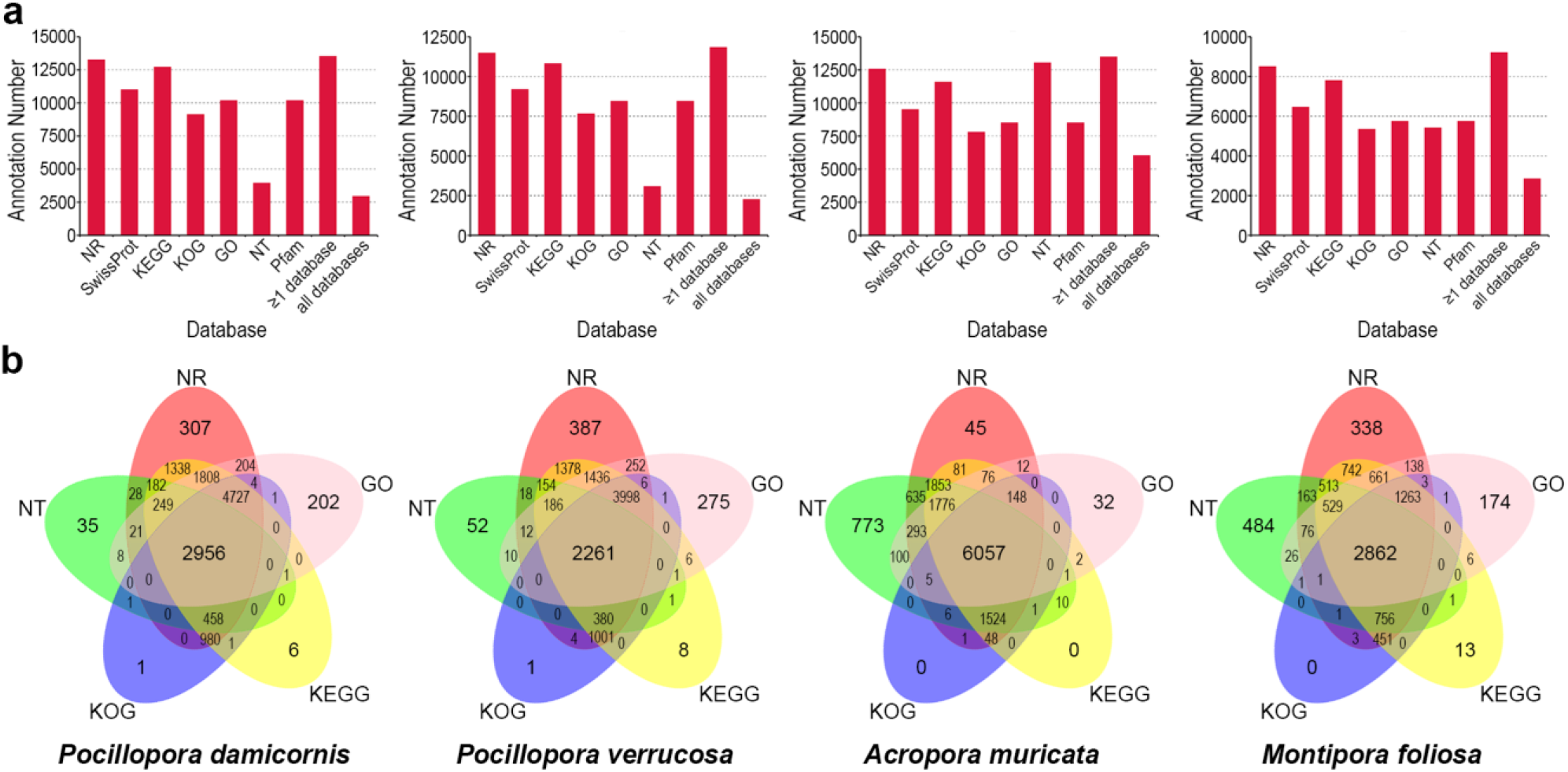
Summary of coral gene functional annotation. **a.** Statistics of the annotation results of four reef-building corals, excluding Symbiodiniaceae sequences, in the NR, Swiss-Prot, KEGG, KOG, GO NT and Pfam databases. The horizontal axis represents the different functional databases, and the vertical axis represents the number of sequences annotated in different functional databases, at least one database and all databases. **b.** Venn plots of the number of annotated sequences of four reef-building corals excluding Symbiodiniaceae sequences obtained using the NR, KEGG, KOG, GO and NT databases. The sum of the numbers in each large circle represents the number of transcripts annotated in one database, and the overlapping circles indicate the numbers of transcripts annotated to these databases simultaneously.

To explore whether the sequence similarity of corals to other species is consistent with their phylogenetic position, we counted the number of sequences annotated based on species information and found that the top five species with the most highly similar genes to those of the four investigated corals were *A. digitifera*, *Exaiptasia pallida*, *Nematostella vectensis*, *B. belcheri* and *Stylophora pistillata* (or *A. millepora*) (Fig. 2; Table 3; Additional file 4: Table S3.6). These species (other than the amphioxus *B. belcheri*) are all cnidarians, and the similarities of the investigated corals to the reference species were all greater than 93%, indicating relatively high accuracy of the annotations. Interestingly, the NR results indicate that the gene expression profiles of all four investigated corals were close to that of *B. belcheri* (Fig. 2; Table 3; Additional file 4: Table S3.6), implying that the presence of intracellular algal cells in both reef-building corals and amphioxus [57] probably impacts the component pattern of the gene expression profiles of the host, leading to a convergence of adaptive evolution.

**Fig. 2.**
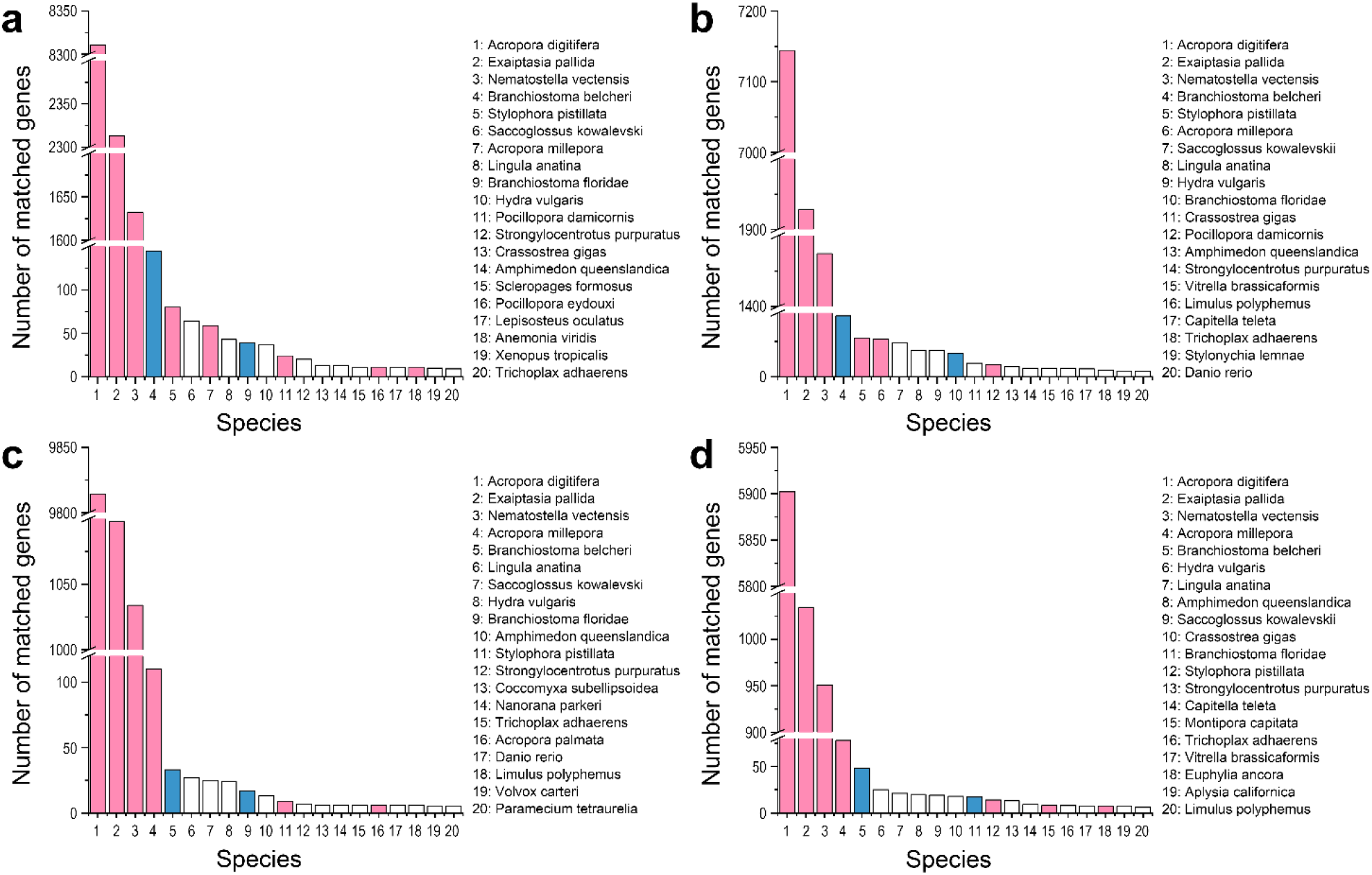
NR database annotation of corals. The top 20 species with the greatest number of top sequence hits to *P. damicornis* (**a**), *P. verrucosa* (**b**), *A. muricata* (**c**) and *M. foliosa* (**d**) are shown. The horizontal axis represents the species ID, and the vertical axis represents the number of coral unigenes annotated to different species. The pink columns represent species belonging to the Actinozoa. The blue columns represent species belonging to the genus *Branchiostoma*.

**Table 3.**
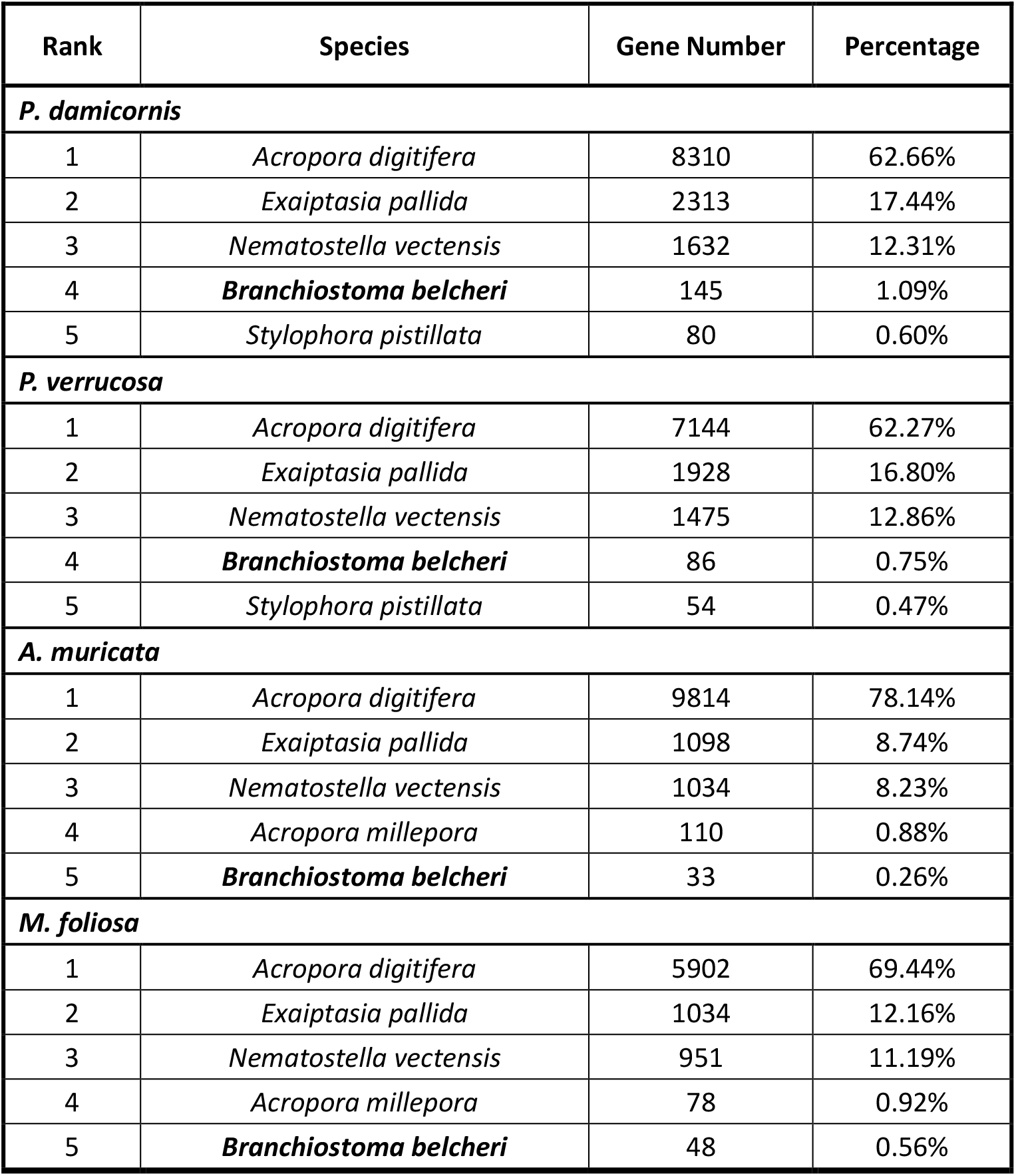
The top five species with the most similar genes to the four investigated corals.

| Rank | Species | Gene Number | Percentage |
| --- | --- | --- | --- |
| ***P. damicornis*** |  |  |  |
| 1 | *Acropora digitifera* | 8310 | 62.66% |
| 2 | *Exaiptasia pallida* | 2313 | 17.44% |
| 3 | *Nematostella vectensis* | 1632 | 12.31% |
| 4 | ***Branchiostoma belcheri*** | 145 | 1.09% |
| 5 | *Stylophora pistillata* | 80 | 0.60% |
| ***P. verrucosa*** |  |  |  |
| 1 | *Acropora digitifera* | 7144 | 62.27% |
| 2 | *Exaiptasia pallida* | 1928 | 16.80% |
| 3 | *Nematostella vectensis* | 1475 | 12.86% |
| 4 | ***Branchiostoma belcheri*** | 86 | 0.75% |
| 5 | *Stylophora pistillata* | 54 | 0.47% |
| ***A. muricata*** |  |  |  |
| 1 | *Acropora digitifera* | 9814 | 78.14% |
| 2 | *Exaiptasia pallida* | 1098 | 8.74% |
| 3 | *Nematostella vectensis* | 1034 | 8.23% |
| 4 | *Acropora millepora* | 110 | 0.88% |
| 5 | ***Branchiostoma belcheri*** | 33 | 0.26% |
| ***M. foliosa*** |  |  |  |
| 1 | *Acropora digitifera* | 5902 | 69.44% |
| 2 | *Exaiptasia pallida* | 1034 | 12.16% |
| 3 | *Nematostella vectensis* | 951 | 11.19% |
| 4 | *Acropora millepora* | 78 | 0.92% |
| 5 | ***Branchiostoma belcheri*** | 48 | 0.56% |

The prediction of CDSs is helpful for potential unigene analysis and provides a basis for subsequent protein analysis. The CDS lengths of the four corals were concentrated in the range of 300–1700 nt (Methods; Additional file 1: Fig. S12a), and a total of 97.91% (*P. damicornis*), 95.67% (*P. verrucosa*), 95.51% (*A. muricata*) and 89.04% (*M. foliosa*) of the unigenes were predicted to contain CDS regions (Additional file 5: Table S4.1). We then identified coral TFs among the predicted protein CDS results based on Pfam TF families (Methods) and found that the five largest TF families in the corals were ZBTB, zf-C2H2, homeobox, bHLH and TF_bZIP (Additional file 1: Fig. S12b; Additional file 5: Table S4.2). The results of simple sequence repeat (SSR) detection showed that most SSR sequences corresponded to poly(A) tails (Additional file 1: Fig. S12c; Additional file 5: Table S4.3), reconfirming the absence of genomic DNA contamination. Transcripts without coding potential were classified as lncRNAs. Generally, lncRNAs are shorter than mRNAs (Additional file 1: Fig. S12e). The number of predicted lncRNAs in *M. foliosa* was much greater than those in the other corals (Additional file 1: Fig. S12d). After CDS, TF, SSR and lncRNA analyses, we quantified the gene expression profiles in each coral by mapping the Illumina sequencing reads to the full-length transcriptome data. This mapping directly providing read count values that could be converted to the expected number of fragments per kilobase of transcript sequence per million base pairs sequenced (FPKM) or transcripts per kilobase million (TPM) values for further ensuring diverse quantitative analysis under different conditions (Additional file 6: Table S5). Based on these data, the Pearson correlation coefficient (R) analysis can be used to reflect the gene expression profile similarity among the samples and thus verify the experimental reliability and the rationality of sample selection.

### Gene expression profile analysis

We next compared the gene expression profiles of the four corals to better understand their ecological and evolutionary relationships. Consistent with the known phylogenetic relationships [130], the gene expression profiles of *P. damicornis* and *P. verrucosa* (both in the Robusta clade) were more similar to each other than to *A. muricata* and *M. foliosa* (both in the Complexa clade), and vice versa (Fig. 3a-d). The gene expression levels of each coral were considered to be consistent across all replicates (Additional file 1: Fig. S13). However, combined analysis indicated that the overall gene expression level of *M. foliosa* was lower than those of the other corals (Fig. 3c), showing that the structure of the gene expression profile of *M. foliosa* was significantly different from those of the other corals (Additional file 6: Table S5). These results indicated that beyond evolutionary aspects, *M. foliosa* exhibits prominent ecological adaptive features [92, 127]. In addition, a Pearson’s R^2^> 0.92 among the three biological replicates of each coral suggested that the sample selection was reasonable and the experimental data were reliable (Fig. 3d). Then, the differences in the coral gene expression profiles were explored. A total of 871 (*P. damicornis*), 1296 (*P. verrucosa*), 4369 (*A. muricata*) and 3911 (*M. foliosa*) differentially expressed genes (DEGs) were identified based on the parameters padj<0.001&|log2(FoldChange)|≥2 (Additional file 1: Fig. S14a; Additional file 7: Table S6); of these, 122 (*P. damicornis*), 167 (*P. verrucosa*), 2333 (*A. muricata*) and 1956 (*M. foliosa*) were particularly prominent, with a |log2(FoldChange)|≥10 (Additional file 1: Fig. S14b; Additional file 7: Table S6). These results showed that the genus *Pocillopora*, of the Robusta clade, presented fewer differentially expressed genes than the genera of the *Acropora* and *Montipora* Complexa clades (Additional file 1: Fig. S14; Additional file 7: Table S6). Consequently, according to the gene expression profile results, biomineralization, symbiosis, and circadian gene families, which provide crucial information on coral population structure and on the evolution of coral gene repertoires, will be highlighted hereafter.

**Fig. 3.**
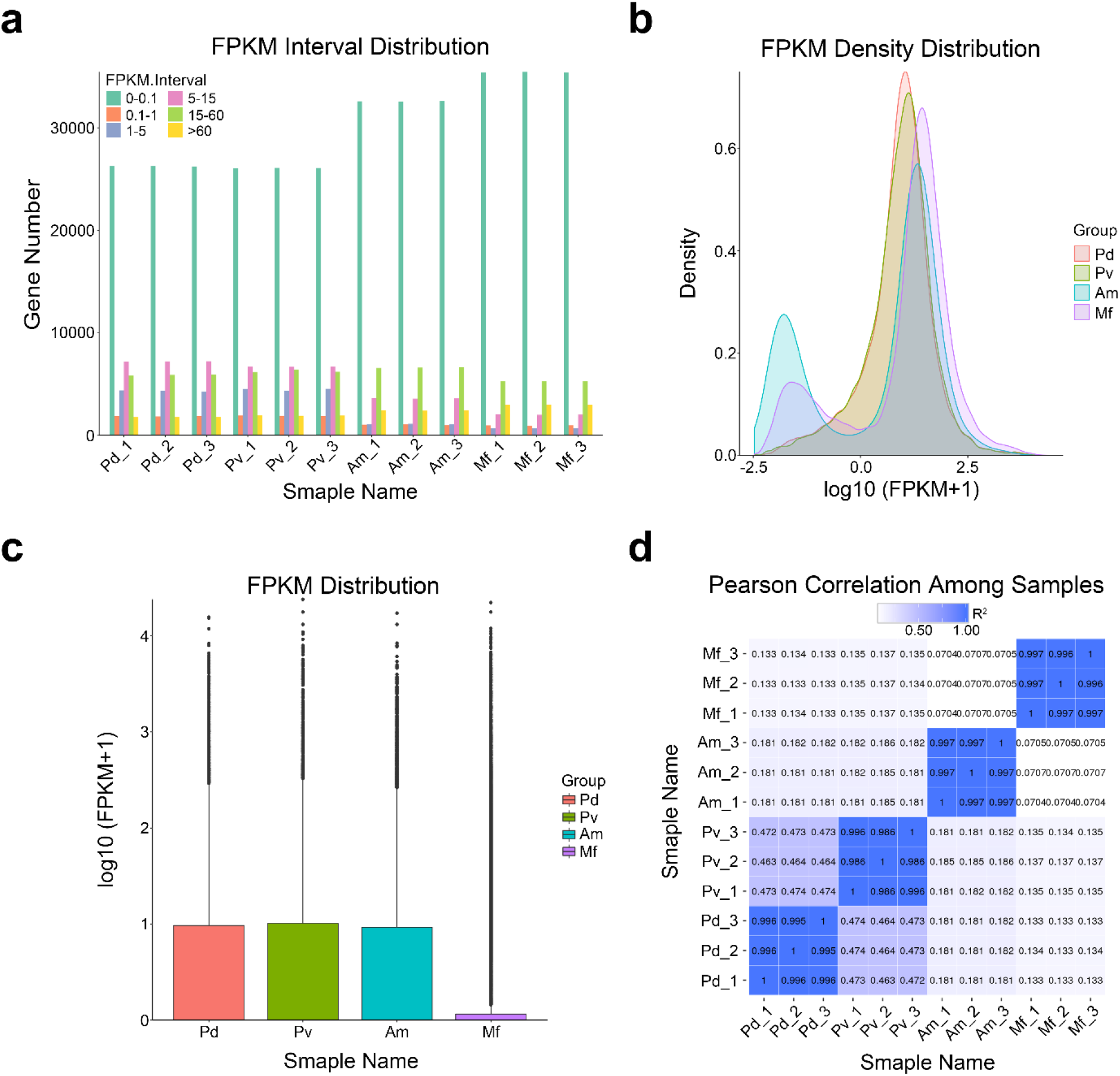
Summary of the coral gene expression level analysis. **a**. FPKM interval distribution. The horizontal axis shows the sample name, different colours represent FPKM intervals, and the vertical axis represents the number of genes in each interval. **b**. FPKM density distribution. The horizontal axis represents the log10 (FPKM+1) values, and the vertical axis represents the density of genes with different expression levels. **c**. FPKM box plot. The horizontal axis shows the sample name, and the vertical axis represents the log10 (FPKM+1) values. Each box plot shows five statistical values, including the maximum, upper quartile, median, lower quartile and minimum, from top to bottom. **d**. Pearson correlation among samples. The closer the value is to 1, the better the correlation. Pd: *P. damicornis*; Pv: *P. verrucosa*; Am: *A. muricata*; Mf: *M. foliosa*.

### Expression analysis of the biomineralization-related gene group

Although the biomineralization mechanisms of reef-building corals are similar (Fig. 4a), they are displayed in a wide range of morphological differences. To gain insight into the molecular-level causes of this phenomenon, we investigated the expression patterns and features of biomineralization-related gene families and found that the expression levels of calcium ATPases and bicarbonate transporters, which are essential for Ca^2+^ and HCO_3_^−^ transport, were higher in the Robusta genus *Pocillopora* than in the Complexa genera *Acropora* and *Montipora* (Fig. 4b-c; Table 4; Additional file 8: Table S7). Among the four investigated corals, *M. foliosa* has the lowest apparent capacity to transport calcium ions to calcifying fluid because of the low expression of calcium ATPases (Fig. 4b; Table 4; Additional file 8: Table S7). CA2 is universally expressed in all four corals at very high levels and accounts for a major proportion of the CA content, implying its key role in coral biomineralization (Fig. 4d; Table 4; Additional file 8: Table S7).

**Fig. 4.**
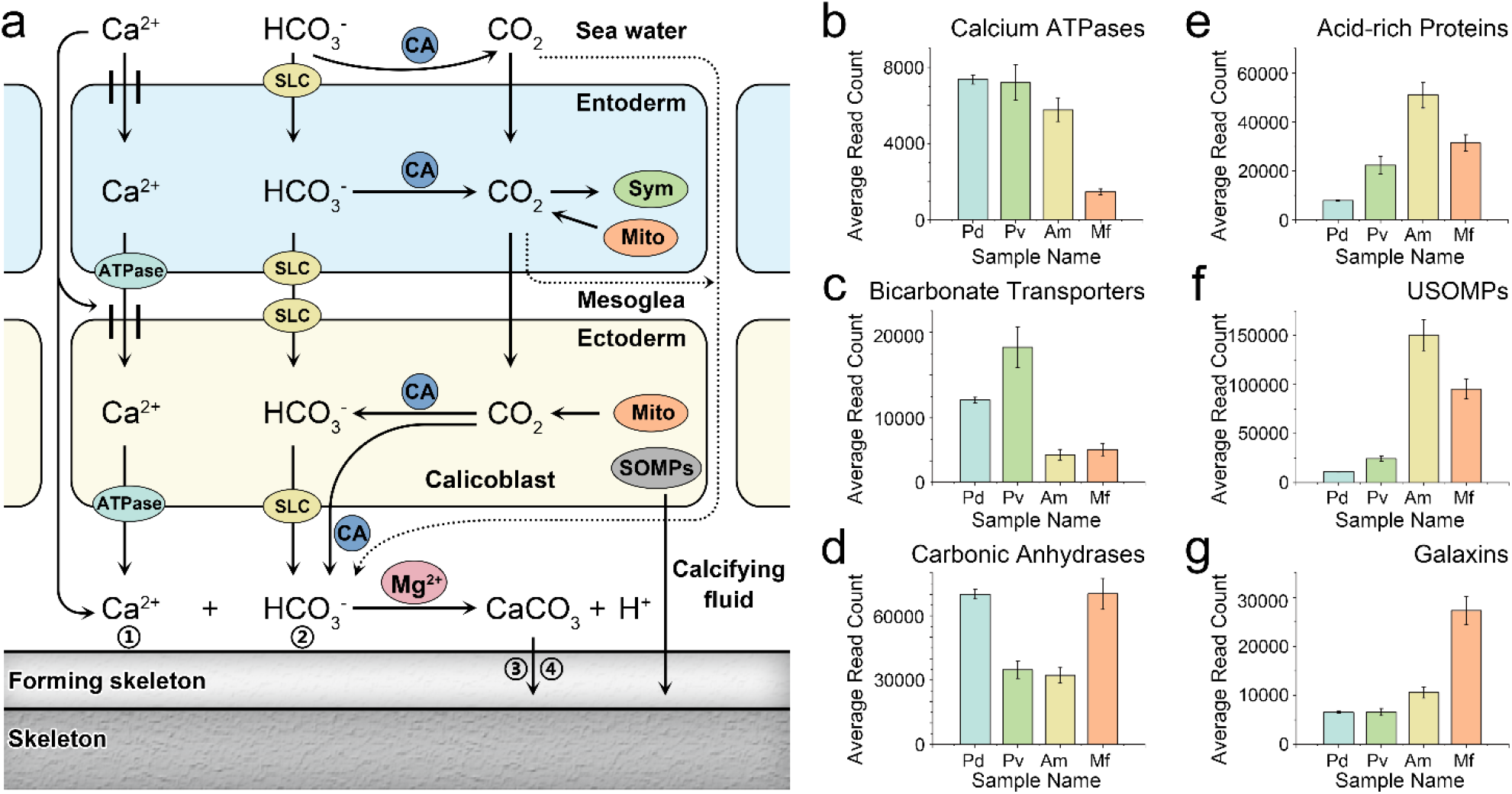
Mechanism of coral biomineralization and related gene expression levels. **a**. The mechanism of reef-building coral biomineralization. (1) Ca^2+^ transport by calcium ATPase or diffusion. (2) CO_2_ can be converted into HCO_3_^−^ by CA and then exits the cells via bicarbonate transporters (i.e., SLC4 and SLC26). (3) Coral acid-rich proteins and Mg^2+^ can promote crystal nucleation, determine growth axes and control crystal growth. (4) The bioprecipitation of aragonite crystals in corals requires additional skeletal organic matrix proteins (USOMPs, galaxins and alpha IV collagen). The average expression levels of genes belonging to calcium ATPases (**b**), bicarbonate transporters (**c**), carbonic anhydrases (**d**), acid-rich proteins(**e**), USOMPs (**f**), and galaxins (**g**). The horizontal axis shows the sample name, and the vertical axis represents the average read count values of each set of genes. ATPase: calcium ATPase; SLC: solute carrier 4 (SLC4) and solute carrier 26 (SLC26); CA: carbonic anhydrase; Sym: Symbiodiniaceae; Mito: mitochondrion; and SOMPs: coral acid-rich proteins, alpha IV collagen, galaxins and uncharacterized skeletal organic matrix proteins (USOMPs). The solid lines represent definite paths, and the dashed lines represent possible paths. Pd: *P. damicornis*; Pv: *P. verrucosa*; Am: *A. muricata*; and Mf: *M. foliosa*.

**Table 4.** Average read count values of key biomineralization-related genes in four investigated corals.

| Class |  | Protein Name | *P.damicornis* | *P. verrucosa* | *A. muricata* | *M. foliosa* |
| --- | --- | --- | --- | --- | --- | --- |
| Calcium ATPase |  | Plasma membrane calcium-transporting ATPase 1 | 0 | 0 | 3640.4 | 28 |
|  |  | Plasma membrane calcium-transporting ATPase 2 | 1574.217 | 685.81 | 6.333333 | 1029.333 |
|  |  | Plasma membrane calcium-transporting ATPase 3 | 1.783333 | 503.5267 | 785.6667 | 409.3333 |
|  |  | Plasma membrane calcium-transporting ATPase 4 | 1502 | 1900.66 | 0 | 0 |
|  |  | Plasma membrane calcium ATPase | 4291.337 | 4124.67 | 0.666667 | 0 |
|  |  | Calcium-transporting ATPase type 2C member 1 | 0 | 0 | 1336.633 | 0 |
| Bicarbonate Transporter |  | solute carrier 4 (SLC4) | 9643.527 | 8724.997 | 5110.58 | 5264 |
|  |  | solute carrier 26 (SLC26) | 2450.687 | 9511.6 | 602.3333 | 1035 |
| Alpha Carbonic Anhydrase |  | carbonic anhydrase 1 | 741.3333 | 216 | 0 | 0 |
|  |  | carbonic anhydrase 2 | 68885.42 | 34358 | 29819.85 | 63346.04 |
|  |  | carbonic anhydrase 3 | 439 | 206.6667 | 0 | 0 |
|  |  | carbonic anhydrase 12 | 0 | 0 | 2448.153 | 6891.21 |
| Acidic Proteins |  | Skeletal aspartic acid-rich protein 1 (SAARP1) | 6040.233 | 20674.29 | 11277.34 | 2966.333 |
|  |  | Skeletal aspartic acid-rich protein 2 (SAARP2) | 165.6667 | 297.3333 | 948 | 0 |
|  |  | Acidic skeletal organic matrix protein (Acidic SOMP) | 1715.333 | 1351.663 | 5737.333 | 4792.333 |
|  |  | Secreted acidic protein 1 (SAP1) | 0 | 0 | 0.333333 | 4101.47 |
|  |  | Secreted acidic protein 2 (SAP2) | 0 | 0 | 12490.67 | 7805.12 |
|  |  | Aspartic and glutamic acid-rich protein | 0 | 0 | 20481.67 | 11682.22 |
| Unique Uncharacterized Proteins | Uncharacterized skeletal organic matrix protein | Uncharacterized skeletal organic matrix protein-1 (USOMP-1) | 0 | 0 | 522.3333 | 0 |
|  |  | Uncharacterized skeletal organic matrix protein-2 (USOMP-2) | 227.6633 | 120.33 | 1067.333 | 4337.667 |
|  |  | Uncharacterized skeletal organic matrix protein-3 (USOMP-3) | 2042.107 | 2688.417 | 1664.333 | 3863.997 |
|  |  | Uncharacterized skeletal organic matrix protein-4 (USOMP-4) | 0 | 0 | 20889.33 | 0.666667 |
|  |  | Uncharacterized skeletal organic matrix protein-5 (USOMP-5) | 7567.913 | 18213.03 | 6069.08 | 2209.667 |
|  |  | Uncharacterized skeletal organic matrix protein-6 (USOMP-6) | 0 | 0 | 118232.9 | 80799.58 |
|  |  | Uncharacterized skeletal organic matrix protein-7 (USOMP-7) | 948.47 | 199.6667 | 2047 | 4131.667 |
|  |  | Uncharacterized skeletal organic matrix protein-8 (USOMP-8) | 168.6667 | 3082 | 0 | 0 |
|  | Galaxin | Galaxin | 6541 | 6595.007 | 5360.667 | 22081.34 |
|  |  | Galaxin 2 | 0 | 0 | 5228.327 | 5230.173 |
| Collagen alpha-6(VI) |  | Collagen alpha-6(VI) chain | 226 | 222.6667 | 3242.003 | 447.3333 |

The expression levels of coral biomineralization-related extracellular matrix (ECM) proteins, including acid-rich proteins, USOMPs, galaxins and collagen alpha-6(VI) chain proteins, were all obviously higher in the Complexa clade than in the Robusta clade (Fig. 4e-g; Table 4; Additional file 8: Table S7). Regarding key acid-rich ECM proteins, *A. muricata* showed the highest expression levels, diversity and balance, followed by *M. foliosa*, while skeletal aspartic acid-rich protein 1 (SAARP1) was the only protein in this group expressed in genus *Pocillopora*. Regarding the ECM accessory proteins USOMPs, *A. muricata* and *M. foliosa* presented extremely higher levels and diversity, and USOMP-6 was the dominant *USOMP* gene, whereas members of the genus *Pocillopora* mainly expressed USOMP-5 (Table 4; Additional file 8: Table S7). Galaxins were the only ECM components with higher expression in *M. foliosa* than in *A. muricata* (Table 4; Additional file 8: Table S7). This result indicates that there are significant differences in the types and expression levels of core calcification genes among different reef-building corals, which may affect their morphological formation and growth model.

### Genes expressed by Symbiodiniaceae (intracellular symbionts of coral) are mainly involved in energy and nutrient production

The secretion of calcium carbonate skeletons to form marine reefs and the reliance on intracellular symbiotic algae (Symbiodiniaceae) for nutrient acquisition are two signature features of reef-building coral physiology [14–21, 128]. Although skeletal formation has been discussed extensively, the contributions of the genes expressed by the Symbiodiniaceae are rarely mentioned. Depending on identifying coral or symbiont transcripts based on alignment to previously published sequences, we found that 235 (*P. damicornis*), 351 (*P. verrucosa*), 220 (*A. muricata*) and 561 (*M. foliosa*) Symbiodiniaceae unigenes were expressed in the investigated coral holobionts (Additional file 9: Table S8). Similar to the analysis process used for corals, Symbiodiniaceae gene expression profiling was performed using seven functional databases, NR, NT, Pfam, KOG, Swiss-Prot, KEGG, and GO, and published Symbiodiniaceae genome data (Additional file 1: Fig. S15-26; Additional file 9: Table S8.5-9). We found 832 transcripts that were expressed by the symbiotic Symbiodiniaceae of all four investigated corals (Additional file 1: Fig. S27), accounting for approximately 70.75% (*P. damicornis*), 70.87% (*P. verrucosa*), 88.79% (*A. muricata*) and 93.69% (*M. foliosa*) of the symbiote transcripts. Overall, gene expression profiles of the Symbiodiniaceae were more similar than those of coral hosts, suggesting considerable gene expression convergence. On the other hand, the gene expression patterns of the symbionts of *P. damicornis* and *P. verrucosa* were more similar to each other than to *A. muricata* and *M. foliosa*, and vice versa, coinciding with the evolutionary relationships of the corals (Additional file 1: Fig. S28).

To explore the functions of these expressed genes, we classified them according to the annotation results using the GO database (Fig. 5; Additional file 1: Fig. S29; Additional file 9: Table S8.11-12). In the biological process category, the identified gene functions were mainly involved in oxidation reduction and carbohydrate metabolism (Fig. 5). In the cellular component category, most of the functional genes were related to photosynthesis, such as light-harvesting complex or photosystem (Additional file 1: Fig. S29). In the molecular function category, the largest group of genes was associated with oxidoreductase or carbonate dehydratase activity (Additional file 1: Fig. S29). These results suggest that the symbiotes of all four corals have similar expression profiles that feature genes related to energy and nutrient production.

**Fig. 5.**
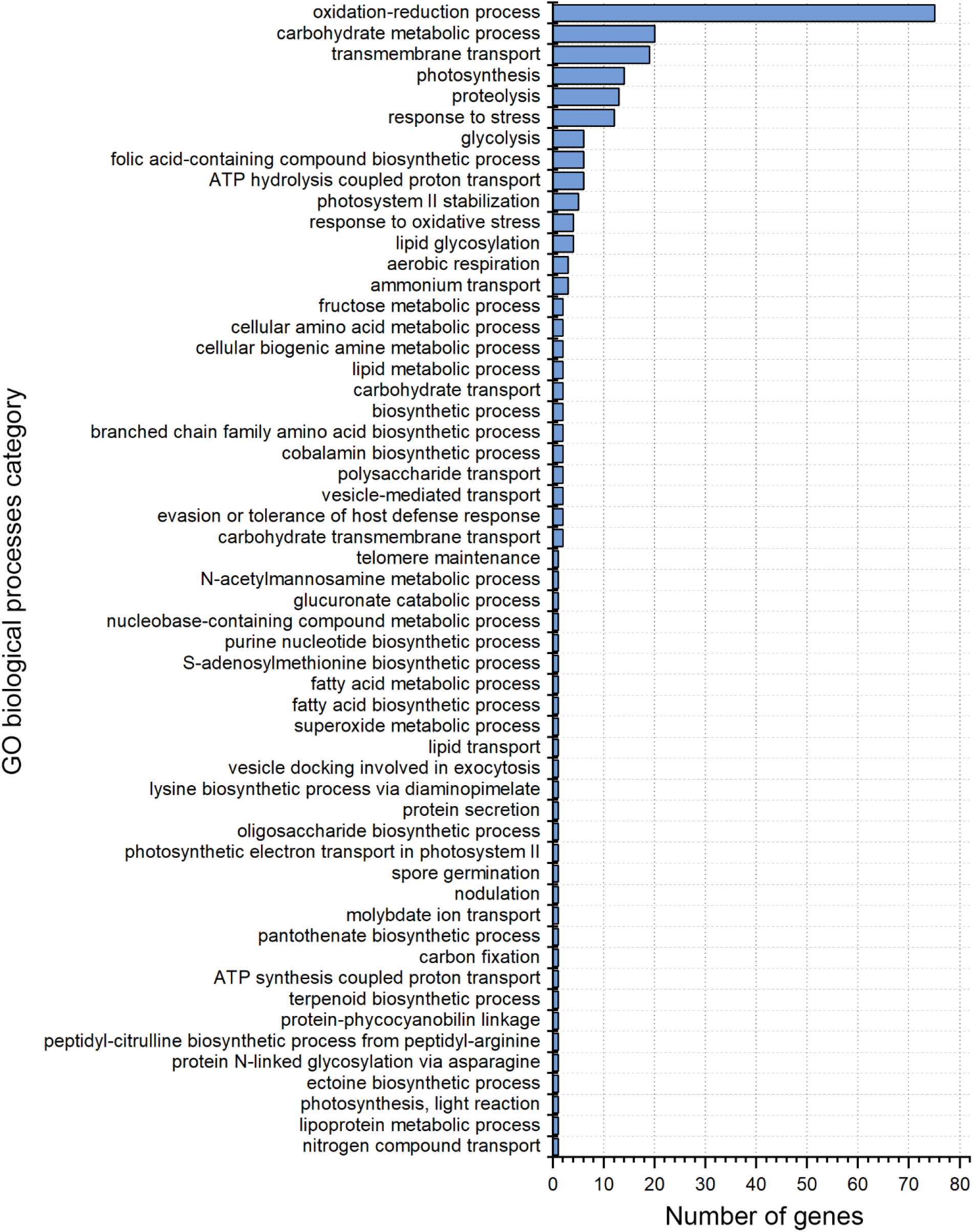
Bar graph of coexpressed Symbiodiniaceae sequences based on GO biological processes categories. The vertical axis represents the GO terms, and the horizontal axis represents the number of transcripts annotated to the terms (including subterms of the terms).

### Phylogenetic analysis of the key circadian clock gene regulation network

The circadian clock system is one of the most universal and fundamental characteristics of life across almost all living species [66]. Corals also exhibit circadian behaviours, and there have been studies describing the molecular mechanisms underlying the regulation of these behaviors [35–37]. However, whether this mechanism is applicable to other corals, other cnidarians and even other animals remains to be clarified. To explore the evolutionary origins of the key members of the circadian clock gene regulation network of reef-building corals, including the four corals studied herein, we selected 27 evolutionarily representative species based on their related gene and genome data available in the NCBI database to investigate the key circadian clock gene regulation network, namely, the *cry1*, *cry2*, *Clock, Npas2*, *cyc, Arntl* (also known as *Bmal1*), *Arntl2* (also known as *Bmal2*), *per1*, *per2*, *per3* and *tim* genes (Table 5; Additional file 10: Table S9), and performed phylogenetic analyses (Fig. 6; Additional file 1: Fig. S30-36). The results showed that Ciliata (*Paramecium tetraurelia*), Ctenophora (*Pleurobrachia bachei*), Scyphozoa (*Aurelia aurita*), Cubozoa (*Chironex fleckeri*) and Turbellaria (*Schmidtea mediterranea*) had no circadian clock gene members (Table 5). Phylogenetic tree analysis illustrated that the key circadian clock gene regulatory network evolved in early metazoans, whereas protozoans and the ctenophore clade did not have this kind of gene regulatory network; moreover, the jellyfish and Turbellaria clades showed the complete loss of this network during evolutionary adaptation (Table 5; Fig. 6; Additional file 1: Fig. S30-36; Additional file 10: Table S9). In the Hydrozoa and Ascidiacea clades, all members of the circadian clock gene regulatory network except for the *tim* gene have been lost. The *tim* gene was the only gene of this group existing in all species with a circadian clock system, suggesting that it is the most conserved and stable circadian clock gene (Table 5; Fig. 6; Additional file 10: Table S9) [92, 130]. Consequently, the phylogenetic analysis suggested that in the early metazoan stage, there were four key gene members of the circadian clock gene regulatory network: *cry*; *Clock* or *Npas2*; *cyc*, *Arntl*(*Bmal1*), or *Arntl2*(*Bmal2*); and *tim*. The *cry1*, *Clock* or *Npas2*, *Arntl (Bmal1)* and *tim* genes were present in all investigated reef-building corals, and *cyc* was present in other corals, as shown in Table 5. Thus, the data suggest the existence of four key conserved members of the reef-building coral circadian clock gene regulation network. The phylogenetic analysis also indicated that the *per* gene is a novel member of the circadian clock gene regulation network in the Bilateria, as it is found in protostomes and deuterostomes but not in early metazoans (Table 5). Interestingly, the phylogenetic analysis results also showed that the two most conserved clades with key circadian clock gene regulation networks are sessile-living anthozoans and free-living vertebrates. Together, these findings indicate that there were four key members of the circadian clock gene regulatory network in early metazoans, while *per*, as the fifth member, first occurred in Bilateria.

**Fig. 6.**
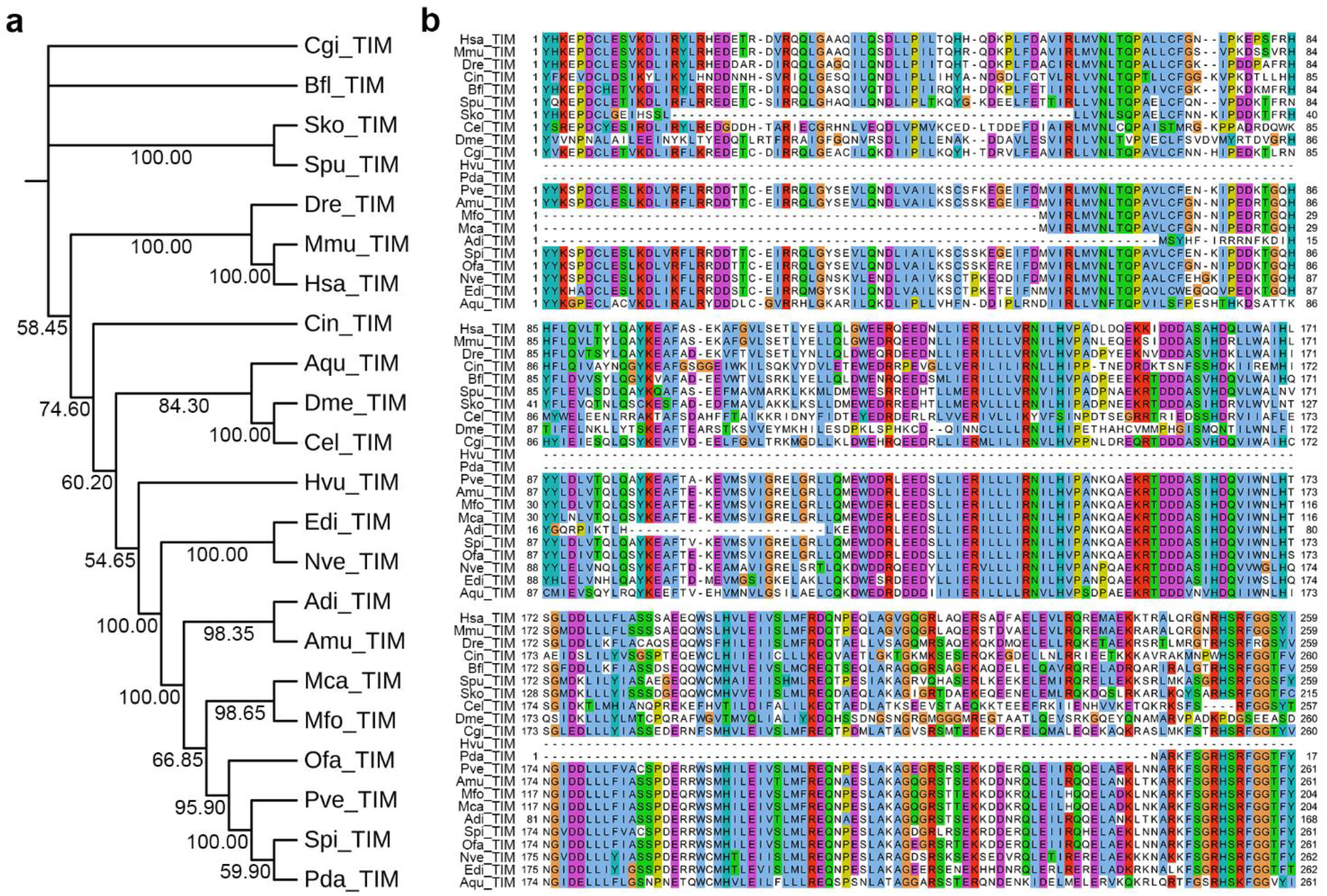
*Tim* gene phylogenetic analysis. **a.** Phylogenetic tree of *tim* genes based on the Neighbor-Joining method. **b.** Conserved domains of *tim* genes among different species. Hsa: *Homo sapiens*; Mmu: *Mus musculus*; Dre: *Danio rerio*; Cin: *Ciona intestinalis*; Bfl: *Branchiostoma floridae*; Spu: *Strongylocentrotus purpuratus*; Sko: *Saccoglossus kowalevskii*; Cel: *Caenorhabditis elegans*; Dme: *Drosophila melanogaster*; Cgi: *Crassostrea gigas*; Hvu: *Hydra vulgaris*; Pda: *Pocillopora damicornis*; Pve: *Pocillopora verrucosa*; Amu: *Acropora muricata*; Mfo: *Montipora foliosa*; Mca: *Montipora capricornis*; Adi: *Acropora digitifera*; Spi: *Stylophora pistillata*; Ofa: *Orbicella faveolata*; Nve: *Nematostella vectensis*; Edi: *Exaiptasia diaphana*; and Aqu: *Amphimedon queenslandica*.

**Table 5.** Presence of key circadian clock genes in 27 species.

| Class | Represent Species | Key Circadian Clock Genes |  |  |  |  |  |  |  |  |  |  |
| --- | --- | --- | --- | --- | --- | --- | --- | --- | --- | --- | --- | --- |
|  |  | *cry1* | *cry2* | *Clock* | *Npas2* | *cyc* | *Arntl* (*Bmal1*) | *Arntl2* (*Bmal2*) | *per1* | *per2* | *per3* | *tim* |
| Mammalia | *Homo sapiens* | Y | Y | Y | Y | - | Y | Y | Y | Y | Y | Y |
|  | *Mus musculus* | Y | Y | Y | Y | - | Y | Y | Y | Y | Y | Y |
| Actinopterygii | *Danio rerio* | Y | Y | Y | Y | - | Y | Y | Y | Y | Y | Y |
| Ascidiacea | *Ciona intestinalis* | - | - | - | - | - | - | - | - | - | - | Y |
| Cephalochordata | *Branchiostoma floridae* | Y | - | Y | Y | - | Y | - | Y | - | - | Y |
| Echinoidea | *Strongylocentrotus purpuratus* | Y | Y | Y | - | Y | - | - | - | - | - | Y |
| Enteropneusta | *Saccoglossus kowalevskii* | - | - | - | - | - | Y | - | - | - | - | Y |
| Nematoda | *Caenorhabditis elegans* | - | - | Y | Y | Y | Y | Y | Y | - | - | Y |
| Insecta | *Drosophila melanogaster* | Y | Y | Y | Y | Y | Y | Y | Y | - | - | Y |
| Bivalvia | *Crassostrea gigas* | Y | Y | Y | - | Y | Y | - | Y | Y | - | Y |
| Turbellaria | *Schmidtea mediterranea* | - | - | - | - | - | - | - | - | - | - | - |
| Actinozoa | *Pocillopora damicornis* | Y | - | Y | Y | Y | Y | - | - | - | - | Y |
|  | *Pocillopora verrucosa* | Y | - | Y | Y | Y | Y | - | - | - | - | Y |
|  | *Acropora muricata* | Y | - | Y | Y | Y | Y | - | - | - | - | Y |
|  | *Montipora foliosa* | Y | - | Y | Y | - | Y | - | - | - | - | Y |
|  | *Montipora capricornis* | Y | - | Y | Y | - | Y | - | - | - | - | Y |
|  | *Acropora digitifera* | Y | - | Y | Y | Y | Y | - | - | - | - | Y |
|  | *Stylophora pistillata* | Y | - | Y | Y | Y | Y | - | - | - | - | Y |
|  | *Orbicella faveolata* | Y | - | Y | Y | Y | Y | - | - | - | - | Y |
|  | *Nematostella vectensis* | Y | - | Y | - | Y | Y | - | - | - | - | Y |
|  | *Exaiptasia diaphana* | Y | - | Y | Y | - | Y | - | - | - | - | Y |
| Hydrozoa | *Hydra vulgaris* | - | - | - | - | - | - | - | - | - | - | Y |
| Cubozoa | *Chironex fleckeri* | - | - | - | - | - | - | - | - | - | - | - |
| Scyphozoa | *Aurelia aurita* | - | - | - | - | - | - | - | - | - | - | - |
| Sponges | *Amphimedon queenslandica* | - | Y | Y | - | Y | Y | Y | - | - | - | Y |
| Ctenophore | *Pleurobrachia bachei* | - | - | - | - | - | - | - | - | - | - | - |
| Ciliata | *Paramecium tetraurelia* | - | - | - | - | - | - | - | - | - | - | - |

'Y' means that the genome of the indicated species contains the gene, while '-' means that the genome of the indicated species does not contain the gene.

## Discussion

Despite much research, our understanding of reef-building coral holobiont transcriptomes remains incomplete, and the short-read lengths intrinsic to the prevailing technologies have limited access to complete genetic information. Here, we sequenced and analysed the full-length transcriptome of four common dominant reef-building coral holobionts with respect to gene functions, structures, and expression. The results revealed differences in the members actually involved in biomineralization processes in reef-building corals and provided new insights into further understanding the molecular mechanisms of coral density. In particular, we isolated and demarcated the gene expression profiles of the symbiote Symbiodiniaceae, which showed higher convergence than the coral hosts and suggested that there may be some association between marine animal intracellular symbiosis and the intracellular digestion of algal cells. Moreover, we confirmed that there were four key members of the circadian clock gene regulatory network in early metazoans and that *per* first occurred in Bilateria, providing a molecular basis for the study of the evolutionary origin of the animal circadian clock system.

### Coral biomineralization and skeleton density

Coral biomineralization has been studied intensively in previous studies, but descriptions of its formation mechanism vary; here, we summarized this process, which consists of four components (Fig. 4a). (1) Calcium ion transport. In calicoblasts and paracells, calcium ions enter the cells through calcium channels and exit via calcium ATPases that exchange two calcium ions for four protons across the cell membrane [39, 40]. Ca^2+^ diffusion among cells with chemical gradients may also participate in calcium deposition [15]. (2) An HCO_3_^−^ source. The source of HCO_3_^−^ is metabolic and environmental CO_2_. Metabolic CO_2_ can be converted into HCO_3_^−^ both intracellularly and extracellularly, which is catalysed by CA under a favourable pH [41–43], and intracellular HCO_3_^−^ exits cells via bicarbonate transporters belonging to two membrane protein families (SLC4 and SLC26) [44]. In calcifying fluid, the HCO_3_^−^ concentration is higher than the Ca^2+^ concentration, and the amount of calcium carbonate deposition is determined by the Ca^2+^ concentration [43]. (3) Acid-rich proteins. Coral acid-rich proteins are key ECM proteins involved in biomineralization and can interact with amorphous calcium carbonate (ACC) directly, promoting crystal nucleation, determining growth axes and controlling crystal growth [45, 46]. (4) Other organic matrix proteins of the ECM. The bioprecipitation of aragonite crystals in corals also requires additional skeletal organic matrix proteins as binders [47], including USOMPs, galaxins and alpha IV collagen [38, 48, 49]. In addition, magnesium plays a crucial role in regulating the formation of different calcium carbonate phases that cooperate with organic matrix molecules to stabilize ACCs [131].

Here, the above core biomineralization-related gene expression level analyses among four reef-building corals indicate that the Robusta genus *Pocillopora* has a greater capacity to produce calcium carbonate than the Complexa genera *Acropora* and *Montipora*, but the opposite situation was observed for biomineralization-related ECM proteins, suggesting that the species of the Complexa clade can produce a larger volume of skeleton than those of the Robusta clade using the same amount of calcium carbonate, which is in accordance with the reported skeletal density data of these stony corals [34, 132]. Combined with the observation of growth patterns [132], it can be inferred that after the Tertiary Period, the Complexa clade corals developed to occupy more space (as observed *A. muricate* and *M. foliosa*), whereas the Robusta clade corals tended to form more dense colonies (as observed in *P. verrucosa*).

### Evolutionary convergence triggered by intracellular algal cells

Coincidently, all of the gene expression profiling analyses of the four investigated corals indicated evolutionary convergence between corals and amphioxus. According to the NR results, the best-hit genes between the four corals and *B. belcheri* all belonged to the top five groups identified among annotated species (Fig. 2; Table 3; Additional file 4: Table S3.6). All of these genes except for some housekeeping genes were related to interactions with intracellular algae, namely, mechanisms for algal entry into host cells, for reducing algal toxicity and for reducing immune rejection (Additional file 11: Table S10).

Upon the entry of algae into host cells, the packaging of algae by assembling symbiosomes (in intracellular symbioses) or phagosomes (in intracellular digestion) requires the direct or indirect participation of proteins that play important roles in clathrin-mediated endocytosis and endosome formation (DENN domain-containing protein 1B, muskelin, fibrillin-2, exocyst complex component 2 and huntingtin) [133–137] and genes that are required to regulate the generation of functional cilia (nephrocystin-4, jouberin, intraflagellar transport protein 74, the fuzzy homologue protein, the centrosomal protein of 104 kDa and the cytosolic Fe-S cluster assembly factor NUBP2) [138–144]. Nephrocystin-4 contains a proline-rich region and is involved in the organization of the subapical actin network in multiciliated epithelial cells by recruiting INT to the basal bodies of motile cilia, and it subsequently interacts with actin-modifying proteins [138]. Jouberin is involved in recruiting the Ras-related protein Rab-8A to the basal body of the primary cilium as a component of a complex localized in the transition zone of primary cilia, acting as a barrier that prevents the diffusion of transmembrane proteins between cilia and plasma membranes [139, 140]. Intraflagellar transport protein 74, together with intraflagellar transport protein 81, binds and transports tubulin within cilia and is required for ciliogenesis [141]. The fuzzy homologue protein is a planar cell polarity effector involved in cilium biogenesis and regulates protein and membrane transport to the cilium [142]. The centrosomal protein of 104 kDa is required for elongating cilium at the tip during cilium formation [143]. On the other hand, the cytosolic Fe-S cluster assembly factor NUBP2 negatively regulates cilium structure and formation [144]. The observed evolutionarily convergence of these factors between corals and amphioxus indicates that the cilia make an important contribution to algal cell entry of host cells in intracellular symbiosis as well as intracellular digestion, alongside complex molecular recognition mechanisms.

All four investigated symbiotic Symbiodiniaceae (Additional file 9: Table S8.6-9) expressed genes encoding polyketide synthase family members, including putative polyketide synthase 3, phthiocerol synthesis polyketide synthase type I PpsA, phthiocerol synthesis polyketide synthase type I PpsB and phthiocerol synthesis polyketide synthase type I PpsC, demonstrating that symbiotic Symbiodiniaceae can produce dinotoxins [145]. Among the genes commonly expressed by all four corals and amphioxus, several have prospective roles in reducing algal toxicity. Epoxide hydrolase 4 (epoxide hydratase 4) is a biotransformation enzyme that catalyses the hydrolysis of arene and aliphatic epoxides to produce water-soluble dihydrodiols [146], converting highly toxic chemical groups to less reactive and easily excluded groups. THAP domain-containing protein 4 is involved in the detoxification of reactive nitrogen species (RNS) and reactive oxygen species (ROS) [147–148]. S-formylglutathione hydrolase is a kind of serine hydrolase involved in the detoxification of formaldehyde [149]. Carbamoyl-phosphate synthase plays an important role in removing excess ammonia from the cell by catalysing the reaction of ammonia with carboxy phosphate to form carbamoyl phosphate [150]. Transmembrane protein 181 (TMEM181) is the rate-limiting factor in intoxication because it binds to cytolethal distending toxins (CDTs) secreted by entered intracellular algae [151]. The above functional analysis reveals specific genes that reduce algal toxicity to support eukaryotic intracellular symbiosis or digestion by opposing cell intoxication in corals and other hosts of algal symbiotes.

Intracellular algae act as large antigens in algal host cells. For example, the four reef-building corals investigated herein and published coral gene expression profiles all include gene repertoires for innate immunity (Additional file 4: Table S3.2-5) [54], including host innate immune responses to invading microbes. These genes are also involved in intracellular digestion (as in amphioxus) [57]. Acid sphingomyelinase-like phosphodiesterase 3b and phosphoinositide 3-kinase adapter protein 1 can help algal cells adapt to the intracellular environment by regulating Toll-like receptors, which play a key role in the innate immune system. Acid sphingomyelinase-like phosphodiesterase 3b is a lipid-modulating phosphodiesterase that acts as a negative regulator of Toll-like receptors [152, 153]. Phosphoinositide 3-kinase adapter protein 1 can prevent excessive inflammatory cytokine production by linking Toll-like receptor signalling to phosphoinositide 3-kinase (PI3K) activation [154]. Taken together, our findings confirm that there are some evolutionary convergences between coral and amphioxus that have been triggered by interactions with intracellular algal cells.

### Evolutionary origins of the *per* gene

The oscillations of both the transcript and protein levels of the *per* gene have a period of approximately 24 hours and play a central role in the molecular mechanism of the biological clock driving circadian rhythms. In *Drosophila*, after PER is produced from *per* mRNA, it dimerizes with Timeless (TIM), and the complex enters the nucleus and inhibits the TFs *per* and *tim*, which in turn lowers the levels of PER and TIM [53]. When TIM is not complexed with PER, another protein, double-time, or DBT, phosphorylates PER, targeting it for degradation [58]. In mammals, an analogous transcription-translation negative feedback loop is observed. One of the three PER proteins (PER1, PER2 and PER3) that is translated from the mammalian homologs of drosophila-per dimerizes via its PAS domain with one of two cryptochrome proteins (CRY1 and CRY2) to form a negative element of the clock. This dimer then interacts with the CLOCK (or NPAS2) and ARNTL (or ARNTL2) heterodimer, inhibiting its activity and thereby negatively regulating its own expression [133]. Despite an extensive search, we failed to detect orthologs of the *per* gene in our coral data. This also may be the case for sea anemones and *Hydra* (Table 5; Additional file 10: Table S9). On the other hand, amphioxus [134, 135] and *C. gigas* [136] likely have *per* genes, and our results show that the biological clock is a common feature of a wide range of Metazoa, while *per* gene originates from Bilateria. It is unclear, however, whether the ancient circadian clock system initially operated without the *per* gene, whether the Radiata system lost this locus via deletion or other mutation, or whether unknown proteins may have functions similar to the *per* gene. These hypotheses need to be further explored in future research.

Overall, the full-length transcriptome maps of reef-building coral may serve as references for expression analyses of coral under environmental stressors linked to global change, which alter the normal function of reef-building corals. As long-read sequencing continues to evolve in throughput, accuracy, accessibility and cost efficiency, full-length transcriptomes will be adopted by researchers and provide us with unprecedented views of serious biological problems. However, genome, small RNA, proteome and single-cell data are still lacking to further understand the biology of reef-building corals. Future efforts will be required to construct a coral genome database as a reference for analysing functional genes and their associated regulatory networks at the transcriptomic and proteomic levels and further refine them to specific cellular lineages.

## Conclusion

The sequencing and analysis of the full-length transcriptome maps of dominant reef-building coral holobionts contributed to a more in-depth understanding of coral physiology. Related insights improve our understanding of the evolution of circadian rhythms and of holobiosis, and they provide a foundation for further work to protect or even manipulate coral skeleton production or symbiosis to promote the survival of these important organisms. These results provide support for remodelling reef-building corals through advanced gene toolkits under current global climate change and ecological deterioration.

## Methods

### Ethics

All coral samples were collected and processed in accordance with local laws for invertebrate protection.

### Sample collection

The corals in the study were collected from the Xisha Islands in the South China Sea (latitude 15°40’-17°10’ N, longitude 111°-113° E).

### Coral culture system

The coral samples were cultured in our laboratory coral tank with conditions conforming to their habitat environment. All samples were raised in RedSea^®^ tank (redsea575, Red Sea Aquatics Ltd., London, UK) at 26℃ and 1.025 salinity (Red Sea Aquatics Ltd., London, UK). The physical conditions of the coral culture system were as follows: three coral lamps (AI^®^, Red Sea Aquatics Ltd., London, UK), a protein skimmer (regal250s, Reef Octopus), a water chiller (tk1000, TECO Ltd., Port Louis, Mauritius), two wave devices (VorTech^TM^ MP40, EcoTech Marine Ltd. Bethlehem, PA, USA), and a calcium reactor (Calreact 200, Reef Octopus).

### Total RNA extraction

The three biological replicates samples for each coral were isolated from three healthy branches in the same coral independent colony to ensure that enough high-quality RNA (> 15 µg) could be obtained for a PacBio cDNA library and three Illumina cDNA libraries. All the RNA extraction procedures followed the manufacturer’s instructions. The total RNA was isolated with TRIzol^®^ LS Reagent (Thermo Fisher Scientific, 10296028, Waltham, MA, USA) and treated with DNase I (Thermo Fisher Scientific, 18068015, Waltham, MA, USA). The high-quality mRNA was isolated with a FastTrack MAG Maxi mRNA Isolation Kit (Thermo Fisher Scientific, K1580-02, Waltham, MA, USA). The RNA extraction procedure was performed according to the following instructions: (1) grinded coral samples (kept the samples submerged in liquid nitrogen at all times); (2) when the samples were ground into small pieces, the TRIzol^®^ LS reagent was added, the ratio of sample to reagent was about 1:3; (3) let samples stand and thaw naturally; (4) continue adding TRIzol^®^ LS reagent until the samples were dissolved, and dispensed into 50 mL centrifuge tubes; (5) centrifuged at 4 ℃ and 3000 rpm for 5-15 min; (6) dispensed the supernatant into 50 mL centrifuge tubes; (7) added BCP (Molecular Research Center, BP 151, Cincinnati, OH, USA) to the above centrifuge tubes, the ratio of sample to reagent is about 5:1, shook well and then stood for 10 min; (8) centrifuged at 4 ℃ and 10500 rpm for 15 min; (9) took the supernatant, added an equal volume of Isopropanol (Amresco, 0918-500ML, Radnor, PA, USA) and mixed well, stood them overnight at −20℃; (10) centrifuged at 4 ℃ and 10500 rpm for 30 min, discarded the supernatant; (11) rinsed 2 times with 75% Ice Ethyl alcohol, Pure (Sigma-Aldrich, E7023-500ML, Taufkirchen, München, Germany). Finally, three samples of each coral were extracted in equal amounts (total >10 µg) and mixed for PacBio full-length transcriptome sequencing, the remainders (>1.5 µg per sample) were used for Illumina sequencing.

### Total RNA quality testing

Before establishing the library, the quality of total RNA must be tested. RNA degradation and contamination were monitored on 1% agarose gels electrophoresis; RNA purity (OD260/280 ratio) was checked using the NanoPhotometer^®^ spectrophotometer (IMPLEN, CA, USA); RNA concentration was quantified using Qubit® RNA Assay Kit in Qubit^®^ 2.0 Flurometer (Life Technologies, CA, USA); RNA integrity was assessed using the RNA Nano 6000 Assay Kit of the Agilent Bioanalyzer 2100 system (Agilent Technologies, CA, USA).

### Illumina cDNA library construction and sequencing

A total amount of 1.5 µg RNA per sample was used as input material for the RNA sample preparations. Sequencing libraries were generated using NEBNext^®^ Ultra™ RNA Library Prep Kit (E7530L) for Illumina^®^ (NEB, Ipswich, MA, USA) following the manufacturer’s recommendations and index codes were added to attribute sequences to each sample. Briefly, mRNA was purified from total RNA using poly-T oligo-attached magnetic beads. Fragmentation was carried out using divalent cations under elevated temperature in NEBNext First Strand Synthesis Reaction Buffer (5X). First-strand cDNA was synthesized using random hexamer primer and M-MuLV Reverse Transcriptase (RNase H^-^). Second strand cDNA synthesis was subsequently performed using DNA Polymerase I and RNase H. Remaining overhangs were converted into blunt ends via exonuclease/polymerase activities. After adenylation of 3’ ends of DNA fragments, NEBNext Adaptor with hairpin loop structure was ligated to prepare for hybridization. In order to select cDNA fragments preferentially 250-300 bp in length, the library fragments were purified with AMPure XP system (Beckman Coulter, Beverly, USA). Then 3 µl USER Enzyme (NEB, USA) was used with size-selected, adaptor-ligated cDNA at 37 ℃ for 15 min followed by 5 min at 95 ℃ before PCR. Then PCR was performed with Phusion High-Fidelity DNA polymerase, Universal PCR primers and Index (X) Primer. At last, PCR products were purified (AMPure XP system) and library quality was assessed on the Agilent Bioanalyzer 2100 system. The clustering of the index-coded samples was performed on a cBot Cluster Generation System using TruSeq PE Cluster Kit v3-cBot-HS (Illumia) according to the manufacturer’s instructions. After cluster generation, the library preparations were sequenced on an Illumina HiSeq X Ten platform and paired-end reads were generated.

### PacBio cDNA library construction and sequencing

The Isoform sequencing (Iso-Seq) library was prepared according to the Isoform Sequencing protocol (Iso-Seq) using the Clontech SMARTer^®^ PCR cDNA Synthesis Kit (Clontech Laboratories (now Takara Laboratories), 634926, Mountain View, CA, USA) and the BluePippin Size Selection System protocol as described by Pacific Biosciences (PN 100-092-800-03). Briefly, Oligo(dT) enriched mRNA was reversely transcribed to cDNA by SMARTer PCR cDNA Synthesis Kit; the synthesized cDNA was then amplified by polymerase chain reaction (PCR) using BluePippin Size Selection System protocol; the Iso-Seq library was constructed by full-length cDNA damage repair, terminal repair and attaching SMRT dumbbell adapters; the sequences of the unattached adapters at both ends of the cDNA were removed by exonuclease digestion; the cDNA obtained above was combined with primers and DNA polymerase to form a complete SMRT bell library. While the library was qualified, the PacBio Sequel II platform was used for sequencing based on the effective concentration and data output requirements of the library.

### Data Filtering and Processing

The Illumina sequencing raw reads of fastq format were firstly processed through in-house perl scripts. In this step, clean data were obtained by removing reads containing adapter, reads containing ploy-N and low-quality reads from raw data. At the same time, Q20, Q30, GC-content and sequence duplication levels of the clean data were calculated. All the downstream analyses were based on clean data with high quality.

The PacBio sequencing raw data were processed by SMRTlink v8.0 software. Circular consensus sequence (CCS) was generated from subread BAM files, parameters: min_length 50, min_passes 1, max_length 15,000. CCS.BAM files were output, which were then classified into full-length and non-full-length reads using lima, removing polyA using refine. Full-length fasta files produced were then fed into the cluster step, which does isoform-level hierarchical clustering (n*log(n)), followed by final Arrow polishing, hq_quiver_min_accuracy 0.99, bin_by_primer false, bin_size_kb 1, qv_trim_5p 100, qv_trim_3p 30.

### Coral and Symbiodiniaceae sequences separation

Aligned consensus reads to coral or Symbiodiniaceae reference genomes respectivly using GMAP v2017-06-20 software [155]. The sequences mapped to Symbiodiniaceae reference genomes belonged to Symbiodiniaceae sequences, the sequences mapped to coral reference genomes belonged to coral sequences.

### Correction and de-redundancy

The RNA-seq data sequenced by the Illumina HiSeq X Ten platform was used to correct additional nucleotide errors in polish consensus sequences obtained in the previous step with LoRDEC v0.7 software [156]. Using CD-HIT v4.6.8 software (parameters: -c 0.95 -T 6 -G 0 - aL 0.00 -aS 0.99)all redundancies were removed in corrected consensus reads to acquire final full-length transcripts and unigenes for subsequent bioinformatics analysis [157].

### Gene functional annotation

Gene functions were annotated using the following databases: NT (NCBI non-redundant nucleotide sequences); NR (NCBI non-redundant protein sequences); Pfam (Protein family); KOG/COG database (Clusters of Orthologous Groups of proteins); Swiss-Prot (A manually annotated and reviewed protein sequence database); KEGG (Kyoto Encyclopedia of Genes and Genomes); GO (Gene Ontology). We used BLAST 2.7.1+ software [158] with the e-value ‘1e-5’ for NT database analysis, Diamond v0.8.36 BLASTX software [159] with the e-value ‘1e-5’ for NR, KOG, Swiss-Prot and KEGG databases analysis, and HMMER 3.1 package [160] for Pfam database analysis.

### Gene structure analysis

ANGEL v2.4 software [161] was used to predict protein CDS (Coding sequence). We used the same species or closely related species confident protein sequences for ANGEL training and then ran the ANGEL prediction for given sequences. Usually, the TFs were identified based on the Pfam files of TF families in AnimalTFDB 3.0 database [162], however, corals were not included in this database, so we identified coral TFs based on the Pfam files of TF families using hmmsearch program in HMMER 3.1 package. SSR of the transcriptomes was identified using MISA v1 [163]. We used CNCI v2 [164], CPC2 v0.1 [165], PfamScan v1.6 [166] and PLEK v1.2 [167] four tools to predict the coding potential of transcripts. Transcripts predicted with coding potential by either/all of the three tools above were filtered out, and those lacking coding potential were our candidate set of lncRNAs.

### Gene expression quantification

The full-length transcriptome obtained above was used as the reference background, and then the clean reads of each sample obtained by Illumina sequencing were mapped to it by bowtie2 V2.3.4 software [168]. The alignment results were estimated by RSEM v1.3.0 software [169] to further obtain the read count values for each transcript, and then transfered read count to FPKM for analyzing the gene expression levels. Using Pearson correlation coefficient analyzed the relationship among the samples.

### Gene differential expression analysis

The unigenes with the same annotation result in the NR database were merged to form a new read count expression matrix. Differential expression analysis of two groups was performed using the DESeq2 R package (1.30.1) [170]. DESeq2 provides statistical routines for determining differential expression in digital gene expression data using a model based on the negative binomial distribution. The resulting p-values were adjusted using Benjamini and Hochberg’s approach for controlling the false discovery rate and were named as padj. Coral genes with padj<0.001&|log2(FoldChange)|>=2 and 10 by DESeq2 were assigned as differentially expressed. Venn diagrams were drawn by VennDiagram R package (1.6.20) and GO classification bar charts were drawn by ggplot2 R package (3.3.5) [171].

### Phylogenetic Analysis

To construct the phylogenetic tree, we selected core clock genes in our own 5 reef-building corals (including a new coral *Montipora capricomis* for verification), and 22 species belonging to 17 classes in the NCBI database (Table 5; Additional file 10: Table S9). The sequences alignment was performed using ClustalW by MEGA X software [172], the sites with two or more gaps in more than 50% species were deleted, repeated the above steps until the missing data were cleaned completely. The trees were constructed using the Neighbor-Joining method [173] under the p-distances model with bootstrapping (2000). All positions with less than 50% site coverage were eliminated, i.e., fewer than 50% alignment gaps, missing data, and ambiguous bases were allowed at any position (partial deletion option). iTOL v6.3.2 [174] was used to visualize the tree.

## Supporting information

Additional file 2: Table S1. Detailed sequence data of four reef-building corals in the NCBI database

Additional file 3: Table S2. Data processing statistics of four reef-building corals

Additional file 4: Table S3. Gene function annotation results statistics

Additional file 5: Table S4. Gene structural analysis results statistics

Additional file 6: Table S5. Gene expression level analysis results statistics

Additional file 7: Table S6. Analysis of differential gene expression in corals

Additional file 8: Table S7. Kernel biomineralization-related genes in four reef-building corals

Additional file 9: Table S8. Symbiodiniaceae bioinformatics analysis

Additional file 10: Table S9. Detailed kernel circadian clock genes

Additional file 11: Table S10. Convergent expression of genes between corals and amphioxus

## Supplementary Information

**Additional file 1: Fig. S1** Overview of PacBio Sequel II SMRT sequencing data processing. **Fig. S2** Intermediate processes of PacBio Sequel II SMRT sequencing data processing. **Fig. S3** GO classifications of coral genes in *P. damicornis*. **Fig. S4** GO classifications of coral genes in *P. verrucosa*. **Fig. S5** GO classifications of coral genes in *A. muricata*. **Fig. S6** GO classifications of coral genes in *M. foliosa*. **Fig. S7** KOG classifications of coral genes. **Fig. S8** KEGG metabolic pathway classifications of coral genes in *P. damicornis*. **Fig. S9** KEGG metabolic pathway classifications of coral genes in *P. verrucosa*. **Fig. S10** KEGG metabolic pathway classifications of coral genes in *A. muricata*. **Fig. S11** KEGG metabolic pathway classifications of coral genes in *M. foliosa*. **Fig. S12** Summary of coral gene structural analysis. **Fig. S13** FPKM box plot of each coral gene expression. **Fig. S14** Venn plots of coral differentially expressed genes. **Fig. S15** Summary of Symbiodiniaceae gene functional annotation. **Fig. S16** NR database annotation of Symbiodiniaceae. **Fig. S17** GO classifications of Symbiodiniaceae genes in *P. damicornis*. **Fig. S18** GO classifications of Symbiodiniaceae genes in *P. verrucosa*. **Fig. S19** GO classifications of Symbiodiniaceae genes in *A. muricata*. **Fig. S20** GO classifications of Symbiodiniaceae genes in *M. foliosa*. **Fig. S21** KOG classifications of Symbiodiniaceae genes. **Fig. S22** KEGG metabolic pathway classifications of Symbiodiniaceae genes in *P. damicornis*. **Fig. S23** KEGG metabolic pathway classifications of Symbiodiniaceae genes in *P. verrucosa*. **Fig. S24** KEGG metabolic pathway classifications of Symbiodiniaceae genes in *A. muricata*. **Fig. S25** KEGG metabolic pathway classifications of Symbiodiniaceae genes in *M. foliosa*. **Fig. S26** Summary of Symbiodiniaceae gene structural analysis. **Fig. S27** Venn plots of statistics on the number of Symbiodiniaceae sequences expressed in four reef-building corals. **Fig. S28** Summary of the Symbiodiniaceae gene expression level analysis. **Fig. S29** Bar graph of coexpressed Symbiodiniaceae sequences based on GO molecular functions and cellular components categories. **Fig. S30** Conserved domains of *cry1* or *2* genes among different species. **Fig. S31** Phylogenetic tree of *cry1* or *2* genes based on the Neighbor-Joining method. **Fig. S32** Conserved domains of *Clock* or *Npas2* genes among different species. **Fig. S33** Phylogenetic tree of *Clock* or *Npas2* genes based on the Neighbor-Joining method. **Fig. S34** Conserved domains of *cyc* or *Arntl* genes among different species. **Fig. S35** Phylogenetic tree of *cyc* or *Arntl* genes based on the Neighbor-Joining method. **Fig. S36** *Per1* or *2* or *3* gene phylogenetic analysis.

**Additional file 2: Table S1.** Detailed sequence data of four reef-building corals in the NCBI database.

**Additional file 3: Table S2.** Data processing statistics of four reef-building corals.

**Additional file 4: Table S3.** Gene function annotation results statistics.

**Additional file 5: Table S4.** Gene structural analysis results statistics.

**Additional file 6: Table S5.** Gene expression level analysis results statistics.

**Additional file 7: Table S6.** Analysis of differential gene expression in corals.

**Additional file 8: Table S7.** Kernel biomineralization-related genes in four reef-building corals.

**Additional file 9: Table S8.** Symbiodiniaceae bioinformatics analysis.

**Additional file 10: Table S9.** Detailed kernel circadian clock genes.

**Additional file 11: Table S10.** Convergent expression of genes between corals and amphioxus.

## Declarations

### Consent for publication

Not applicable.

### Availability of data and materials

The datasets (reef-building coral holobionts full-length and short-read transcriptome sequencing raw data) generated during the current study are available at the Sequence Read Archive (SRA) publicly available repository, [https://www.ncbi.nlm.nih.gov/sra/]. The accession numbers are SRR9613489 and SRR12904788-90 for *Pocillopora damicornis*, SRR12963486 and SRR12959238-40 for *Pocillopora verrucosa*, SRR12963485 and SRR12959195, SRR12959206, SRR12959217 for *Acropora muricata*, SRR12963484 and SRR12959182-4 for *Montipora foliosa*.

The datasets (Unigene, CDS, PEP and UTR sequences) analysed during the current study are available in https://doi.org/10.6084/m9.figshare.19403021.

### Authors’ contributions

CH conceived the project. ZL, XL, and JC made suggestions for the project. TH performed the experiment, data analysis, writing and editing. All authors discussed the results and commented on the manuscript. The authors read and approved the final manuscript.

## Acknowledgments

We thank M. Zhu from the Nanjing Institute of Geology and Paleontology, CAS for scientific guidance, M. Zhu, J. Lu, and Z. Gai from the Institute of Vertebrate Paleontology and Paleoanthropology, CAS for technical and scientific guidance, and Y. Loya from the Israel Academy of Sciences for scientific guidance.

## References

1. Hoegh-Guldberg O. Climate change, coral bleaching and the future of the world’s coral reefs. Mar Freshw Res. 1999;50(8):839.

2. Copper P. Ancient reef ecosystem expansion and collapse. Coral Reefs. 1994;13(1):3–11.

3. Reaka-Kudla ML. The global biodiversity of coral reefs: a comparison with rain forests. In: Reaka-Kudla ML, Wilson DE, Wilson EO, editors. Biodiversity II: Understanding and Protecting Our Biological Resources. Washington, D.C.: Joseph Henry Press; 1997. p. 83–108.

4. Smith SV. Coral-reef area and the contributions of reefs to processes and resources of the world’s oceans. Nature. 1978;273(5659):225-6.

5. Spalding MD, Grenfell AM. New estimates of global and regional coral reef areas. Coral Reefs. 1997;16(4):225–30.

6. Carpenter KE, Abrar M, Aeby G, Aronson RB, Banks S, Bruckner A, et al. One-Third of Reef-Building Corals Face Elevated Extinction Risk from Climate Change and Local Impacts. Science. 2008;321(5888):560-3.

7. Moberg F, Folke C. Ecological goods and services of coral reef ecosystems. Ecol Econ. 1999;29(2):215–33.

8. Normile D. Bringing Coral Reefs Back From the Living Dead. Science. 2009;325(5940):559-61.

9. Wilkinson C. Status of coral reefs of the world: 2008. Townsville, Australia: Global Coral Reef Monitoring Network and Reef and Rainforest Research Centre; 2008.

10. Yu K. Coral reefs in the South China Sea: Their response to and records on past environmental changes. Sci China Earth Sci. 2012;55(8):1217–29.

11. Miller DJ, Ball EE, Technau U. Cnidarians and ancestral genetic complexity in the animal kingdom. Trends Genet. 2005;21(10):536–9.

12. Finnerty JR. Origins of Bilateral Symmetry: Hox and Dpp Expression in a Sea Anemone. Science. 2004;304(5675):1335-7.

13. Genikhovich G, Technau U. On the evolution of bilaterality. Development. 2017;144(19):3392–404.

14. Falini G, Fermani S, Goffredo S. Coral biomineralization: A focus on intra-skeletal organic matrix and calcification. Semin Cell Dev Biol. 2015;46:17–26.

15. Allemand D, Tambutté É, Zoccola D, Tambutté S. Coral Calcification, Cells to Reefs. In: Dubinsky Z, Stambler N, editors. Coral Reefs: An Ecosystem in Transition. Springer, Dordrecht; 2011. p. 119-50.

16. Aranda M, Li Y, Liew YJ, Baumgarten S, Simakov O, Wilson MC, et al. Genomes of coral dinoflagellate symbionts highlight evolutionary adaptations conducive to a symbiotic lifestyle. Sci Rep. 2016;6:39734.

17. Gonzalez-Pech RA, Stephens TG, Chen YB, Mohamed AR, Cheng YY, Shah S, et al. Comparison of 15 dinoflagellate genomes reveals extensive sequence and structural divergence in family Symbiodiniaceae and genus Symbiodinium. BMC Biol. 2021;19:73.

18. Lin S, Cheng S, Song B, Zhong X, Lin X, Li W, et al. The Symbiodinium kawagutii genome illuminates dinoflagellate gene expression and coral symbiosis. Science. 2015;350(6261):691-4.

19. Liu HL, Stephens TG, Gonzalez-Pech RA, Beltran VH, Lapeyre B, Bongaerts P, et al. Symbiodinium genomes reveal adaptive evolution of functions related to coral-dinoflagellate symbiosis. Commun Biol. 2018;1:95.

20. Shoguchi E, Beedessee G, Tada I, Hisata K, Kawashima T, Takeuchi T, et al. Two divergent Symbiodinium genomes reveal conservation of a gene cluster for sunscreen biosynthesis and recently lost genes. BMC Genomics. 2018;19:458.

21. Shoguchi E, Shinzato C, Kawashima T, Gyoja F, Mungpakdee S, Koyanagi R, et al. Draft assembly of the Symbiodinium minutum nuclear genome reveals dinoflagellate gene structure. Curr Biol. 2013;23(15):1399–408.

22. Gardner SG, Camp EF, Smith DJ, Kahlke T, Osman EO, Gendron G, et al. Coral microbiome diversity reflects mass coral bleaching susceptibility during the 2016 El Nino heat wave. Ecol Evol. 2019;9(3):938–56.

23. Zhou Z, Zhang G, Chen G, Ni X, Guo L, Yu X, et al. Elevated ammonium reduces the negative effect of heat stress on the stony coral Pocillopora damicornis. Mar Pollut Bull. 2017;118(1-2):319–27.

24. Zhang Y, Zhou Z, Wang L, Huang B. Transcriptome, expression, and activity analyses reveal a vital heat shock protein 70 in the stress response of stony coral Pocillopora damicornis. Cell Stress Chaperones. 2018;23(4):711–21.

25. Mollica NR, Guo W, Cohen AL, Huang KF, Foster GL, Donald HK, et al. Ocean acidification affects coral growth by reducing skeletal density. Proc Natl Acad Sci U S A. 2018;115(8):1754–9.

26. Kline DI, Teneva L, Okamoto DK, Schneider K, Caldeira K, Miard T, et al. Living coral tissue slows skeletal dissolution related to ocean acidification. Nat Ecol Evol. 2019;3(10):1438–44.

27. Hu M, Zheng X, Fan CM, Zheng Y. Lineage dynamics of the endosymbiotic cell type in the soft coral Xenia. Nature. 2020;582(7813):534-8.

28. Levy S, Elek A, Grau-Bove X, Menendez-Bravo S, Iglesias M, Tanay A, et al. A stony coral cell atlas illuminates the molecular and cellular basis of coral symbiosis, calcification, and immunity. Cell. 2021;184(11):2973–87.

29. Orgel J, Sella I, Madhurapantula RS, Antipova O, Mandelberg Y, Kashman Y, et al. Molecular and ultrastructural studies of a fibrillar collagen from octocoral (Cnidaria). J Exp Biol. 2017;220:3327–35.

30. Tang J, Ni X, Zhou Z, Wang L, Lin S. Acute microplastic exposure raises stress response and suppresses detoxification and immune capacities in the scleractinian coral Pocillopora damicornis. Environ Pollut. 2018;243:66–74.

31. Zhou Z, Zhao S, Tang J, Liu Z, Wu Y, Wang Y, et al. Altered Immune Landscape and Disrupted Coral-Symbiodinium Symbiosis in the Scleractinian Coral Pocillopora damicornis by Vibrio coralliilyticus Challenge. Front Physiol. 2019;10:366.

32. Cunning R, Bay RA, Gillette P, Baker AC, Traylor-Knowles N. Comparative analysis of the Pocillopora damicornis genome highlights role of immune system in coral evolution. Sci Rep. 2018;8(1):16134.

33. Guo Z, Liao X, Chen J, He C, Lu, Z. Binding pattern reconstructions of FGF-FGFR budding-inducing signaling in reef-building corals. Front Physiol. 2022;12:759370.

34. Li YX, Han TY, Bi K, Liang K, Chen JY, Lu J, et al. The 3D Reconstruction of Pocillopora Colony Sheds Light on the Growth Pattern of This Reef-Building Coral. iScience. 2020;23(6):101069.

35. Shoguchi E, Tanaka M, Shinzato C, Kawashima T, Satoh N. A genome-wide survey of photoreceptor and circadian genes in the coral, Acropora digitifera. Gene. 2013;515(2):426–31.

36. Levy O, Appelbaum L, Leggat W, Gothlif Y, Hayward DC, Miller DJ, et al. Light-responsive cryptochromes from a simple multicellular animal, the coral Acropora millepora. Science. 2007;318(5849):467-70.

37. Vize PD. Transcriptome analysis of the circadian regulatory network in the coral Acropora millepora. Biol Bull. 2009;216(2):131–7.

38. Wang X, Zoccola D, Liew YJ, Tambutte E, Cui G, Allemand D, et al. The Evolution of Calcification in Reef-Building Corals. Mol Biol Evol. 2021;38(9):3543–55.

39. Zoccola D, Tambutte E, Kulhanek E, Puverel S, Scimeca JC, Allemand D, et al. Molecular cloning and localization of a PMCA P-type calcium ATPase from the coral Stylophora pistillata. Biochim Biophys Acta. 2004;1663(1-2):117–26.

40. Zoccola D, Tambutte E, Senegas-Balas F, Michiels JF, Failla JP, Jaubert J, et al. Cloning of a calcium channel alpha1 subunit from the reef-building coral, Stylophora pistillata. Gene. 1999;227(2):157–67.

41. Bertucci A, Moya A, Tambutte S, Allemand D, Supuran CT, Zoccola D. Carbonic anhydrases in anthozoan corals-A review. Bioorg Med Chem. 2013;21(6):1437–50.

42. Venn AA, Tambutte E, Lotto S, Zoccola D, Allemand D, Tambutte S. Imaging intracellular pH in a reef coral and symbiotic anemone. Proc Natl Acad Sci U S A. 2009;106(39):16574–9.

43. Hopkinson BM, Tansik AL, Fitt WK. Internal carbonic anhydrase activity in the tissue of scleractinian corals is sufficient to support proposed roles in photosynthesis and calcification. J Exp Biol. 2015;218(13):2039–48.

44. Zoccola D, Ganot P, Bertucci A, Caminiti-Segonds N, Techer N, Voolstra CR, et al. Bicarbonate transporters in corals point towards a key step in the evolution of cnidarian calcification. Sci Rep. 2015;5:9983.

45. Mass T, Drake JL, Haramaty L, Kim JD, Zelzion E, Bhattacharya D, et al. Cloning and Characterization of Four Novel Coral Acid-Rich Proteins that Precipitate Carbonates In Vitro. Curr Biol. 2013;23(12):1126–31.

46. Puverel S, Tambutté E, Pereira-Mouriès L, Zoccola D, Allemand D, Tambutte S. Soluble organic matrix of two Scleractinian corals: partial and comparative analysis. Comp Biochem Physiol B Biochem Mol Biol. 2005;141(4):480–7.

47. Bhattacharya D, Agrawal S, Aranda M, Baumgarten S, Belcaid M, Drake JL, et al. Comparative genomics explains the evolutionary success of reef-forming corals. Elife. 2016;5:e13288.

48. Drake JL, Mass T, Haramaty L, Zelzion E, Bhattacharya D, Falkowski PG. Proteomic analysis of skeletal organic matrix from the stony coral Stylophora pistillata. Proc Natl Acad Sci U S A. 2013;110(10):3788–93.

49. Ramos-Silva P, Kaandorp J, Huisman L, Marie B, Zanella-Cleon I, Guichard N, et al. The skeletal proteome of the coral Acropora millepora: the evolution of calcification by co-option and domain shuffling. Mol Biol Evol. 2013;30(9):2099–112.

50. Brocks JJ, Jarrett AJM, Sirantoine E, Hallmann C, Hoshino Y, Liyanage T. The rise of algae in Cryogenian oceans and the emergence of animals. Nature. 2017;548(7669):578-81.

51. Mills DB. The origin of phagocytosis in Earth history. Interface Focus. 2020;10(4):20200019.

52. Knoll AH. Biogeochemistry: Food for early animal evolution. Nature. 2017;548(7669):528-30.

53. Cong P, Ma X, Williams M, Siveter DJ, Siveter DJ, Gabbott SE, et al. Host-specific infestation in early Cambrian worms. Nat Ecol Evol. 2017;1(10):1465–9.

54. Davy SK, Allemand D, Weis VM. Cell Biology of Cnidarian-Dinoflagellate Symbiosis. Microbiol Mol Biol Rev. 2012;76(2):229–61.

55. Sebe-Pedros A, Saudemont B, Chomsky E, Plessier F, Mailhe MP, Renno J, et al. Cnidarian Cell Type Diversity and Regulation Revealed by Whole-Organism Single-Cell RNA-Seq. Cell. 2018;173(6):1520–34.

56. Guibert I, Lecellier G, Torda G, Pochon X, Berteaux-Lecellier V. Metabarcoding reveals distinct microbiotypes in the giant clam Tridacna maxima. Microbiome. 2020;8(1):57.

57. He CP, Han TY, Liao X, Zhou YX, Wang XQ, Guan R, et al. Phagocytic intracellular digestion in amphioxus (Branchiostoma). Proc Biol Sci. 2018;285(1880):20180438.

58. Weis VM. Cell Biology of Coral Symbiosis: Foundational Study Can Inform Solutions to the Coral Reef Crisis. Integr Comp Biol. 2019;59(4):845–55.

59. Kobayashi J, Kubota T. Bioactive macrolides and polyketides from marine dinoflagellates of the genus Amphidinium. J Nat Prod. 2007;70(3):451–60.

60. Murata M, Yasumoto T. The structure elucidation and biological activities of high molecular weight algal toxins: maitotoxin, prymnesins and zooxanthellatoxins. Nat Prod Rep. 2000;17(3):293–314.

61. Hellyer SD. Chapter Four-Marine-derived nicotinic receptor antagonist toxins: Pinnatoxins and alpha conotoxins. In: Novelli A, Fernández-Sánchez M-T, Aschner M, Costa LG, editors. Advances in Neurotoxicology, vol 6. Academic Press; 2021. p. 105–91.

62. Reitzel AM, Behrendt L, Tarrant AM. Light Entrained Rhythmic Gene Expression in the Sea Anemone Nematostella vectensis: The Evolution of the Animal Circadian Clock. Plos One. 2010;5(9): e12805.

63. Jindrich K, Roper KE, Lemon S, Degnan BM, Reitzel AM, Degnan SM. Origin of the Animal Circadian Clock: Diurnal and Light-Entrained Gene Expression in the Sponge Amphimedon queenslandica. Front Mar Sci. 2017;4:327.

64. Lowrey PL, Takahashi JS. Mammalian circadian biology: elucidating genome-wide levels of temporal organization. Annu Rev Genomics Hum Genet. 2004;5:407–41.

65. Yang Z, Liu B, Su J, Liao J, Lin C, Oka Y. Cryptochromes Orchestrate Transcription Regulation of Diverse Blue Light Responses in Plants. Photochem Photobiol. 2017;93(1):112–27.

66. Dunlap JC. Molecular bases for circadian clocks. Cell. 1999;96(2):271–90.

67. Krishnan B, Levine JD, Lynch MKS, Dowse HB, Funes P, Hall JC, et al. A new role for cryptochrome in a Drosophila circadian oscillator. Nature. 2001;411(6835):313-7.

68. Griffin EA, Jr., Staknis D, Weitz CJ. Light-independent role of CRY1 and CRY2 in the mammalian circadian clock. Science. 1999;286(5440):768-71.

69. Zhu HS, Sauman I, Yuan Q, Casselman A, Emery-Le M, Emery P, et al. Cryptochromes define a novel circadian clock mechanism in monarch butterflies that may underlie sun compass navigation. PLoS Biol. 2008;6(1):138–55.

70. Zhu H, Yuan Q, Briscoe AD, Froy O, Casselman A, Reppert SM. The two CRYs of the butterfly. Curr Biol. 2005;15(23):R953–4.

71. Hoeksema BW, Rogers A, Quibilan MC. Pocillopora damicornis. The IUCN Red List of Threatened Species 2014: e.T133222A54216898. https://dx.doi.org/10.2305/IUCN.UK.2014-1.RLTS.T133222A54216898.en. Accessed on 11 December 2021.

72. Richards ZT, Delbeek JT, Lovell ER, Bass D, Aeby G, Reboton C. Acropora formosa. The IUCN Red List of Threatened Species 2014: e.T133644A54299282. https://dx.doi.org/10.2305/IUCN.UK.2014-1.RLTS.T133644A54299282.en. Accessed on 11 December 2021.

73. DeVantier L, Hodgson G, Huang D, Johan O, Licuanan A, Obura DO, et al. Montipora foliosa. The IUCN Red List of Threatened Species 2014: e.T133071A54189890. https://dx.doi.org/10.2305/IUCN.UK.2014-1.RLTS.T133071A54189890.en. Accessed on 11 December 2021.

74. Hoeksema BW, Rogers A, Quibilan MC. Pocillopora verrucosa. The IUCN Red List of Threatened Species 2014: e.T133197A54210095. https://dx.doi.org/10.2305/IUCN.UK.2014-1.RLTS.T133197A54210095.en. Accessed on 11 December 2021.

75. Van Dijk EL, Jaszczyszyn Y, Naquin D, Thermes C. The Third Revolution in Sequencing Technology. Trends Genet. 2018;34(9):666–81.

76. Lu H, Giordano F, Ning Z. Oxford Nanopore MinION Sequencing and Genome Assembly. Genomics Proteomics Bioinformatics. 2016;14(5):265–79.

77. Rhoads A, Au KF. PacBio Sequencing and Its Applications. Genomics Proteomics Bioinformatics. 2015;13(5):278–89.

78. Vidal-Dupiol J, Zoccola D, Tambutté E, Grunau C, Cosseau C, Smith KM, et al. Genes Related to Ion-Transport and Energy Production Are Upregulated in Response to CO2-Driven pH Decrease in Corals: New Insights from Transcriptome Analysis. PLoS One. 2013;8(3):e58652.

79. Mayfield AB, Wang YB, Chen CS, Lin CY, Chen SH. Compartment-specific transcriptomics in a reef-building coral exposed to elevated temperatures. Mol Ecol. 2014;23(23):5816–30.

80. Yuan C, Zhou Z, Zhang Y, Chen G, Yu X, Ni X, et al. Effects of elevated ammonium on the transcriptome of the stony coral Pocillopora damicornis. Mar Pollut Bull. 2017;114(1):46–52.

81. Zhou Z, Yu X, Tang J, Wu Y, Wang L, Huang B. Systemic response of the stony coral Pocillopora damicornis against acute cadmium stress. Aquat Toxicol. 2018;194:132–9.

82. Mass T, Putnam HM, Drake JL, Zelzion E, Gates RD, Bhattacharya D, et al. Temporal and spatial expression patterns of biomineralization proteins during early development in the stony coral Pocillopora damicornis. Proc Biol Sci. 2016;283(1829):20160322.

83. Crowder CM, Meyer E, Fan TY, Weis VM. Impacts of temperature and lunar day on gene expression profiles during a monthly reproductive cycle in the brooding coral Pocillopora damicornis. Mol Ecol. 2017;26(15):3913–25.

84. Wecker P, Lecellier G, Guibert I, Zhou Y, Bonnard I, Berteaux-Lecellier V. Exposure to the environmentally-persistent insecticide chlordecone induces detoxification genes and causes polyp bail-out in the coral P. damicornis. Chemosphere. 2018;195:190–200.

85. Johnston EC, Forsman ZH, Flot J-F, Schmidt-Roach S, Pinzón JH, Knapp ISS, et al. A genomic glance through the fog of plasticity and diversification in Pocillopora. Sci Rep. 2017;7(1):1–11.

86. Quattrini AM, Faircloth BC, Duenas LF, Bridge TCL, Brugler MR, Calixto-Botia IF, et al. Universal target-enrichment baits for anthozoan (Cnidaria) phylogenomics: New approaches to long-standing problems. Mol Ecol Resour. 2018;18(2):281–95.

87. Flot JF, Tillier S. The mitochondrial genome of Pocillopora (Cnidaria: Scleractinia) contains two variable regions: the putative D-loop and a novel ORF of unknown function. Gene. 2007;401(1-2):80–7.

88. Walters BM, Connelly MT, Young B, Traylor-Knowles N. The Complicated Evolutionary Diversification of the Mpeg-1/Perforin-2 Family in Cnidarians. Front Immunol. 2020;11:1690.

89. Morikawa MK, Palumbi SR. Using naturally occurring climate resilient corals to construct bleaching-resistant nurseries. Proc Natl Acad Sci U S A. 2019;116(21):10586–91.

90. Buitrago-Lopez C, Mariappan KG, Cardenas A, Gegner HM, Voolstra CR. The Genome of the Cauliflower Coral Pocillopora verrucosa. Genome Biol Evol. 2020;12(10):1911–7.

91. Lee STM, Keshavmurthy S, Fontana S, Takuma M, Chou WH, Chen CA. Transcriptomic response in Acropora muricata under acute temperature stress follows preconditioned seasonal temperature fluctuations. BMC Res Notes. 2018;11(1):119.

92. Shinzato C, Khalturin K, Inoue J, Zayasu Y, Kanda M, Kawamitsu M, et al. Eighteen Coral Genomes Reveal the Evolutionary Origin of Acropora Strategies to Accommodate Environmental Changes. Mol Biol Evol. 2021;38(1):16–30.

93. Shinzato C, Zayasu Y, Kanda M, Kawamitsu M, Satoh N, Yamashita H, et al. Using Seawater to Document Coral-Zoothanthella Diversity: A New Approach to Coral Reef Monitoring Using Environmental DNA. Front Mar Sci. 2018;5:28.

94. Ceh J, Raina JB, Soo RM, van Keulen M, Bourne DG. Coral-Bacterial Communities before and after a Coral Mass Spawning Event on Ningaloo Reef. PLoS One. 2012;7(5):e36920.

95. Mhuantong W, Nuryadi H, Trianto A, Sabdono A, Tangphatsornruang S, Eurwilaichitr L, et al. Comparative analysis of bacterial communities associated with healthy and diseased corals in the Indonesian sea. PeerJ. 2019;7:e8137.

96. Diaz JM, Hansel CM, Apprill A, Brighi C, Zhang T, Weber L, et al. Species-specific control of external superoxide levels by the coral holobiont during a natural bleaching event. Nat Commun. 2016;7:13801.

97. Beatty DS, Clements CS, Stewart FJ, Hay ME. Intergenerational effects of macroalgae on a reef coral: major declines in larval survival but subtle changes in microbiomes. Mar Ecol Prog Ser. 2018;589:97–114.

98. Beatty DS, Valayil JM, Clements CS, Ritchie KB, Stewart FJ, Hay ME. Variable effects of local management on coral defenses against a thermally regulated bleaching pathogen. Sci Adv. 2019;5(10):eaay1048.

99. Epstein HE, Torda G, Munday PL, van Oppen MJH. Parental and early life stage environments drive establishment of bacterial and dinoflagellate communities in a common coral. ISME J. 2019;13(6):1635–8.

100. van Oppen MJH, Bongaerts P, Frade P, Peplow LM, Boyd SE, Nim HT, et al. Adaptation to reef habitats through selection on the coral animal and its associated microbiome. Mol Ecol. 2018;27(14):2956–71.

101. Osman EO, Suggett DJ, Voolstra CR, Pettay DT, Clark DR, Pogoreutz C, et al. Coral microbiome composition along the northern Red Sea suggests high plasticity of bacterial and specificity of endosymbiotic dinoflagellate communities. Microbiome. 2020;8(1):8.

102. Li J, Long L, Zou Y, Zhang S. Microbial community and transcriptional responses to increased temperatures in coral Pocillopora damicornis holobiont. Environ Microbiol. 2021;23(2):826–43.

103. Brener-Raffalli K, Clerissi C, Vidal-Dupiol J, Adjeroud M, Bonhomme F, Pratlong M, et al. Thermal regime and host clade, rather than geography, drive Symbiodinium and bacterial assemblages in the scleractinian coral Pocillopora damicornis sensu lato. Microbiome. 2018;6:39.

104. Chen B, Yu K, Liao Z, Yu X, Qin Z, Liang J, et al. Microbiome community and complexity indicate environmental gradient acclimatisation and potential microbial interaction of endemic coral holobionts in the South China Sea. Sci Total Environ. 2021;765:142690.

105. Sun F, Yang H, Wang G, Shi Q. Combination Analysis of Metatranscriptome and Metagenome Reveal the Composition and Functional Response of Coral Symbionts to Bleaching During an El Nino Event. Front Microbiol. 2020;11:448.

106. Ziegler M, Grupstra CGB, Barreto MM, Eaton M, BaOmar J, Zubier K, et al. Coral bacterial community structure responds to environmental change in a host-specific manner. Nat Commun. 2019;10(1):3092.

107. Yang Q, Zhang Y, Ahmad M, Ling J, Zhou W, Zhang Y, et al. Microbial community structure shifts and potential Symbiodinium partner bacterial groups of bleaching coral Pocillopora verrucosa in South China Sea. Ecotoxicology. 2021;30(5):966–74.

108. Pogoreutz C, Radecker N, Cardenas A, Gardes A, Wild C, Voolstra CR. Dominance of Endozoicomonas bacteria throughout coral bleaching and mortality suggests structural inflexibility of the Pocillopora verrucosa microbiome. Ecol Evol. 2018;8(4):2240–52.

109. Neave MJ, Michell CT, Apprill A, Voolstra CR. Endozoicomonas genomes reveal functional adaptation and plasticity in bacterial strains symbiotically associated with diverse marine hosts. Sci Rep. 2017;7:40579.

110. Ziegler M, Roik A, Porter A, Zubier K, Mudarris MS, Ormond R, et al. Coral microbial community dynamics in response to anthropogenic impacts near a major city in the central Red Sea. Mar Pollut Bull. 2016;105(2):629–40.

111. Neave MJ, Rachmawati R, Xun L, Michell CT, Bourne DG, Apprill A, et al. Differential specificity between closely related corals and abundant Endozoicomonas endosymbionts across global scales. ISME J. 2017;11(1):186–200.

112. van de Water JAJM, Courtial L, Houlbrèque F, Jacquet S, Ferrier-Pagès C. Ultra-Violet Radiation Has a Limited Impact on Seasonal Differences in the Acropora Muricata Holobiont. Front Mar Sci. 2018;5:275.

113. Sweet M, Bythell J. White syndrome in Acropora muricata: nonspecific bacterial infection and ciliate histophagy. Mol Ecol. 2015;24(5):1150–9.

114. Tombacz D, Sharon D, Olah P, Csabai Z, Snyder M, Boldogkoi Z. Strain Kaplan of Pseudorabies Virus Genome Sequenced by PacBio Single-Molecule Real-Time Sequencing Technology. Genome Announc. 2014;2(4):e00628–14.

115. Chin CS, Alexander DH, Marks P, Klammer AA, Drake J, Heiner C, et al. Nonhybrid, finished microbial genome assemblies from long-read SMRT sequencing data. Nat Methods. 2013;10(6):563–9.

116. Chen S, Qiu G, Yang M. SMRT sequencing of full-length transcriptome of seagrasses Zostera japonica. Sci Rep. 2019;9(1):14537.

117. Zhang SJ, Wang CQ, Yan SY, Fu AS, Luan XK, Li YM, et al. Isoform Evolution in Primates through Independent Combination of Alternative RNA Processing Events. Mol Biol Evol. 2017;34(10):2453–68.

118. Weirather JL, Afshar PT, Clark TA, Tseng E, Powers LS, Underwood JG, et al. Characterization of fusion genes and the significantly expressed fusion isoforms in breast cancer by hybrid sequencing. Nucleic Acids Res. 2015;43(18):e116.

119. Huang KK, Huang JW, Wu JKL, Lee MH, Tay ST, Kumar V, et al. Long-read transcriptome sequencing reveals abundant promoter diversity in distinct molecular subtypes of gastric cancer. Genome Biol. 2021;22(1):44.

120. Naftaly AS, Pau S, White MA. Long-read RNA sequencing reveals widespread sex-specific alternative splicing in threespine stickleback fish. Genome Res. 2021;31(8): 1486–97.

121. Li W, Jaroszewski L, Godzik A. Tolerating some redundancy significantly speeds up clustering of large protein databases. Bioinformatics. 2002;18(1):77–82.

122. Finn RD, Coggill P, Eberhardt RY, Eddy SR, Mistry J, Mitchell AL, et al. The Pfam protein families database: towards a more sustainable future. Nucleic Acids Res. 2016;44(D1):D279–85.

123. Tatusov RL, Fedorova ND, Jackson JD, Jacobs AR, Kiryutin B, Koonin EV, et al. The COG database: an updated version includes eukaryotes. BMC Bioinformatics. 2003;4:41.

124. Bairoch A, Apweiler R. The SWISS-PROT protein sequence database and its supplement TrEMBL in 2000. Nucleic Acids Res. 2000;28(1):45–8.

125. Kanehisa M, Goto S, Kawashima S, Okuno Y, Hattori M. The KEGG resource for deciphering the genome. Nucleic Acids Res. 2004;32(suppl_1):D277-80.

126. Ashburner M, Ball CA, Blake JA, Botstein D, Butler H, Cherry JM, et al. Gene ontology: tool for the unification of biology. Nat Genet. 2000;25(1):25–9.

127. Shumaker A, Putnam HM, Qiu H, Price DC, Zelzion E, Harel A, et al. Genome analysis of the rice coral Montipora capitata. Sci Rep. 2019;9(1):2571.

128. Shinzato C, Shoguchi E, Kawashima T, Hamada M, Hisata K, Tanaka M, et al. Using the Acropora digitifera genome to understand coral responses to environmental change. Nature. 2011;476(7360):320-3.

129. Ying H, Hayward DC, Cooke I, Wang W, Moya A, Siemering KR, et al. The Whole-Genome Sequence of the Coral Acropora millepora. Genome Biol Evol. 2019;11(5):1374–9.

130. Ying H, Cooke I, Sprungala S, Wang W, Hayward DC, Tang Y, et al. Comparative genomics reveals the distinct evolutionary trajectories of the robust and complex coral lineages. Genome Biol. 2018;19(1):175.

131. Meibom A, Cuif JP, Hillion FO, Constantz BR, Juillet-Leclerc A, Dauphin Y, et al. Distribution of magnesium in coral skeleton. Geophys Res Lett. 2004;31(23):L23306.

132. Li YX, Liao X, Bi K, Han TY, Chen JY, Lu J, et al. Micro-CT reconstruction reveals the colony pattern regulations of four dominant reef-building corals. Ecol Evol. 2021;11:16266–79.

133. Marat AL, McPherson PS. The connecdenn family, Rab35 guanine nucleotide exchange factors interfacing with the clathrin machinery. J Biol Chem. 2010;285(14):10627–37.

134. Heisler FF, Loebrich S, Pechmann Y, Maier N, Zivkovic AR, Tokito M, et al. Muskelin Regulates Actin Filament-and Microtubule-Based GABAA Receptor Transport in Neurons. Neuron. 2011;70(1):66–81.

135. Kielty CM, Baldock C, Lee D, Rock MJ, Ashworth JL, Shuttleworth CA. Fibrillin: from microfibril assembly to biomechanical function. Philos Trans R Soc Lond B Biol Sci. 2002;357(1418):207-17.

136. Sommer B, Oprins A, Rabouille C, Munro S. The exocyst component Sec5 is present on endocytic vesicles in the oocyte of Drosophila melanogaster. J Cell Biol. 2005;169(6):953–63.

137. Waelter S. The huntingtin interacting protein HIP1 is a clathrin and alpha-adaptin-binding protein involved in receptor-mediated endocytosis. Hum Mol Genet. 2001;10(17):1807–17.

138. Yasunaga T, Hoff S, Schell C, Helmstadter M, Kretz O, Kuechlin S, et al. The polarity protein Inturned links NPHP4 to Daam1 to control the subapical actin network in multiciliated cells. J Cell Biol. 2015;211(5):963–73.

139. Hsiao YC, Tong ZJ, Westfall JE, Ault JG, Page-McCaw PS, Ferland RJ. Ahi1, whose human ortholog is mutated in Joubert syndrome, is required for Rab8a localization, ciliogenesis and vesicle trafficking. Hum Mol Genet. 2009;18(20):3926–41.

140. Chih B, Liu P, Chinn Y, Chalouni C, Komuves LG, Hass PE, et al. A ciliopathy complex at the transition zone protects the cilia as a privileged membrane domain. Nat Cell Biol. 2012;14(1):61–72.

141. Bhogaraju S, Cajanek L, Fort C, Blisnick T, Weber K, Taschner M, et al. Molecular Basis of Tubulin Transport Within the Cilium by IFT74 and IFT81. Science. 2013;341(6149):1009-12.

142. Gerondopoulos A, Strutt H, Stevenson NL, Sobajima T, Levine TP, Stephens DJ, et al. Planar Cell Polarity Effector Proteins Inturned and Fuzzy Form a Rab23 GEF Complex. Curr Biol. 2019;29(19):3323–30.

143. Tammana TVS, Tammana D, Diener DR, Rosenbaum J. Centrosomal protein CEP104 (Chlamydomonas FAP256) moves to the ciliary tip during ciliary assembly. J Cell Sci. 2013;126(21):5018–29.

144. Kypri E, Christodoulou A, Maimaris G, Lethan M, Markaki M, Lysandrou C, et al. The nucleotide-binding proteins Nubp1 and Nubp2 are negative regulators of ciliogenesis. Cell Mol Life Sci. 2014;71(3):517–38.

145. Shimizu Y. Microalgal metabolites. Curr Opin Microbiol. 2003;6(3):236–43.

146. Morisseau C. Role of epoxide hydrolases in lipid metabolism. Biochimie. 2013;95(1):91–5.

147. De Simone G, Di Masi A, Polticelli F, Ascenzi P. Human nitrobindin: the first example of an all-β-barrel ferric heme-protein that catalyzes peroxynitrite detoxification. FEBS Open Bio. 2018;8(12):2002–10.

148. De Simone G, di Masi A, Vita GM, Polticelli F, Pesce A, Nardini M, et al. Mycobacterial and Human Nitrobindins: Structure and Function. Antioxid Redox Signal. 2020;33(4):229–46.

149. Cummins I, McAuley K, Fordham-Skelton A, Schwoerer R, Steel PG, Davis BG, et al. Unique regulation of the active site of the serine esterase S-formylglutathione hydrolase. J Mol Biol. 2006;359(2):422–32.

150. Diez-Fernandez C, Hu LY, Cervera J, Haberle J, Rubio V. Understanding carbamoyl phosphate synthetase (CPS1) deficiency by using the recombinantly purified human enzyme: Effects of CPS1 mutations that concentrate in a central domain of unknown function. Mol Genet Metab. 2014;112(2):123–32.

151. Carette JE, Guimaraes CP, Varadarajan M, Park AS, Wuethrich I, Godarova A, et al. Haploid Genetic Screens in Human Cells Identify Host Factors Used by Pathogens. Science. 2009;326(5957):1231-5.

152. Heinz LX, Baumann CL, Koberlin MS, Snijder B, Gawish R, Shui GH, et al. The Lipid-Modifying Enzyme SMPDL3B Negatively Regulates Innate Immunity. Cell Rep. 2015;11(12):1919–28.

153. Gorelik A, Heinz LX, Illes K, Superti-Furga G, Nagar B. Crystal Structure of the Acid Sphingomyelinase-like Phosphodiesterase SMPDL3B Provides Insights into Determinants of Substrate Specificity. J Biol Chem. 2016;291(46):24054–64.

154. Ni MJ, MacFarlane AW, Toft M, Lowell CA, Campbell KS, Hamerman JA. B-cell adaptor for PI3K (BCAP) negatively regulates Toll-like receptor signaling through activation of PI3K. Proc Natl Acad Sci U S A. 2012;109(1):267–72.

155. Wu TD, Watanabe CK. GMAP: a genomic mapping and alignment program for mRNA and EST sequences. Bioinformatics. 2005;21(9):1859–75.

156. Salmela L, Rivals E. LoRDEC: accurate and efficient long read error correction. Bioinformatics. 2014;30(24):3506–14.

157. Fu L, Niu B, Zhu Z, Wu S, Li W. CD-HIT: accelerated for clustering the next-generation sequencing data. Bioinformatics. 2012;28(23):3150–2.

158. Altschul SF, Gish W, Miller W, Myers EW, Lipman DJ. Basic local alignment search tool. J Mol Biol. 1990;215(3):403–10.

159. Buchfink B, Xie C, Huson DH. Fast and sensitive protein alignment using DIAMOND. Nat Methods. 2015;12(1):59–60.

160. Mistry J, Finn RD, Eddy SR, Bateman A, Punta M. Challenges in homology search: HMMER3 and convergent evolution of coiled-coil regions. Nucleic Acids Res. 2013;41(12):e121.

161. Shimizu K, Adachi J, Muraoka Y. ANGLE: a sequencing errors resistant program for predicting protein coding regions in unfinished cDNA. J Bioinform Comput Biol. 2006;4(3):649–64.

162. Hu H, Miao YR, Jia LH, Yu QY, Zhang Q, Guo AY. AnimalTFDB 3.0: a comprehensive resource for annotation and prediction of animal transcription factors. Nucleic Acids Res. 2019;47(D1):D33–8.

163. Thiel T, Michalek W, Varshney RK, Graner A. Exploiting EST databases for the development and characterization of gene-derived SSR-markers in barley (Hordeum vulgare L.). Theor Appl Genet. 2003;106(3):411–22.

164. Sun L, Luo H, Bu D, Zhao G, Yu K, Zhang C, et al. Utilizing sequence intrinsic composition to classify protein-coding and long non-coding transcripts. Nucleic Acids Res. 2013;41(17):e166.

165. Kang YJ, Yang DC, Kong L, Hou M, Meng YQ, Wei L, et al. CPC2: a fast and accurate coding potential calculator based on sequence intrinsic features. Nucleic Acids Res. 2017;45(W1):W12–6.

166. Li W, Cowley A, Uludag M, Gur T, McWilliam H, Squizzato S, et al. The EMBL-EBI bioinformatics web and programmatic tools framework. Nucleic Acids Res. 2015;43(W1):W580–4.

167. Li A, Zhang J, Zhou Z. PLEK: a tool for predicting long non-coding RNAs and messenger RNAs based on an improved k-mer scheme. BMC Bioinformatics. 2014;15:311.

168. Langmead B, Salzberg SL. Fast gapped-read alignment with Bowtie 2. Nat Methods. 2012;9(4):357–9.

169. Li B, Dewey CN. RSEM: accurate transcript quantification from RNA-Seq data with or without a reference genome. BMC Bioinformatics. 2011;12(1):1–16.

170. Love MI, Huber W, Anders S. Moderated estimation of fold change and dispersion for RNA-seq data with DESeq2. Genome Biol. 2014;15(12):550.

171. Wickham H. ggplot2. Wiley Interdiscip Rev Comput Stat. 2011;3(2):180–5.

172. Kumar S, Stecher G, Li M, Knyaz C, Tamura K. MEGA X: Molecular Evolutionary Genetics Analysis across Computing Platforms. Mol Biol Evol. 2018;35(6):1547–9.

173. Saitou N, Nei M. The Neighbor-Joining Method: a New Method for Reconstructing Phylogenetic Trees. Mol Biol Evol. 1987;4(4):406–25.

174. Letunic I, Bork P. Interactive Tree Of Life (iTOL) v5: an online tool for phylogenetic tree display and annotation. Nucleic Acids Res. 2021;49(W1):W293–6.

