## Additional file 1: Figures S1-36 for "Full-length transcriptome maps of reef-building coral illuminate the molecular basis of calcification, symbiosis, and circadian genes"

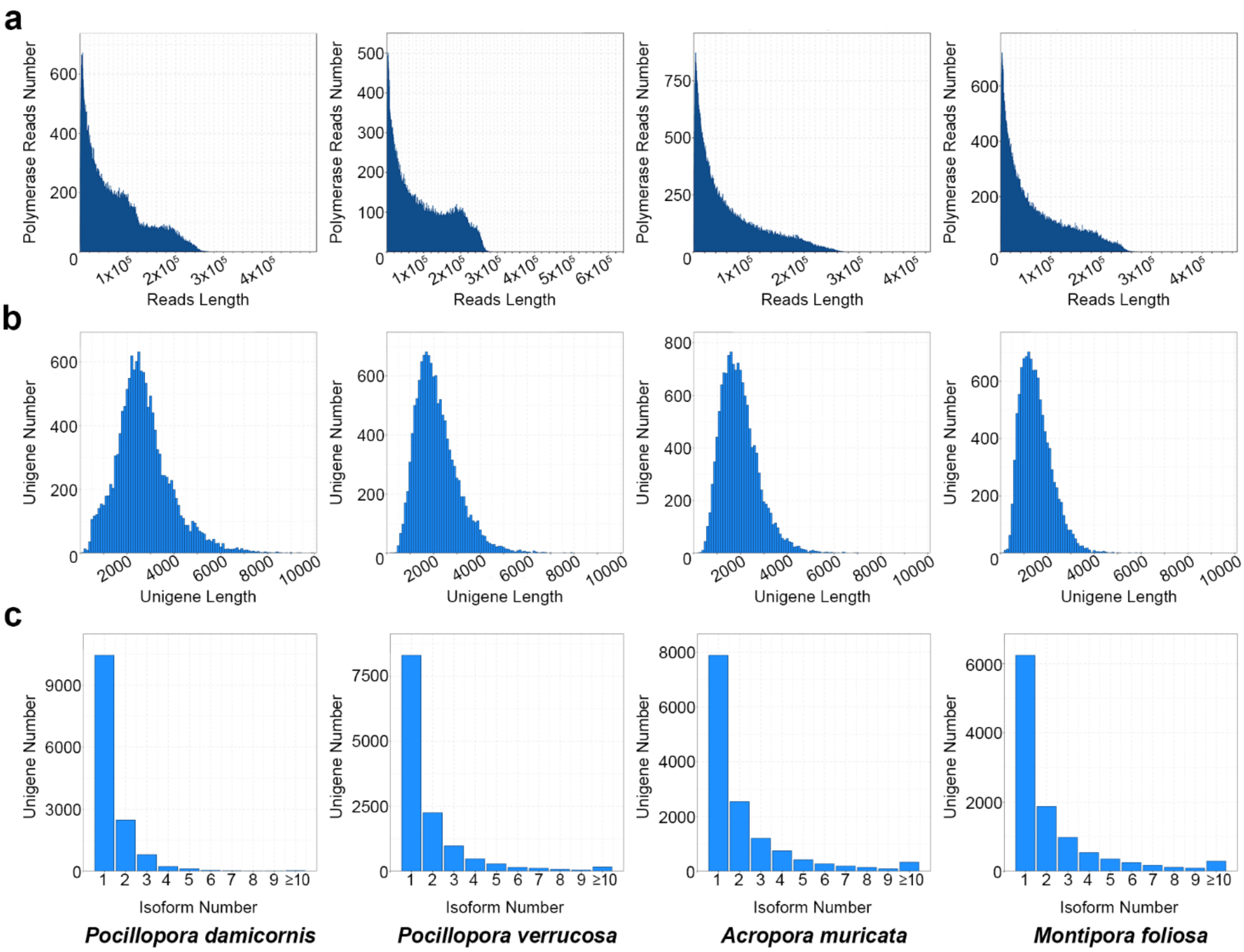


**Fig. S1** Overview of PacBio Sequel II SMRT sequencing data processing. **a.** Polymerase reads length distribution. The horizontal axis represents reads length, the vertical axis represents the number of different read lengths. **b.** Unigene length distribution. The horizontal axis represents unigene length, the vertical axis represents the number of different unigene lengths. **c.** Correspondence between unigene and isoform. The horizontal axis represents isoform number, the vertical axis represents the number of unigenes containing the same number of isoforms.

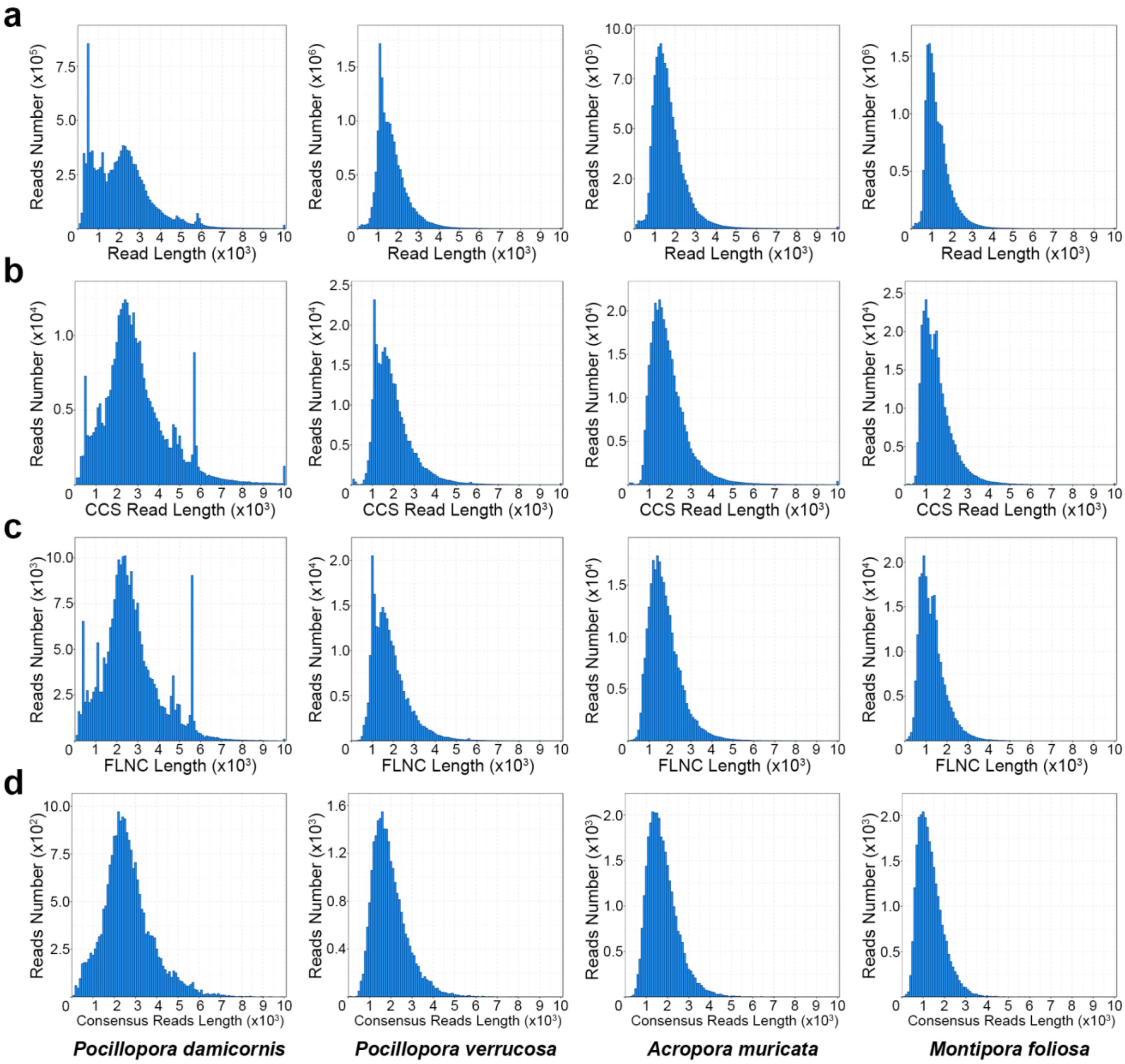


**Fig. S2** Intermediate processes of PacBio Sequel II SMRT sequencing data processing. **a.** Subreads length distribution. The horizontal axis represents reads length, the vertical axis represents the number of different read lengths. **b.** CCS reads length distribution. The horizontal axis represents CCS reads length, the vertical axis represents the number of different read lengths. **c.** Full-Length non-chimericRead (FLNC) length distribution. The horizontal axis represents FLNC length, the vertical axis represents the number of different read lengths. **d.** Polished consensus reads length distribution. The horizontal axis represents reads length, the vertical axis represents the number of different read lengths.

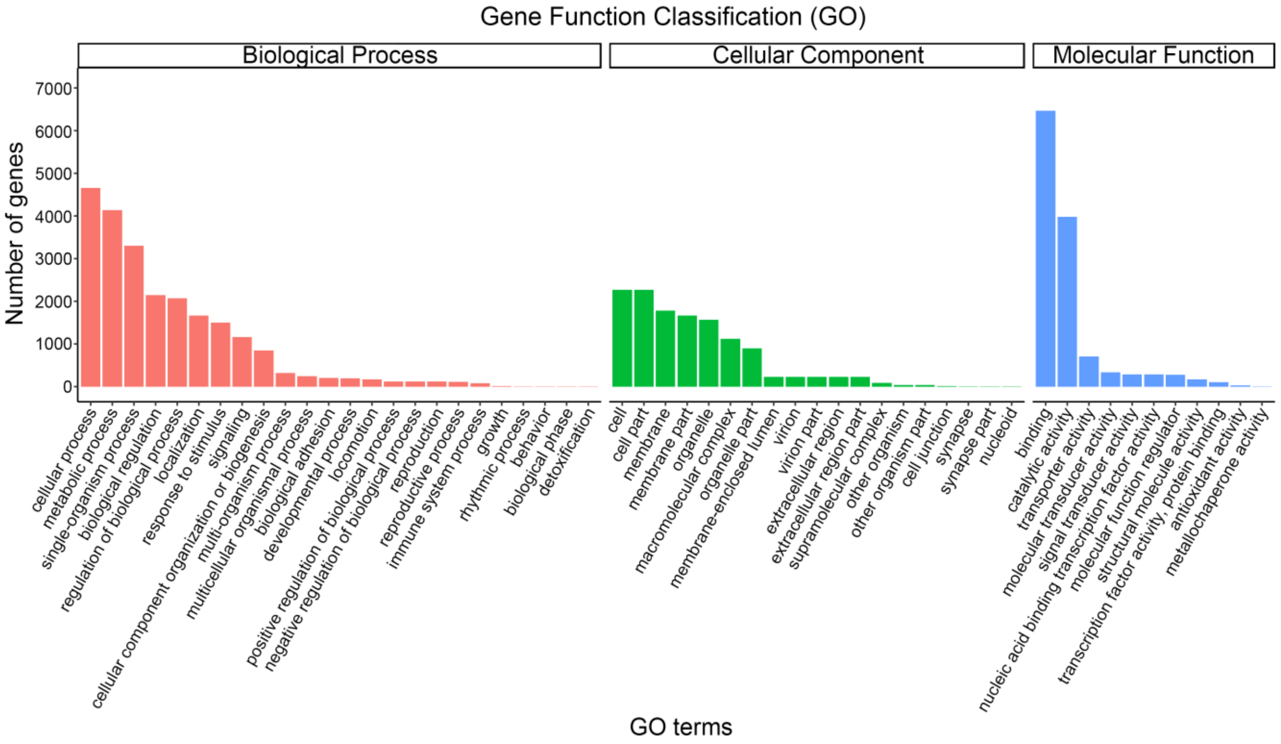


**Fig. S3** GO classifications of coral genes in *P. damicornis*. The horizontal axis represents the GO terms at the next level of the three major GO categories, the vertical axis represents the number of genes annotated to the term (including subterms of the term). Three different categories represent the three basic classifications of Go terms (from left to right, biological processes, cellular components, and molecular functions).

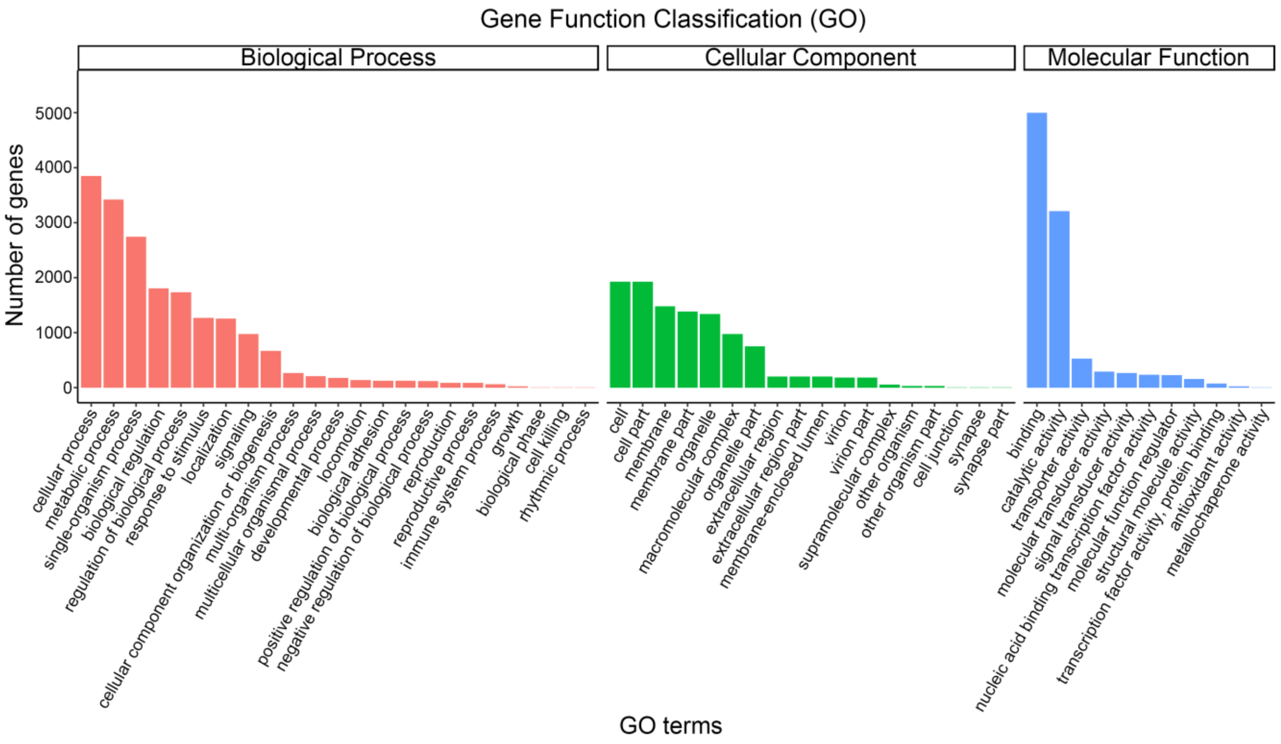


**Fig. S4** GO classifications of coral genes in *P. verrucosa*. The horizontal axis represents the GO terms at the next level of the three major GO categories, the vertical axis represents the number of genes annotated to the term (including subterms of the term). Three different categories represent the three basic classifications of Go terms (from left to right, biological processes, cellular components, and molecular functions).

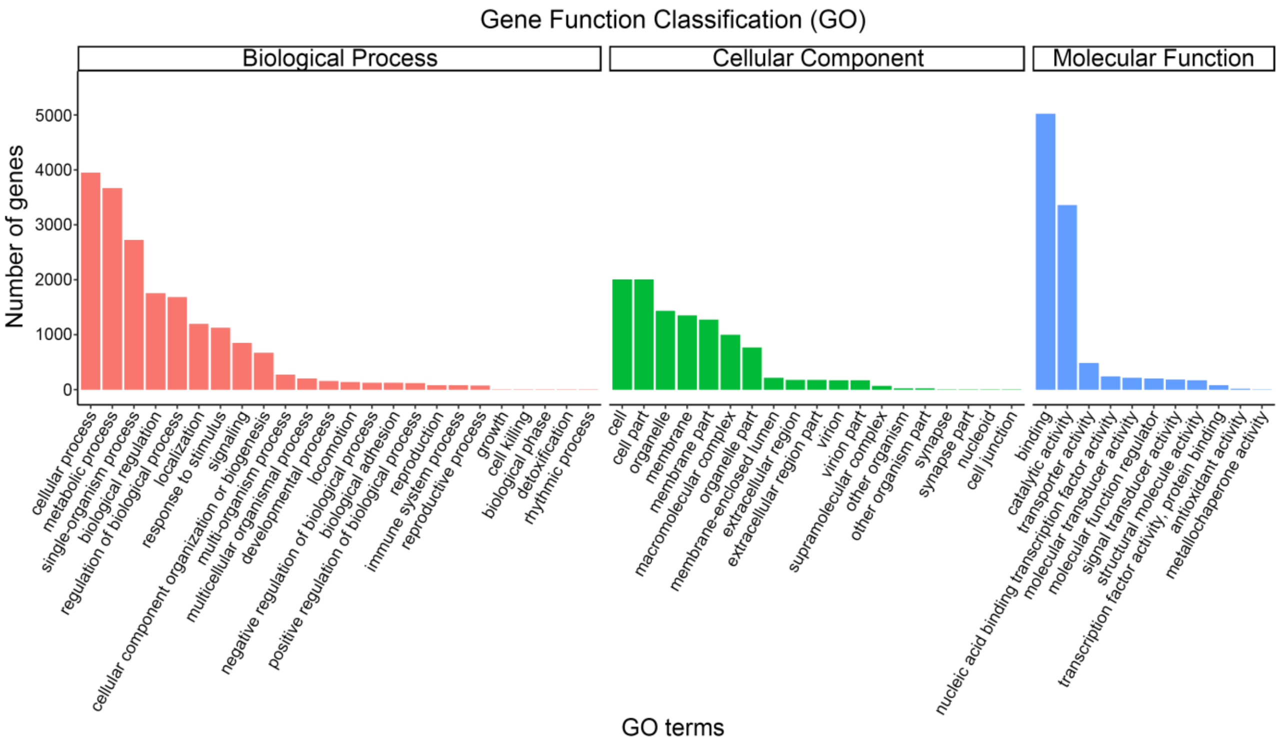


**Fig. S5** GO classifications of coral genes in *A. muricata*. The horizontal axis represents the GO terms at the next level of the three major GO categories, the vertical axis represents the number of genes annotated to the term (including subterms of the term). Three different categories represent the three basic classifications of Go terms (from left to right, biological processes, cellular components, and molecular functions).

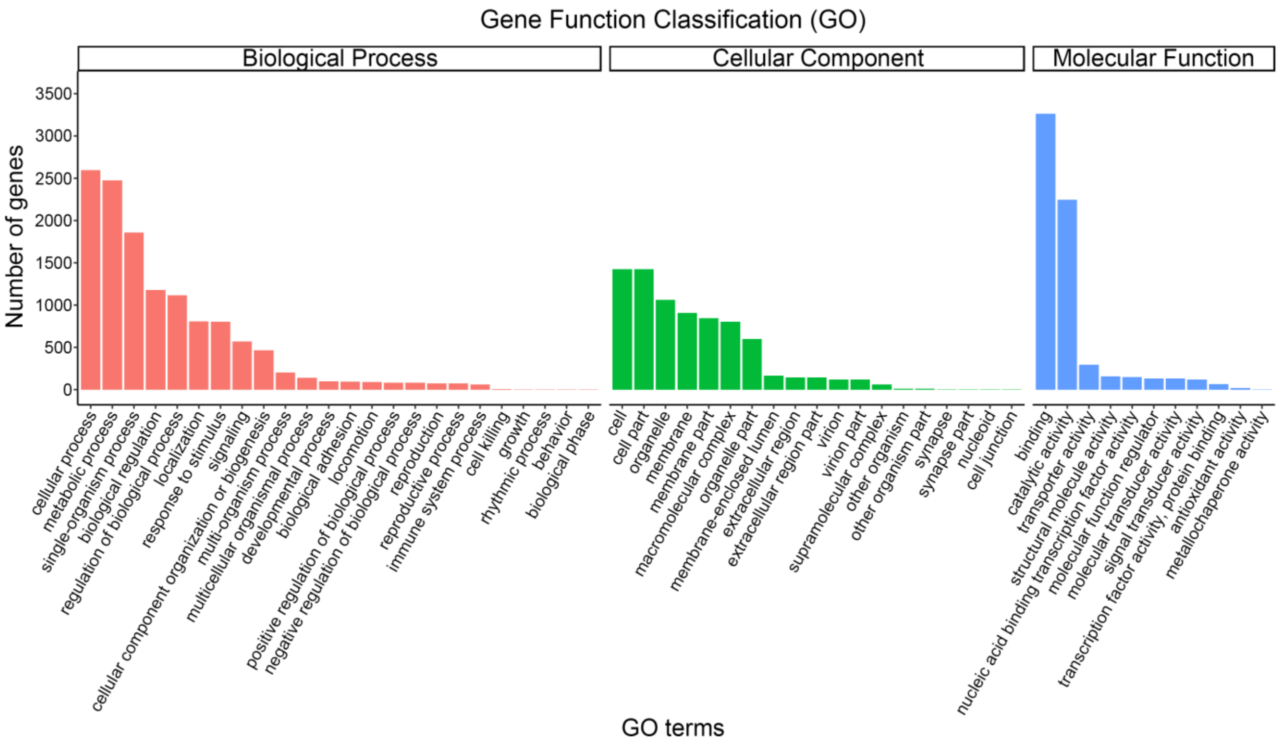


**Fig. S6** GO classifications of coral genes in *M. foliosa*. The horizontal axis represents the GO terms at the next level of the three major GO categories, the vertical axis represents the number of genes annotated to the term (including subterms of the term). Three different categories represent the three basic classifications of Go terms (from left to right, biological processes, cellular components, and molecular functions).

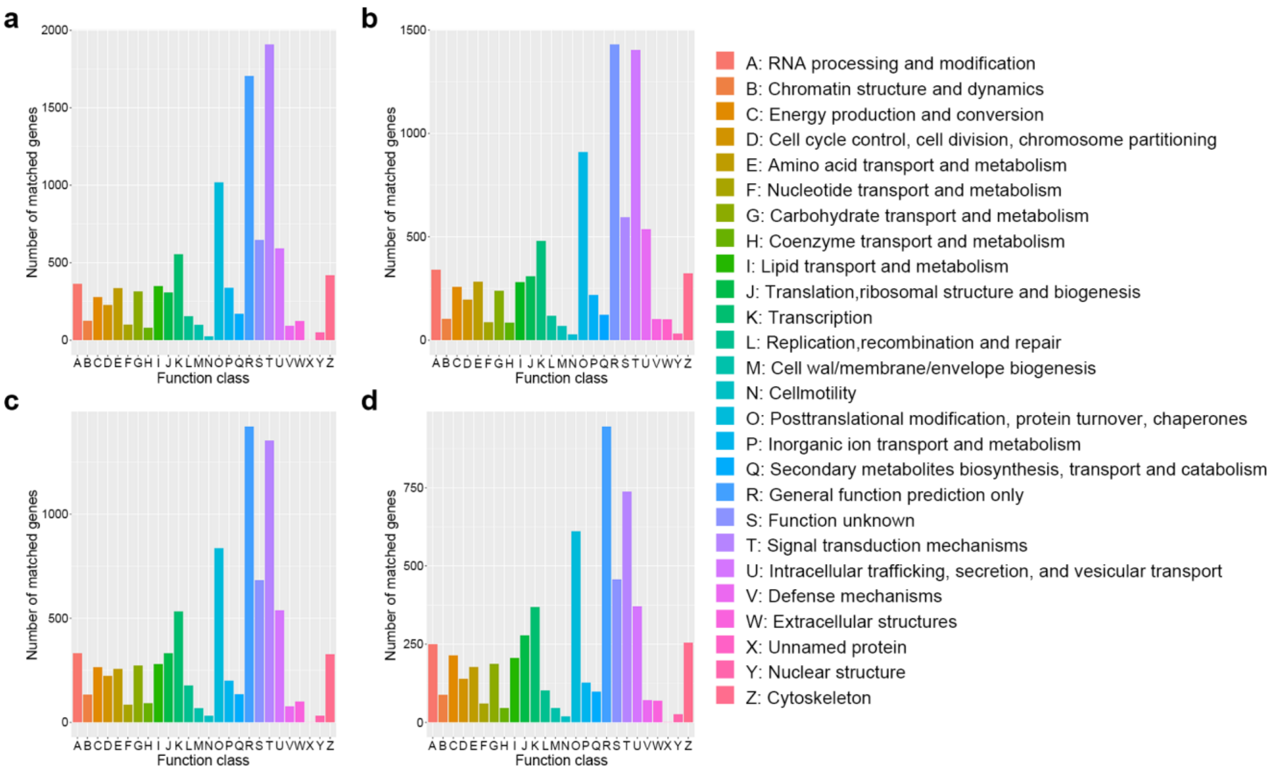


**Fig. S7** KOG classifications of coral genes. **a-d** represent *P. damicornis, P. verrucosa, A. muricata* and *M. foliosa* respectively. The horizontal axis is the name of the 26 function classes of KOG/COG, and the vertical axis is the number of genes matched to different classes.

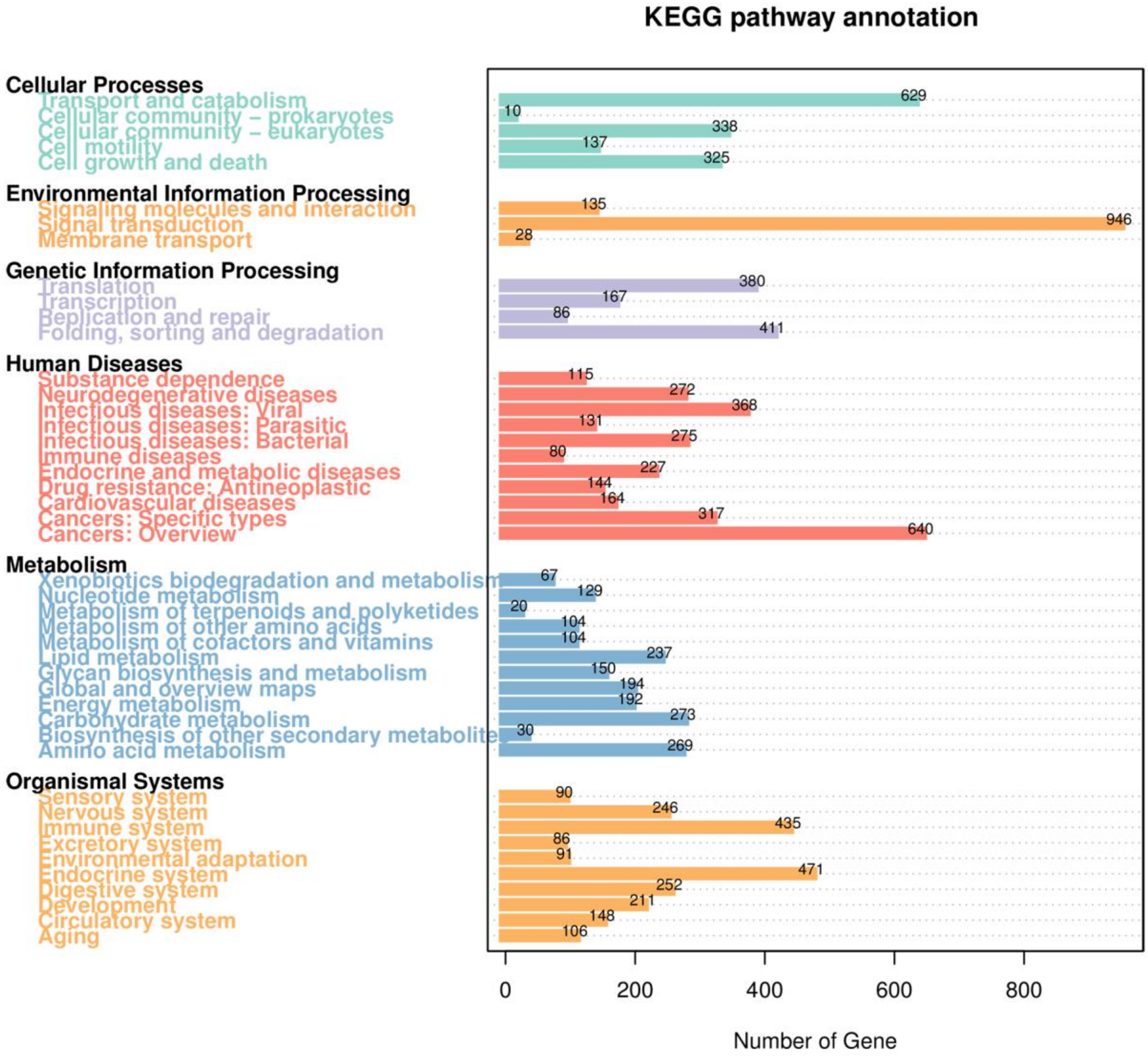


**Fig. S8** KEGG metabolic pathway classifications of coral genes in *P. damicornis*. On the left are the different KEGG categories, including cellular processes, environmental information processing, genetic information processing, human diseases, metabolism and organismal systems. On the right is the number of their corresponding genes.

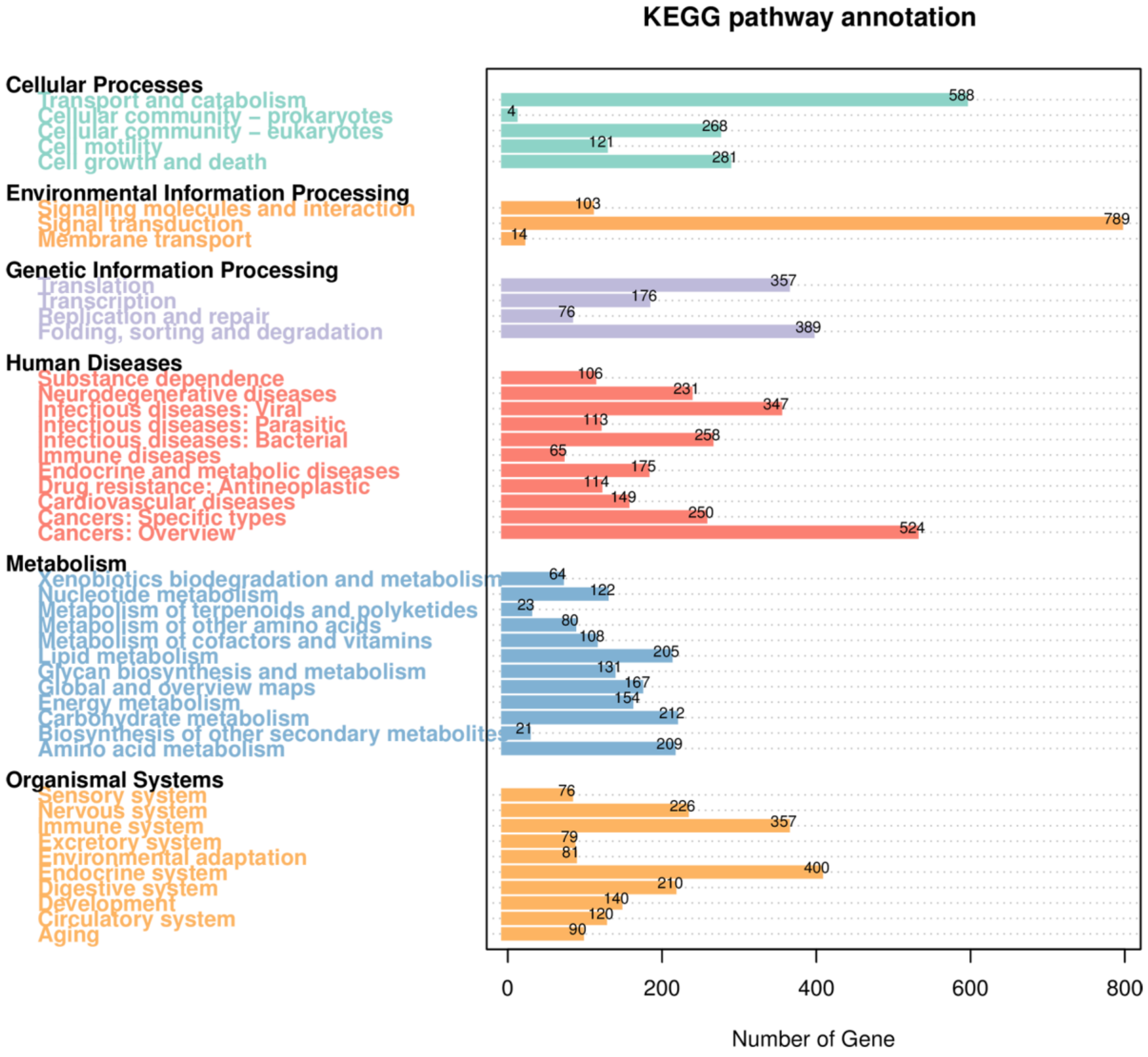


**Fig. S9** KEGG metabolic pathway classifications of coral genes in *P. verrucosa*. On the left are the different KEGG categories, including cellular processes, environmental information processing, genetic information processing, human diseases, metabolism and organismal systems. On the right is the number of their corresponding genes.

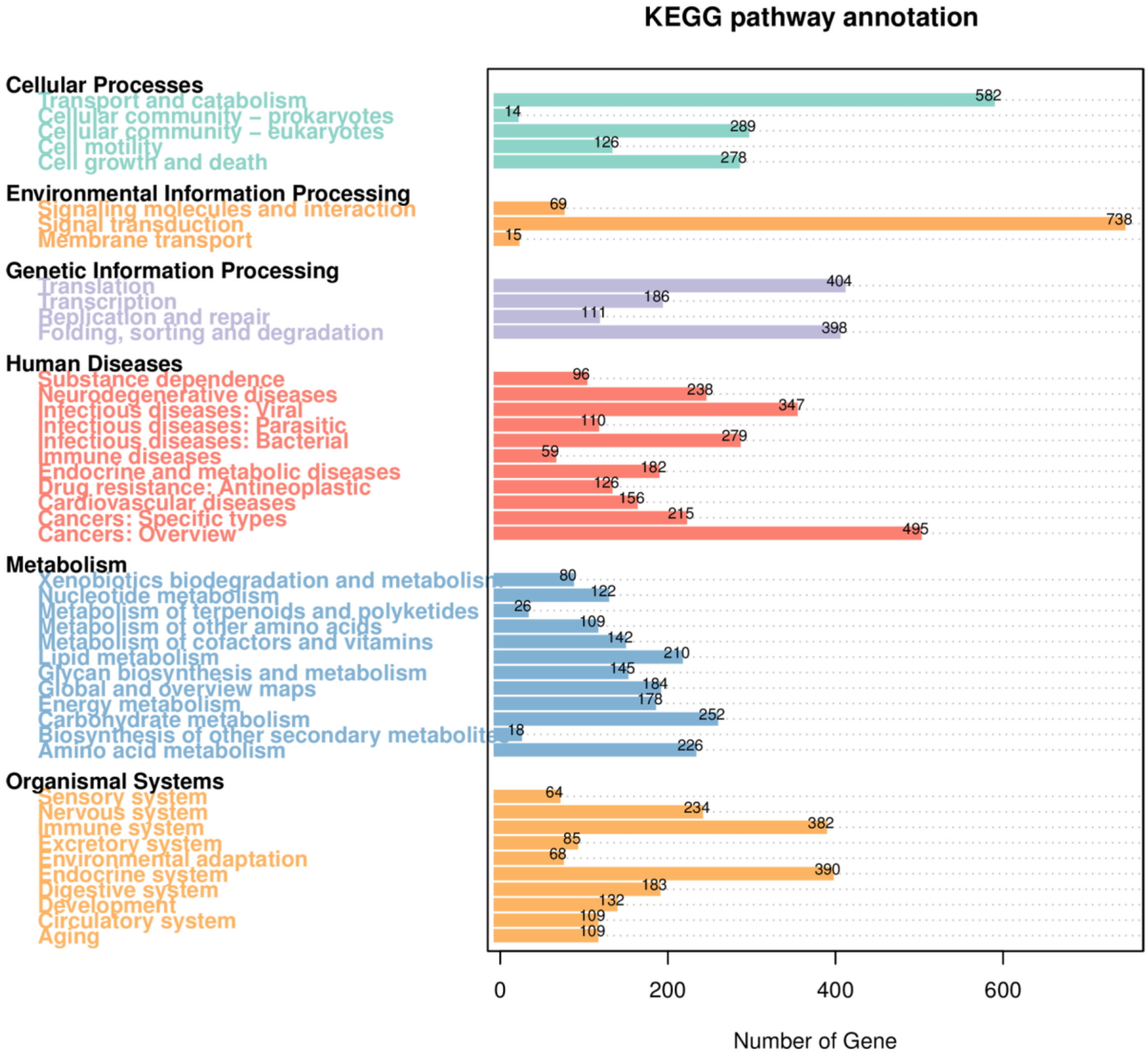


**Fig. S10** KEGG metabolic pathway classifications of coral genes in *A. muricata*. On the left are the different KEGG categories, including cellular processes, environmental information processing, genetic information processing, human diseases, metabolism and organismal systems. On the right is the number of their corresponding genes.

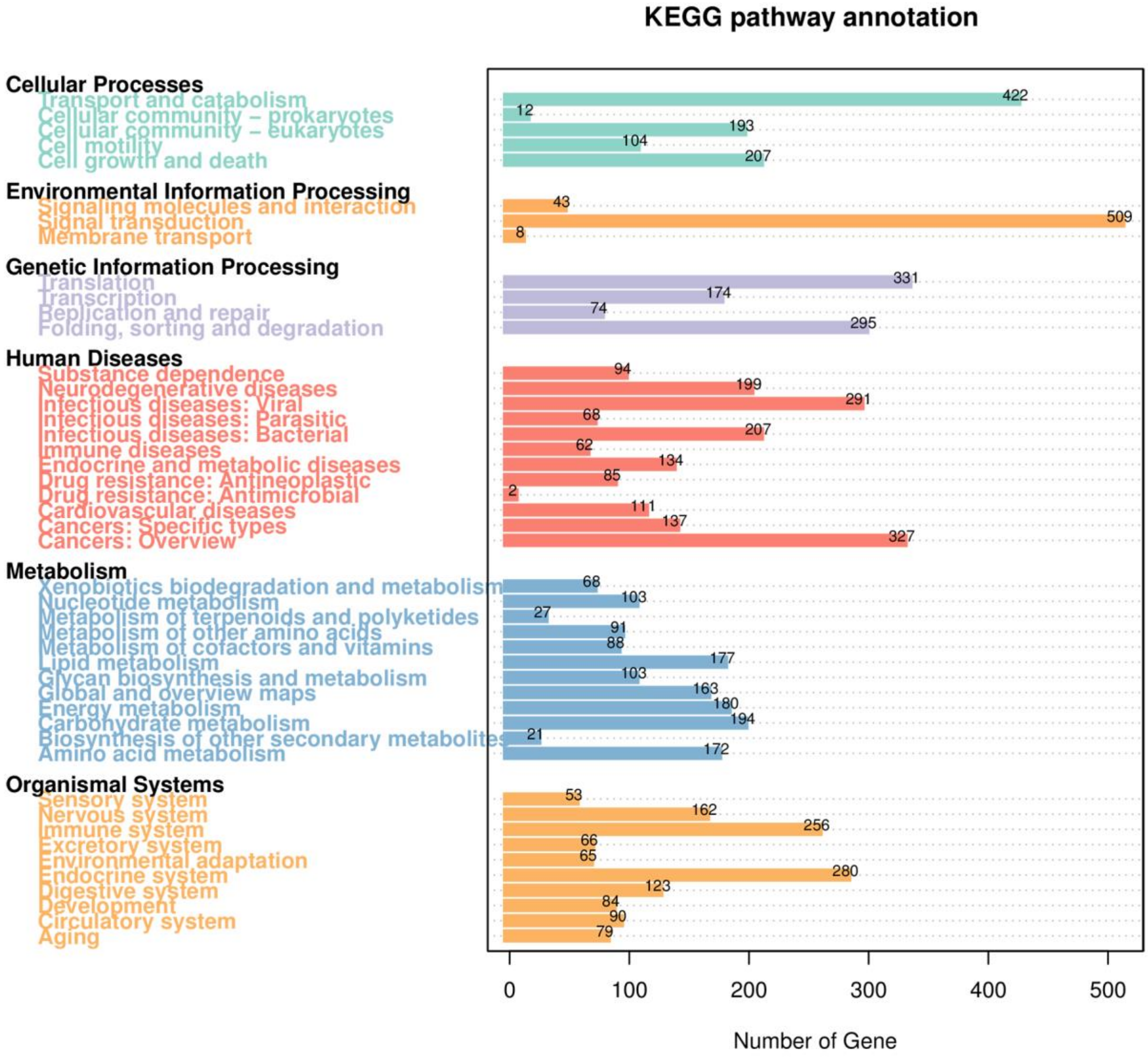


**Fig. S11** KEGG metabolic pathway classifications of coral genes in *M. foliosa*. On the left are the different KEGG categories, including cellular processes, environmental information processing, genetic information processing, human diseases, metabolism and organismal systems. On the right is the number of their corresponding genes.

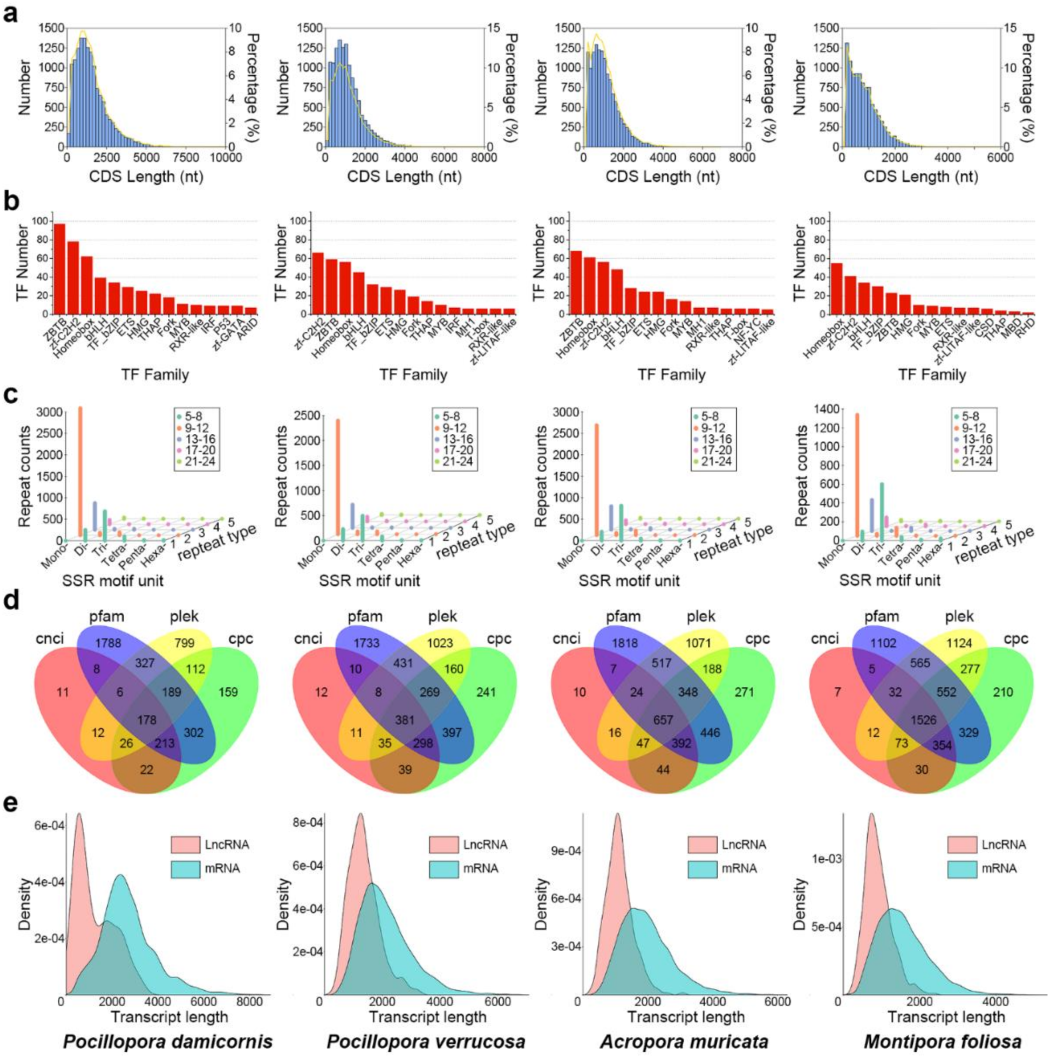


**Fig. S12** Summary of coral gene structural analysis. **a.** CDS length distribution. The horizontal axis represents the length of the predicted CDS, the vertical axis represents the number (blue bar chart) and percentage (yellow curve chart) of transcripts of the CDS. **b**. Predicted TF family. The horizontal axis represents the top 15 predicted transcription factor families in coral, and the vertical axis represents the number of them. **c**. SSR motif distribution. The X coordinate (SSR motif unit) is the SSR type, which refers to the number of repeating bases, the Y coordinate (repeat type) is the number of base (SSR motif unit) repeats, using different colors mean different repeat number intervals (the details are seen in the legend), the Z coordinate (Repeat counts) is the number of SSRs. **d**. Venn plots of predicted lncRNA. The sum of the numbers in each large circle represents the number of lncRNA predicted by one of CNCI, PLEK, CPC2 and Pfam databases, and the overlapping circles indicate the number of lncRNA predicted by these two or more databases simultaneously. **e**. LncRNA and mRNA length distribution comparison. The horizontal axis is the length of transcripts, the vertical axis is their density of distribution.

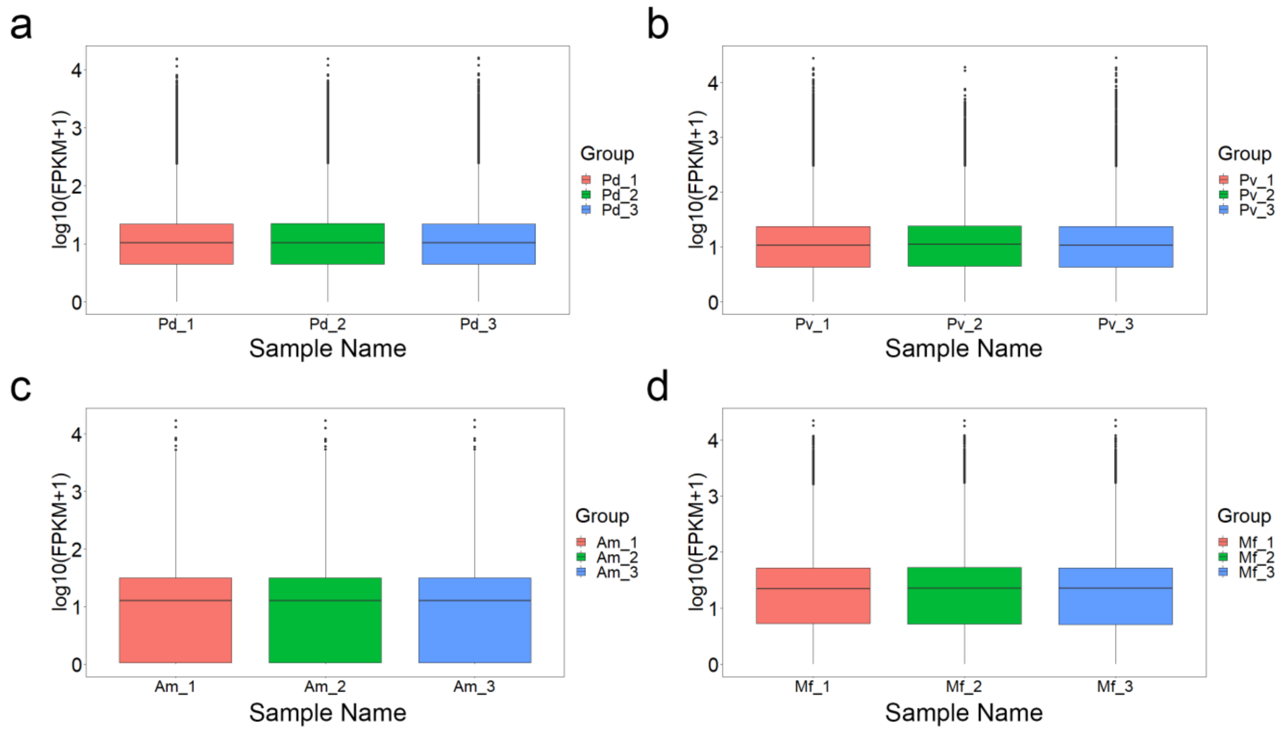


**Fig. S13** FPKM box plot of each coral gene expression. The horizontal axis is the sample name, *P. damicornis* (**a**), *P. verrucosa* (**b**), *A. muricata* (**c**) and *M. foliosa* (**d**), the vertical axis is log10 (FPKM+1). Each box plot shows five statistics results, including maximum, upper quartile, median, lower quartile and minimum from top to down.

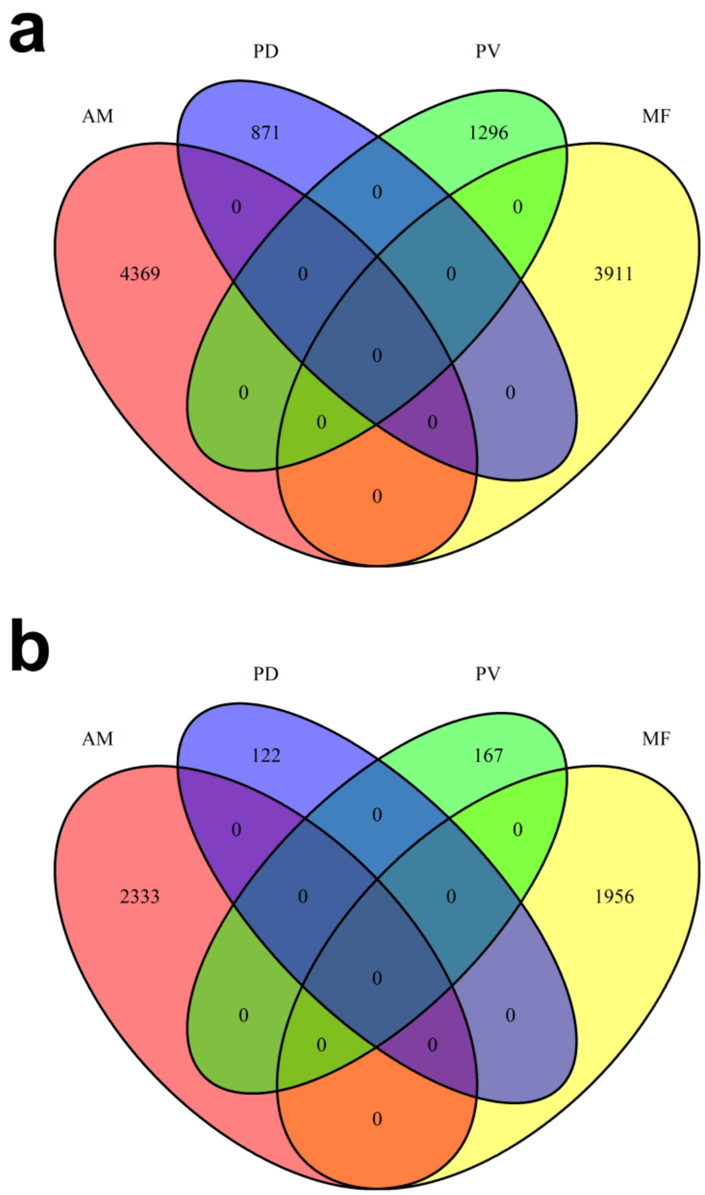


**Fig. S14** Venn plots of coral differentially expressed genes. **a.** Similarities and differences of four coral differentially expressed genes in the case of padj<0.001 and |log2(FoldChange)|>=2. **b.** Similarities and differences of four coral differentially expressed genes in the case of padj<0.001 and |log2(FoldChange)|>=10. Numbers represent the number of differential genes. PD: *P. damicornis*; PV: *P. verrucosa*; AM: *A. muricata*; and MF: *M. foliosa*.

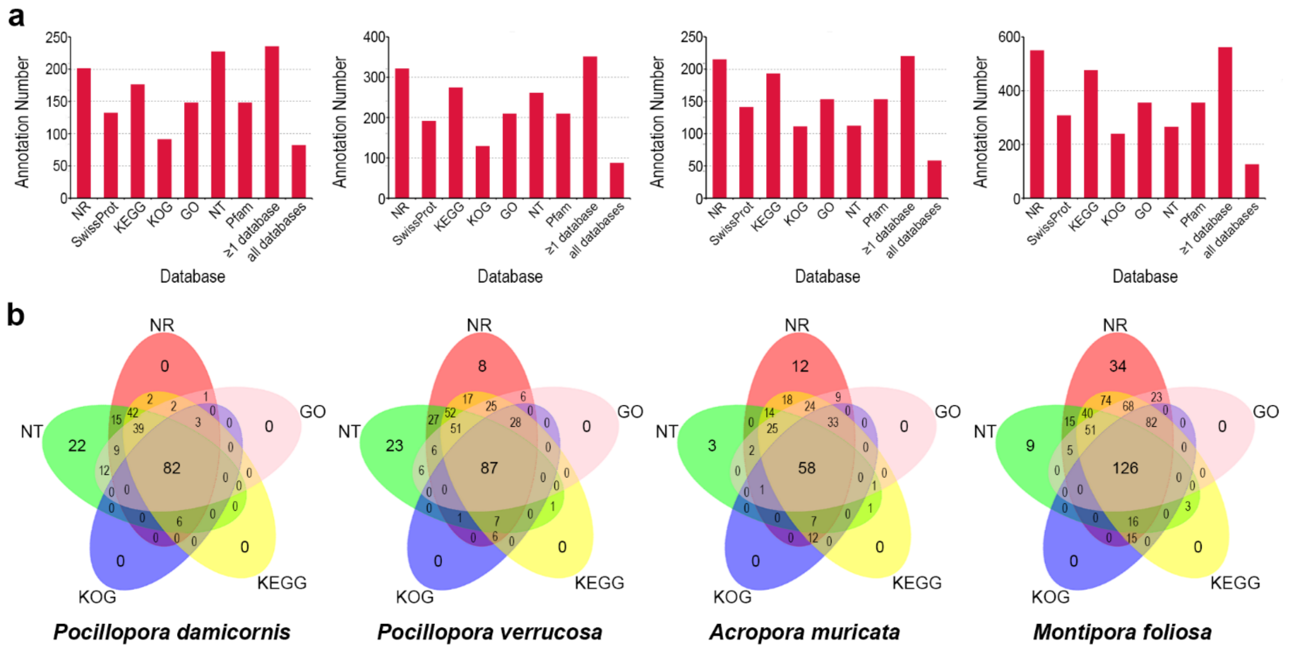


**Fig. S15** Summary of Symbiodiniaceae gene functional annotation. **a.** Statistics of the annotation results of Symbiodiniaceae in four reef-building corals, in the NR, Swiss-Prot, KEGG, KOG, GO NT and Pfam databases. The horizontal axis represents the different functional databases, and the vertical axis represents the number of sequences annotated in different functional databases, at least one database and all databases. **b.** Venn plots of the number of annotated sequences of Symbiodiniaceae in four reef-building corals obtained using the NR, KEGG, KOG, GO and NT databases. The sum of the numbers in each large circle represents the number of transcripts annotated in one database, and the overlapping circles indicate the number of transcripts annotated to these databases simultaneously.

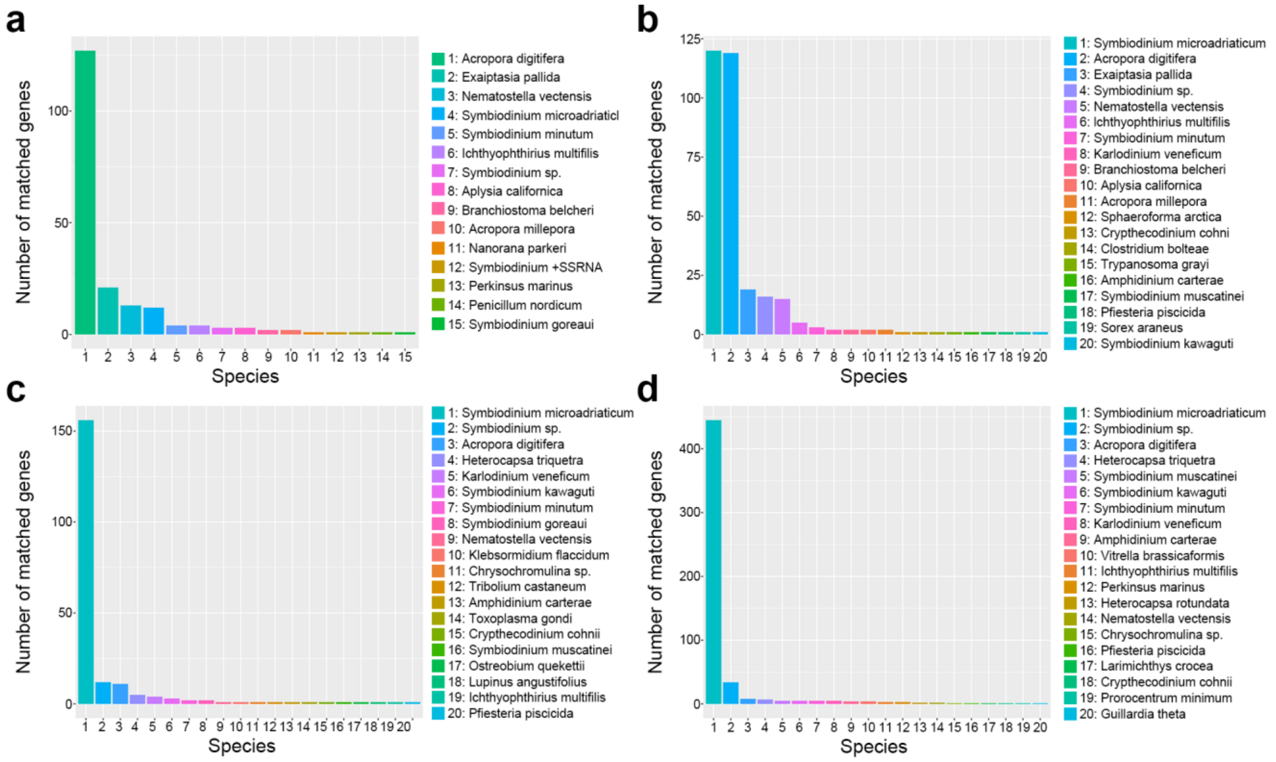


**Fig. S16** NR database annotation. The top 20 species with the greatest number of top sequence hits to Symbiodiniaceae sequences of *P. damicornis* (**a**), *P. verrucosa* (**b**), *A. muricata* (**c**) and *M. foliosa* (**d**) are shown. The horizontal axis represents the species ID, and the vertical axis represents the number of Symbiodiniaceae unigenes annotated to different species.

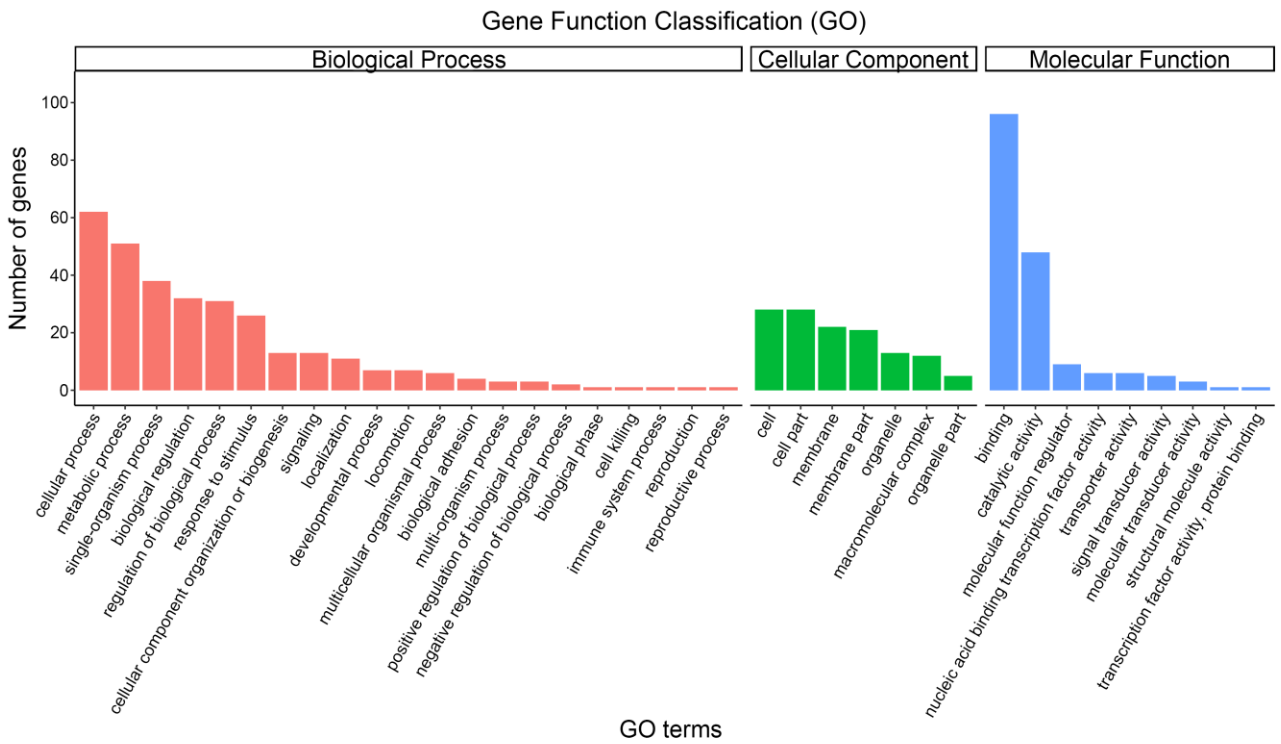


**Fig. S17** GO classifications of Symbiodiniaceae genes in *P. damicornis*. The horizontal axis represents the GO terms at the next level of the three major GO categories, the vertical axis represents the number of genes annotated to the term (including subterms of the term). Three different categories represent the three basic classifications of Go terms (from left to right, biological processes, cellular components, and molecular functions).

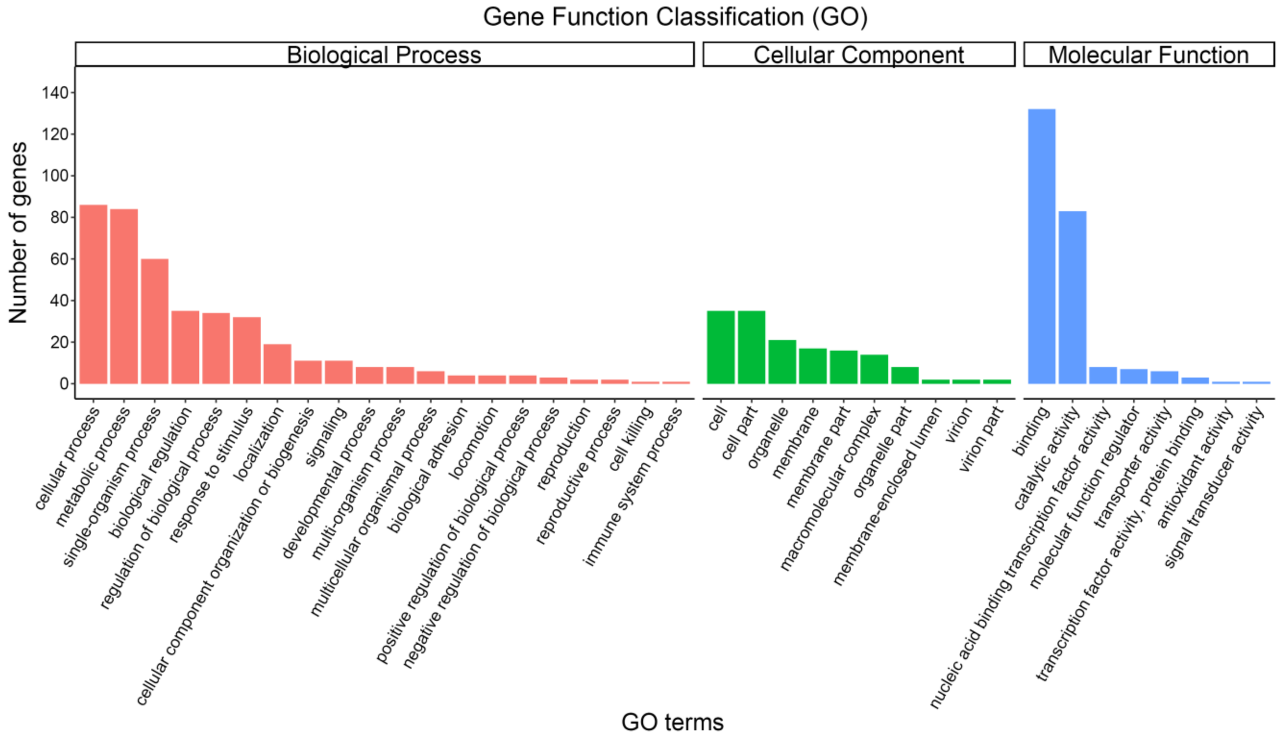


**Fig. S18** GO classifications of Symbiodiniaceae genes in *P. verrucosa*. The horizontal axis represents the GO terms at the next level of the three major GO categories, the vertical axis represents the number of genes annotated to the term (including subterms of the term). Three different categories represent the three basic classifications of Go terms (from left to right, biological processes, cellular components, and molecular functions).

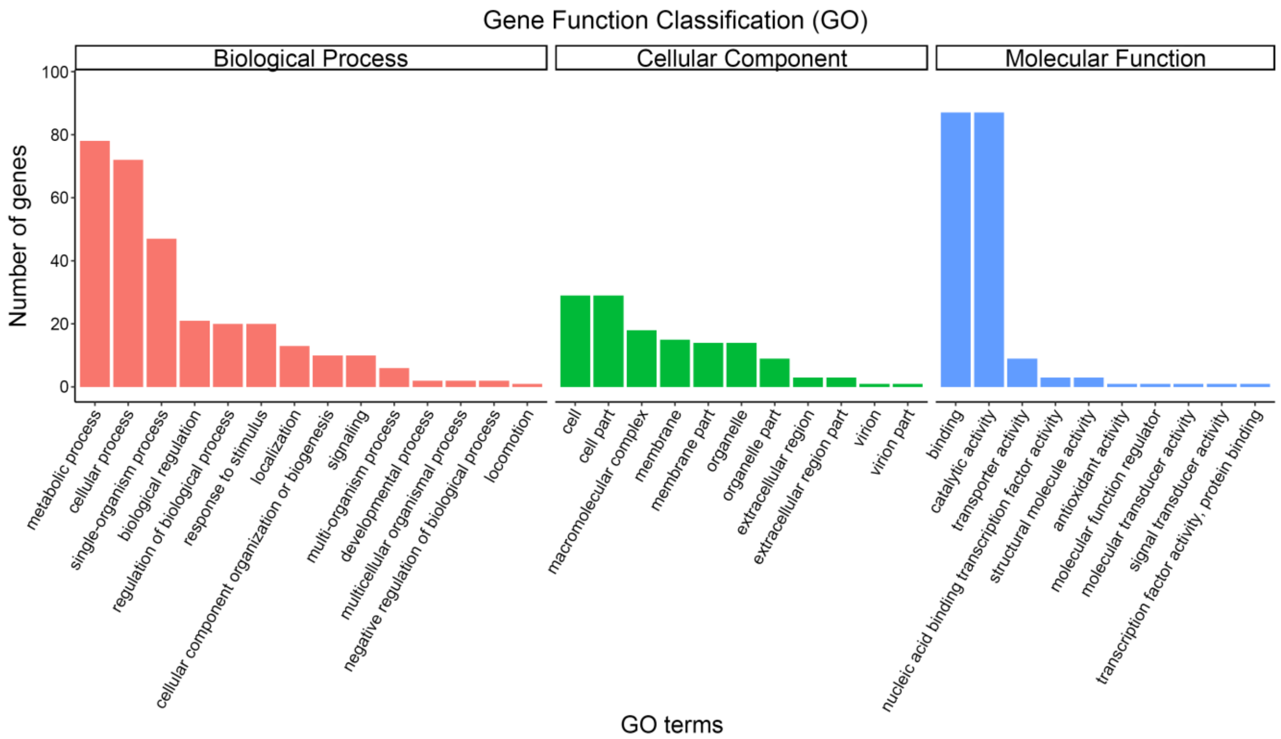


**Fig. S19** GO classifications of Symbiodiniaceae genes in *A. muricata*. The horizontal axis represents the GO terms at the next level of the three major GO categories, the vertical axis represents the number of genes annotated to the term (including subterms of the term). Three different categories represent the three basic classifications of Go terms (from left to right, biological processes, cellular components, and molecular functions).

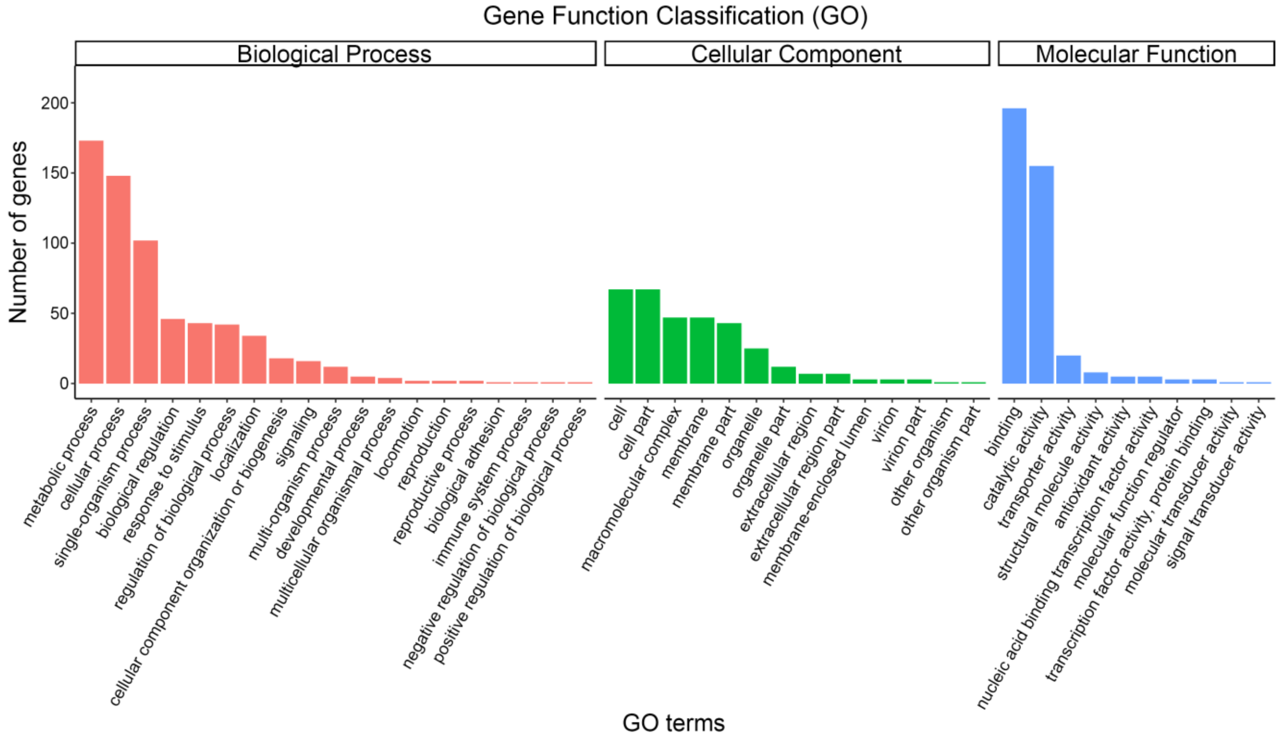


**Fig. S20** GO classifications of Symbiodiniaceae genes in *M. foliosa*. The horizontal axis represents the GO terms at the next level of the three major GO categories, the vertical axis represents the number of genes annotated to the term (including subterms of the term). Three different categories represent the three basic classifications of Go terms (from left to right, biological processes, cellular components, and molecular functions).

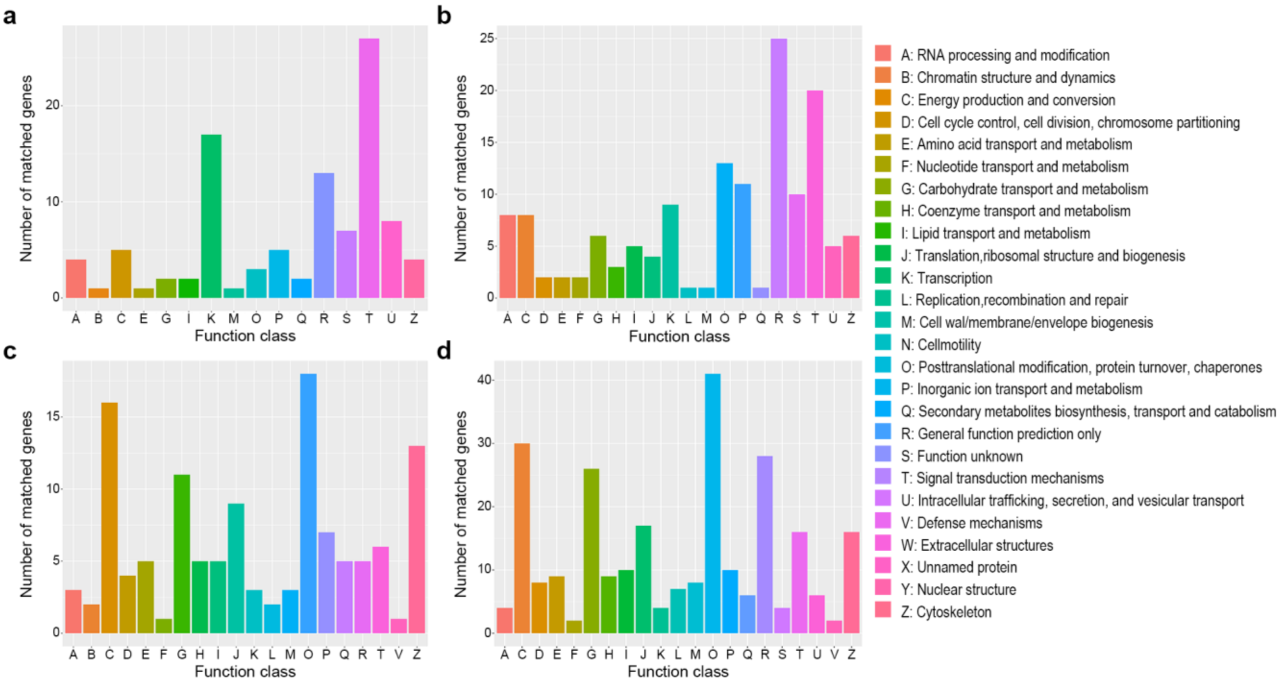


**Fig. S21** KOG classifications of Symbiodiniaceae genes. **a-d** represent *P. damicornis, P. verrucosa, A. muricata* and *M. foliosa* respectively. The horizontal axis is the name of the 26 function classes of KOG/COG, and the vertical axis is the number of genes matched to different classes.

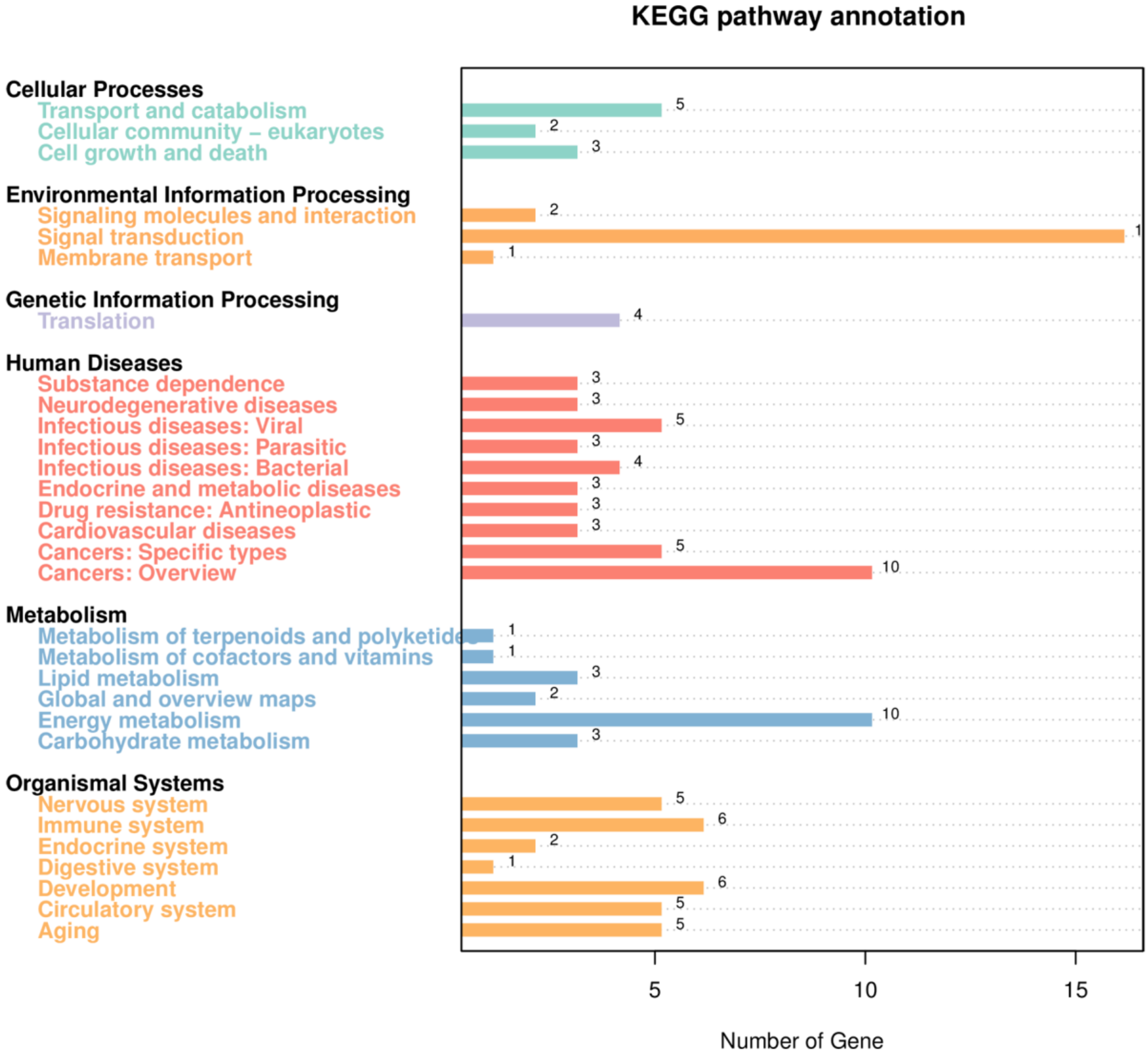


**Fig. S22** KEGG metabolic pathway classifications of Symbiodiniaceae genes in *P. damicornis*. On the left are the different KEGG categories, including cellular processes, environmental information processing, genetic information processing, human diseases, metabolism and organismal systems. On the right is the number of their corresponding genes.

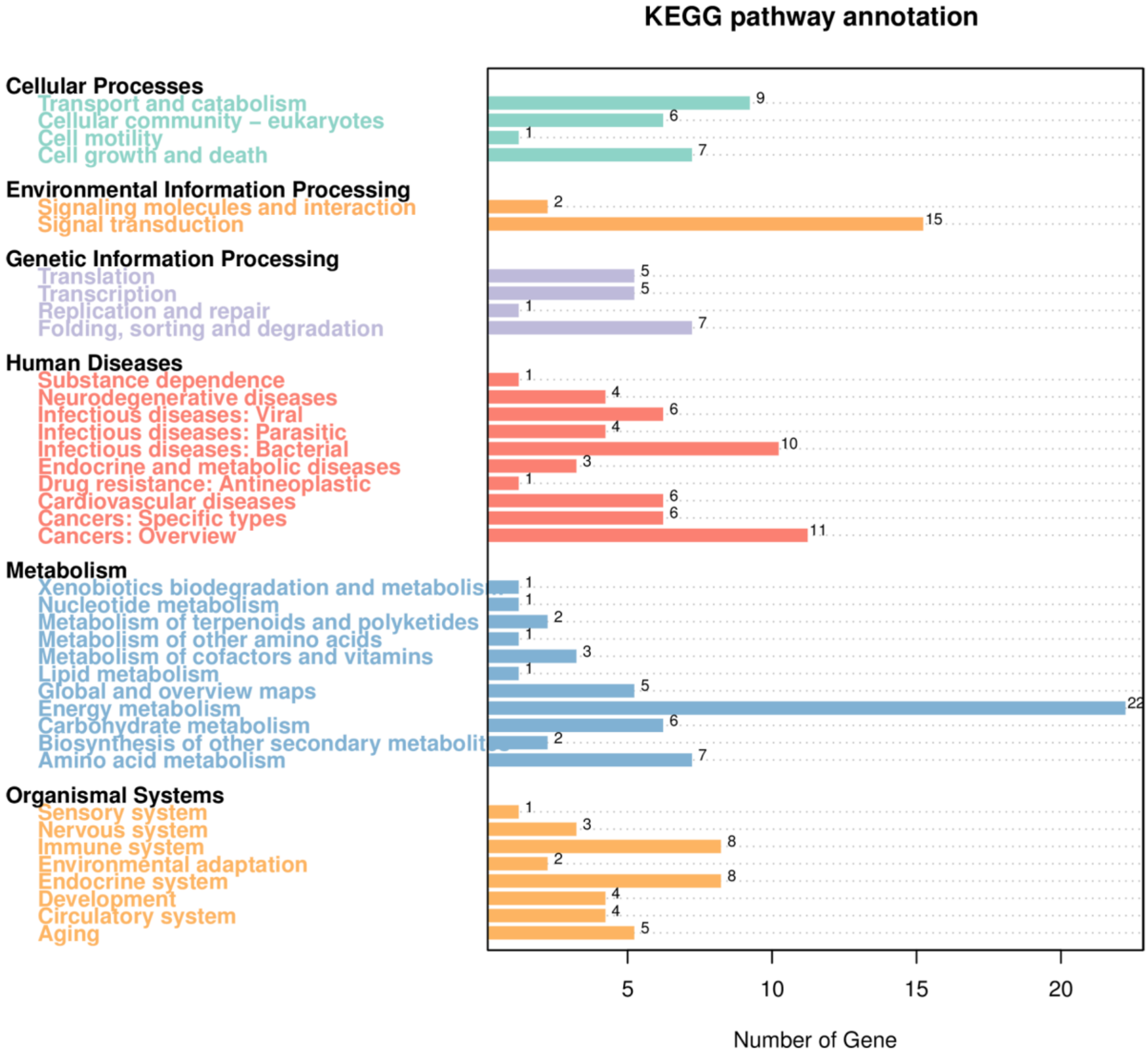


**Fig. S23** KEGG metabolic pathway classifications of Symbiodiniaceae genes in *P. verrucosa*. On the left are the different KEGG categories, including cellular processes, environmental information processing, genetic information processing, human diseases, metabolism and organismal systems. On the right is the number of their corresponding genes.

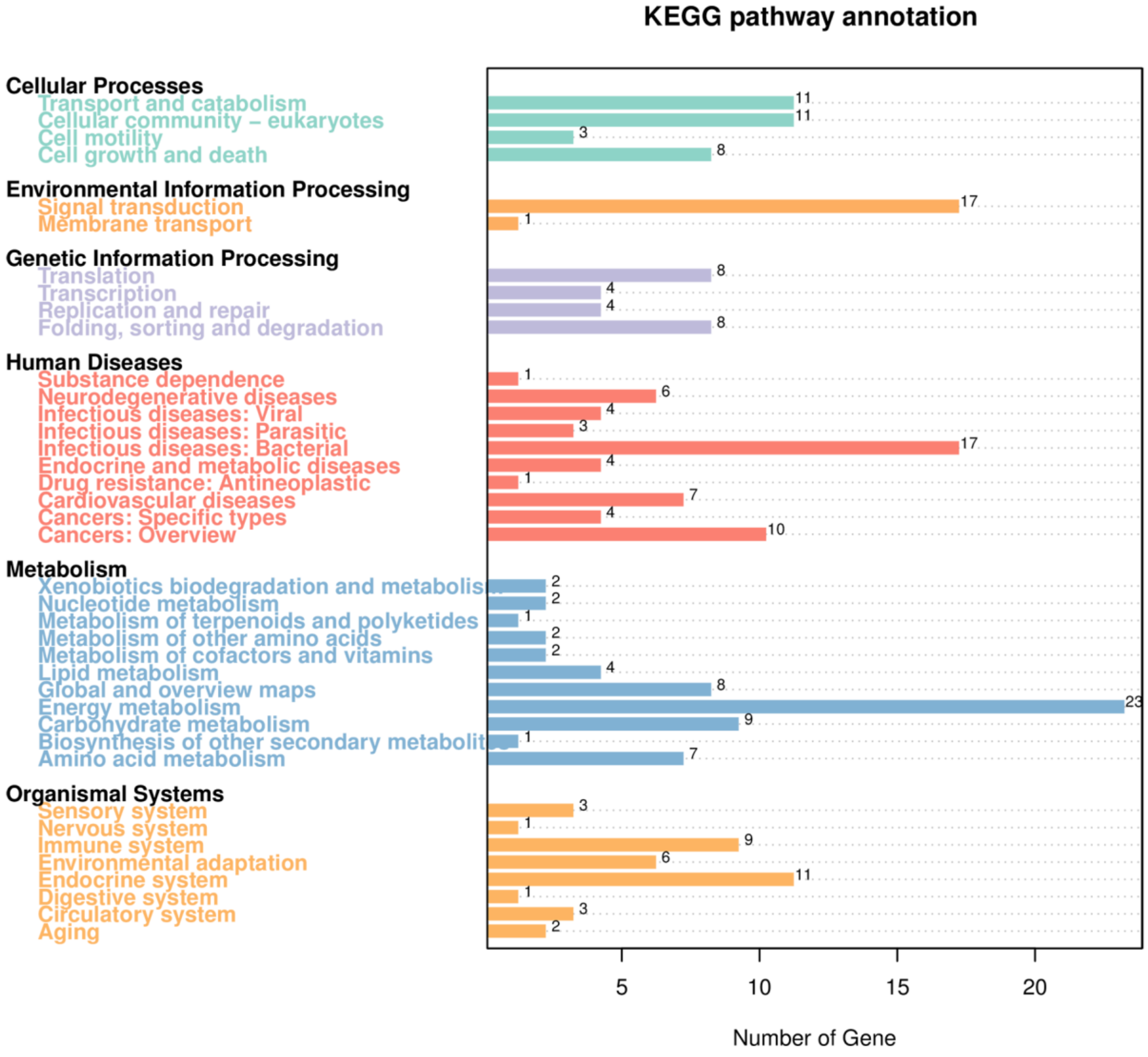


**Fig. S24** KEGG metabolic pathway classifications of Symbiodiniaceae genes in *A. muricata*. On the left are the different KEGG categories, including cellular processes, environmental information processing, genetic information processing, human diseases, metabolism and organismal systems. On the right is the number of their corresponding genes.

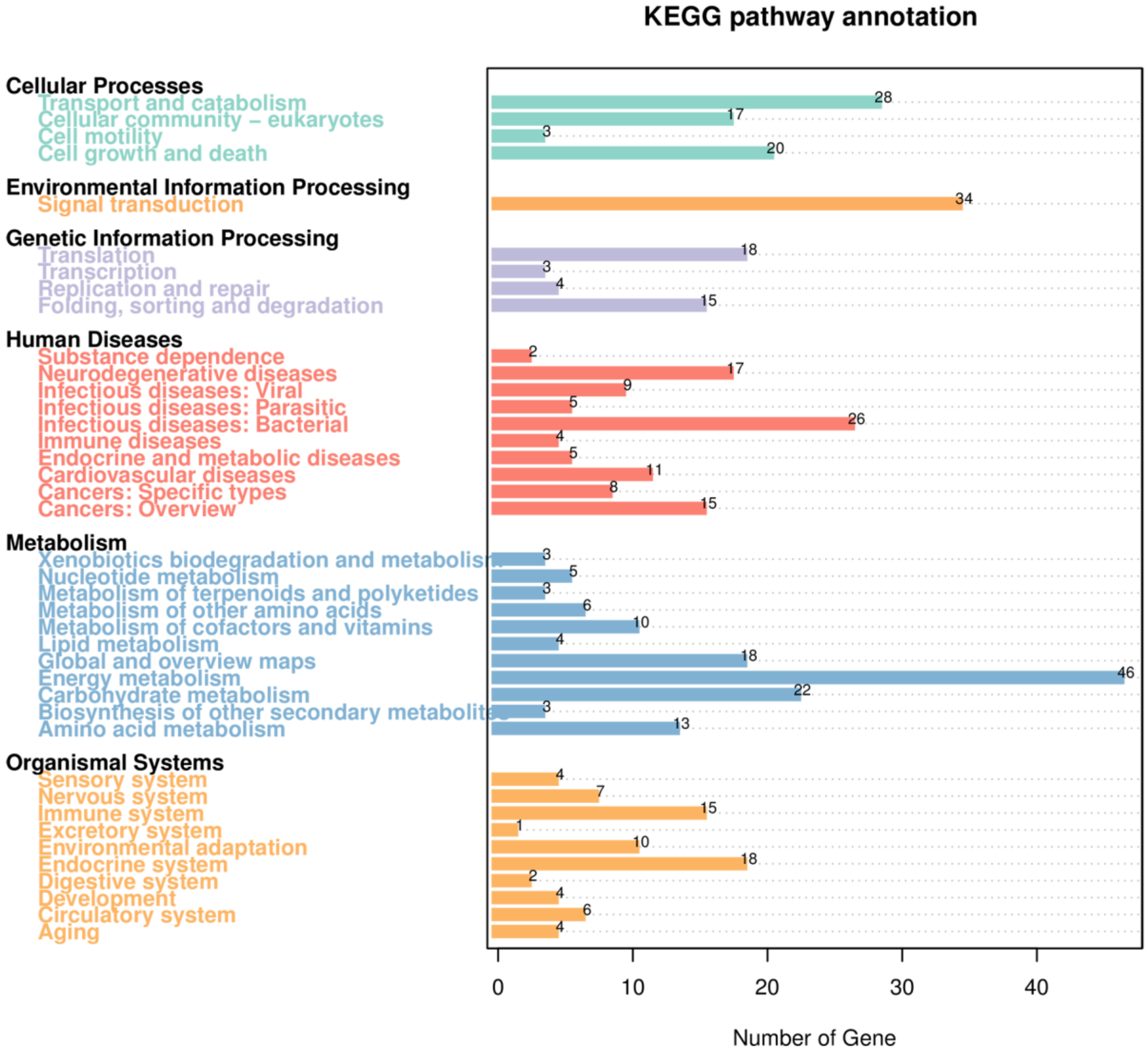


**Fig. S25** KEGG metabolic pathway classifications of Symbiodiniaceae genes in *M. foliosa*. On the left are the different KEGG categories, including cellular processes, environmental information processing, genetic information processing, human diseases, metabolism and organismal systems. On the right is the number of their corresponding genes.

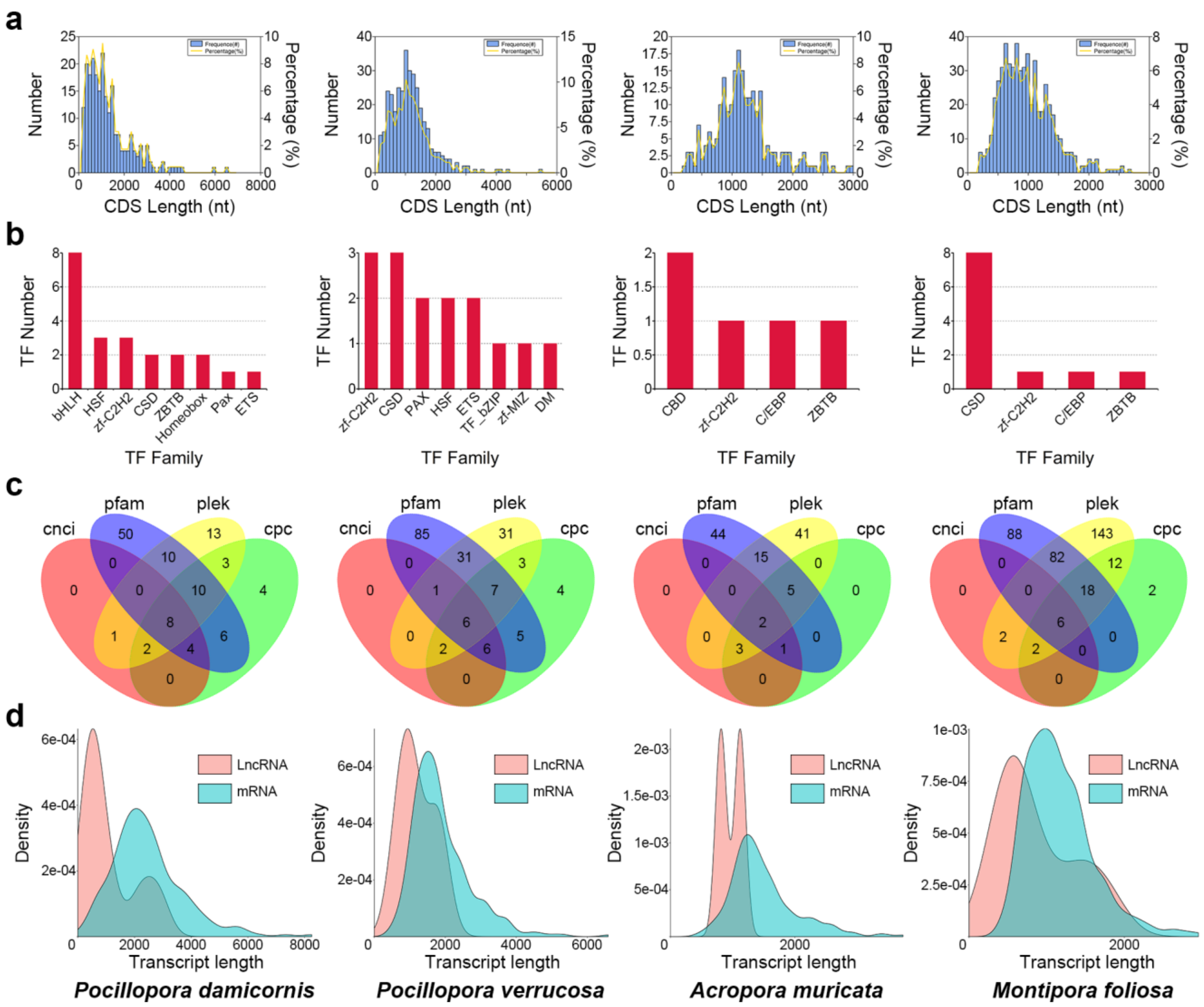


**Fig. S26** Summary of Symbiodiniaceae gene structural analysis. **a.** CDS length distribution. The horizontal axis represents the length of the predicted CDS, the vertical axis represents the number (blue bar chart) and percentage (yellow curve chart) of transcripts of the CDS. **b**. Predicted TF family. The horizontal axis represents the top 8 or 4 predicted transcription factor families, and the vertical axis represents the number of them. **c**. Venn plots of predicted lncRNA. The sum of the numbers in each large circle represents the number of lncRNA predicted by one of CNCI, PLEK, CPC2 and Pfam databases, and the overlapping circles indicate the number of lncRNA predicted by these two or more databases simultaneously. **d**. LncRNA and mRNA length distribution comparison. The horizontal axis is the length of transcripts, the vertical axis is their density of distribution.

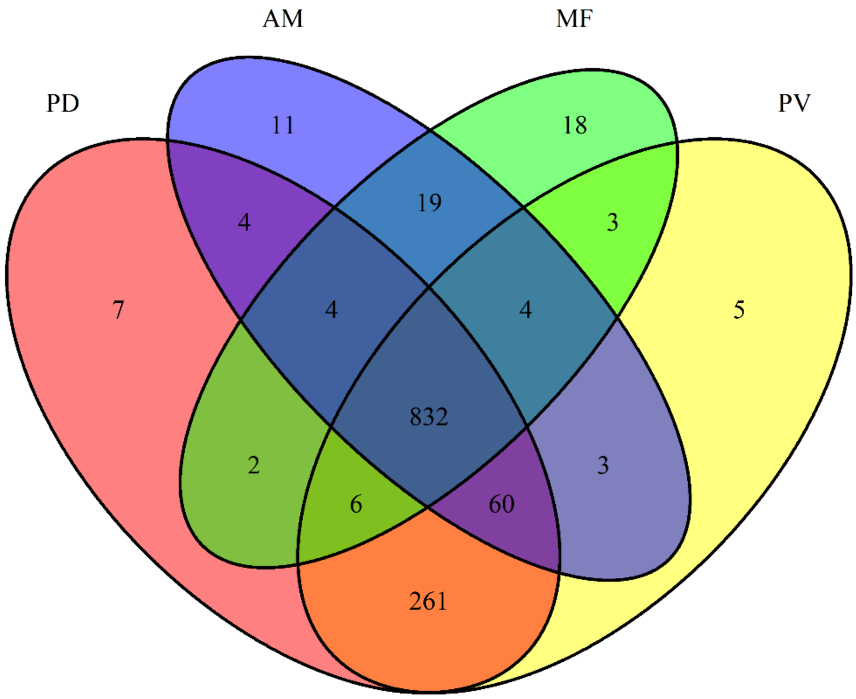


**Fig. S27** Venn plots of statistics on the number of Symbiodiniaceae sequences expressed in four reef-building corals. The sum of the numbers in each large circle represents the number of Symbiodiniaceae sequences expressed in one of the corals, and the overlapping circles indicate the number of Symbiodiniaceae sequences expressed in these two or more corals simultaneously. PD: *P. damicornis*; PV: *P. verrucosa*; AM: *A. muricata*; and MF: *M. foliosa*

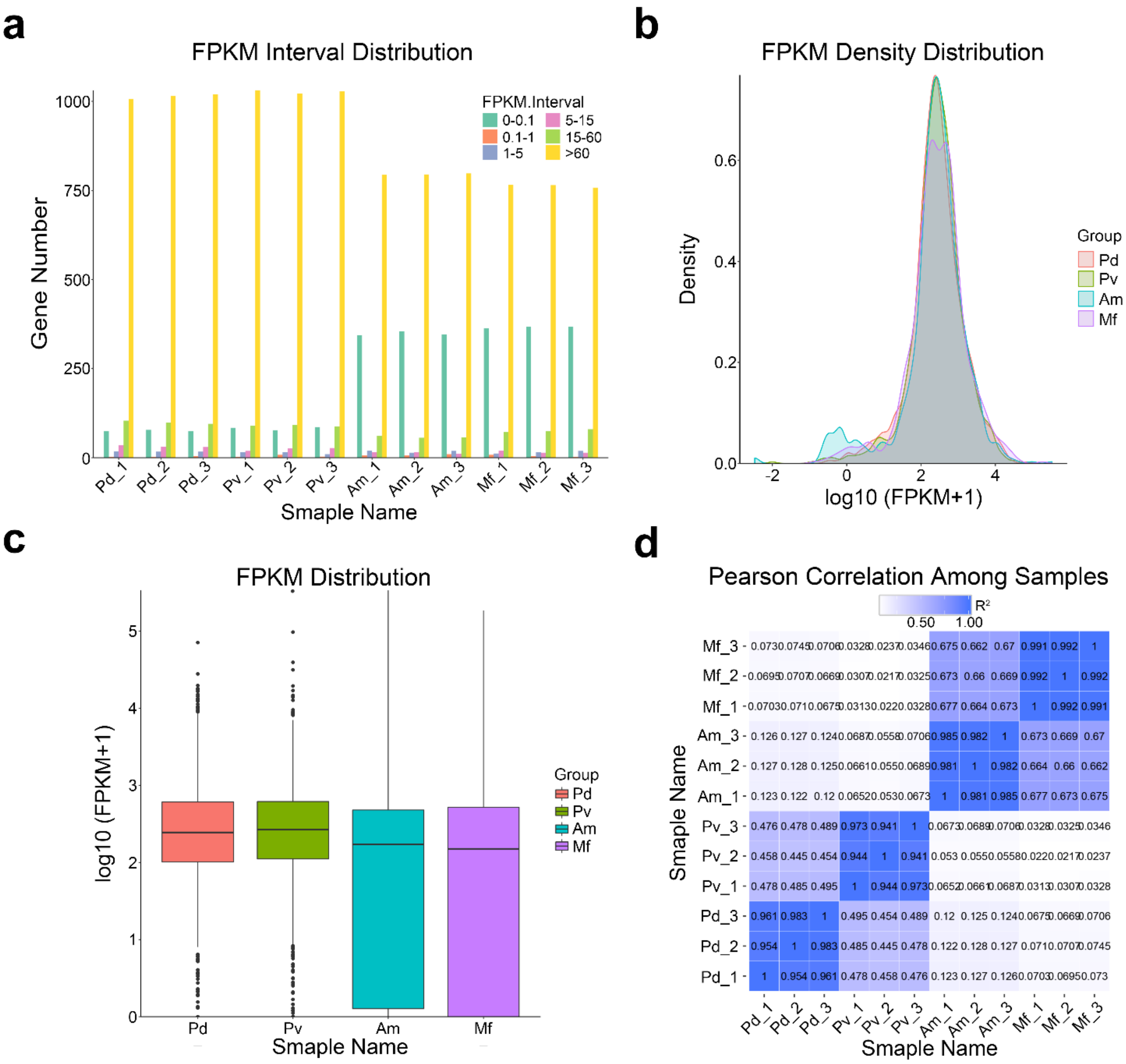


**Fig. S28** Summary of the Symbiodiniaceae gene expression level analysis. **a**. FPKM interval distribution. The horizontal axis shows the sample name, different colours represent FPKM intervals, and the vertical axis represents the number of genes in each interval. **b**. FPKM density distribution. The horizontal axis represents the log10 (FPKM+1) values, and the vertical axis represents the density of genes with different expression levels. **c**. FPKM box plot. The horizontal axis shows the sample name, and the vertical axis represents the log10 (FPKM+1) values. Each box plot shows five statistical values, including the maximum, upper quartile, median, lower quartile and minimum, from top to bottom. **d**. Pearson correlation among samples. The closer the value is to 1, the better the correlation. Pd: *P. damicornis*; Pv: *P. verrucosa*; Am: *A. muricata*; and Mf: *M. foliosa*.

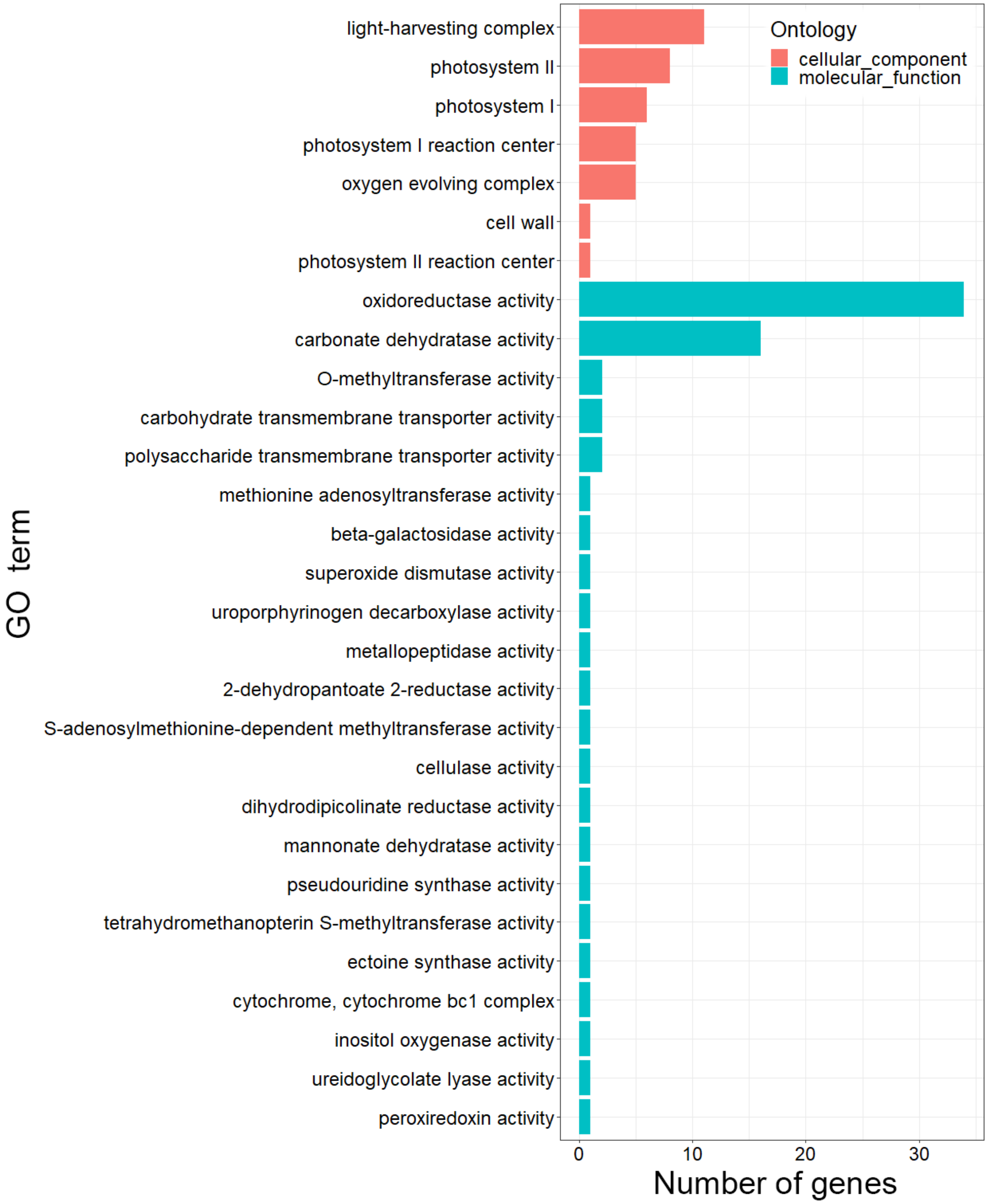


**Fig. S29** Bar graph of coexpressed Symbiodiniaceae sequences based on GO molecular functions and cellular components categories. The vertical axis represents the GO terms (orange represents molecular functions and blue-green represents cellular components), and the horizontal axis represents the number of transcripts annotated to the terms (including sub terms of the terms).

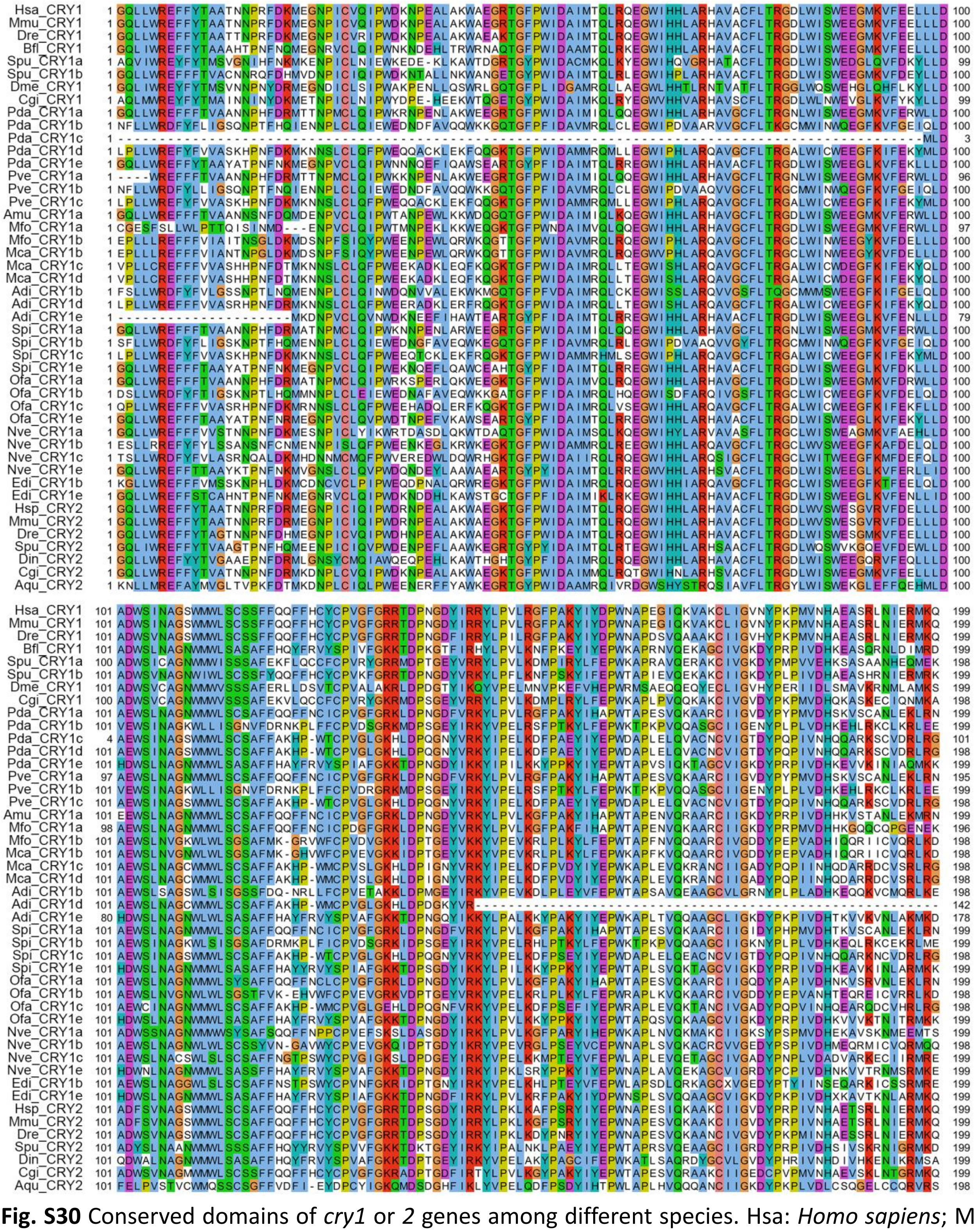

**Fig. S30** Conserved domains of *cry1* or *2* genes among different species. Hsa: *Homo sapiens*; Mmu: *Mus musculus*; Dre: *Danio rerio*; Bfl: *Branchiostoma floridae*; Spu: *Strongylocentrotus purpuratus*; Dme: *Drosophila melanogaster*; Cgi: *Crassostrea gigas*; Pda: *Pocillopora damicornis*; Pve: *Pocillopora verrucosa*; Amu: *Acropora muricata*; Mfo: *Montipora foliosa*; Mca: *Montipora capricornis*; Adi: *Acropora digitifera*; Spi: *Stylophora pistillata*; Ofa: *Orbicella faveolata*; Nve: *Nematostella vectensis*; Edi: *Exaiptasia diaphana*; Din: *Drosophila innubila*; and Aqu: *Amphimedon queenslandica*.

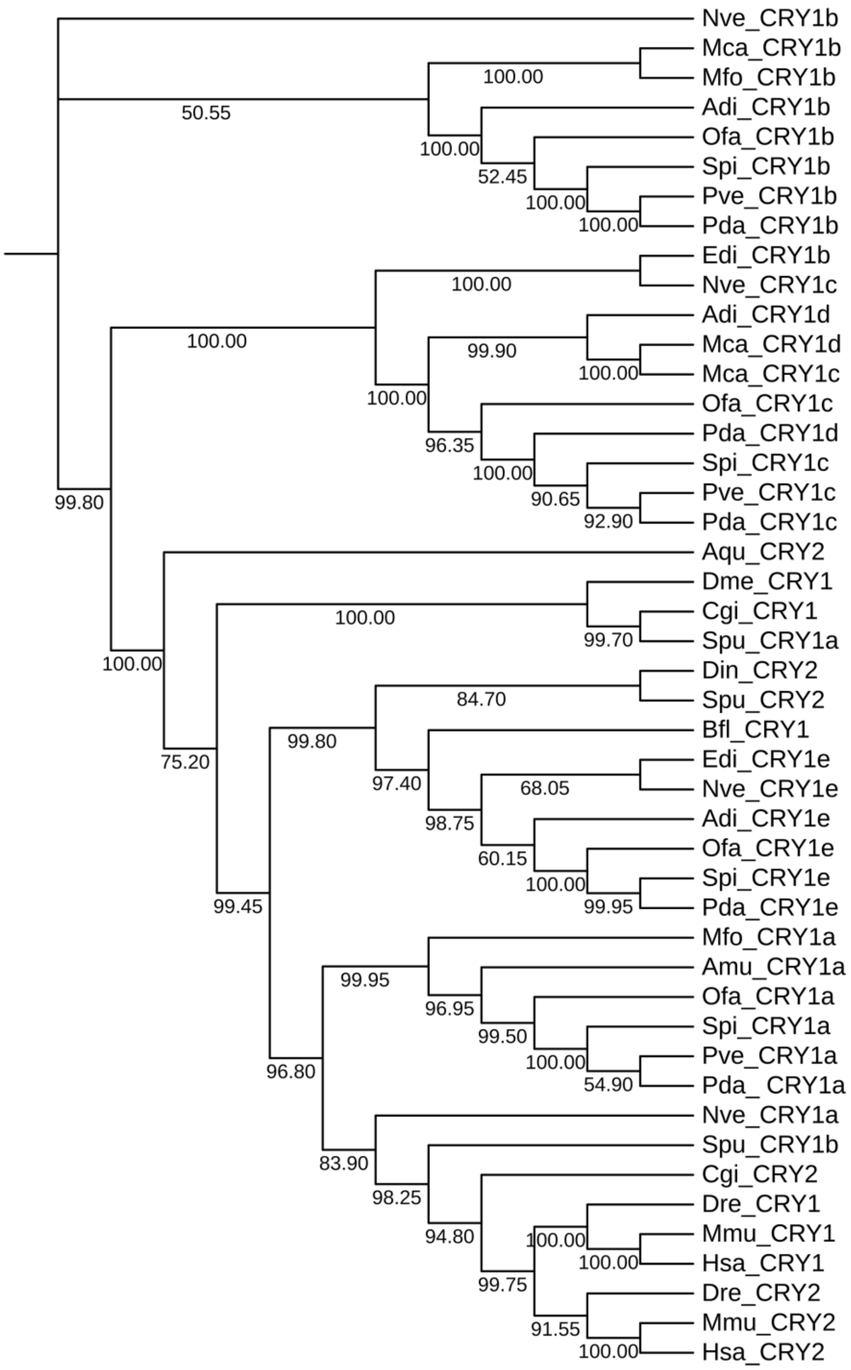


**Fig. S31** Phylogenetic tree of *cry1* or *2* genes based on the Neighbor-Joining method. Hsa: *Homo sapiens*; Mmu: *Mus musculus*; Dre: *Danio rerio*; Bfl: *Branchiostoma floridae*; Spu: *Strongylocentrotus purpuratus*; Dme: *Drosophila melanogaster*; Cgi: *Crassostrea gigas*; Pda: *Pocillopora damicornis*; Pve: *Pocillopora verrucosa*; Amu: *Acropora muricata*; Mfo: *Montipora foliosa*; Mca: *Montipora capricornis*; Adi: *Acropora digitifera*; Spi: *Stylophora pistillata*; Ofa: *Orbicella faveolata*; Nve: *Nematostella vectensis*; Edi: *Exaiptasia diaphana*; Din: *Drosophila innubila*; and Aqu: *Amphimedon queenslandica*.

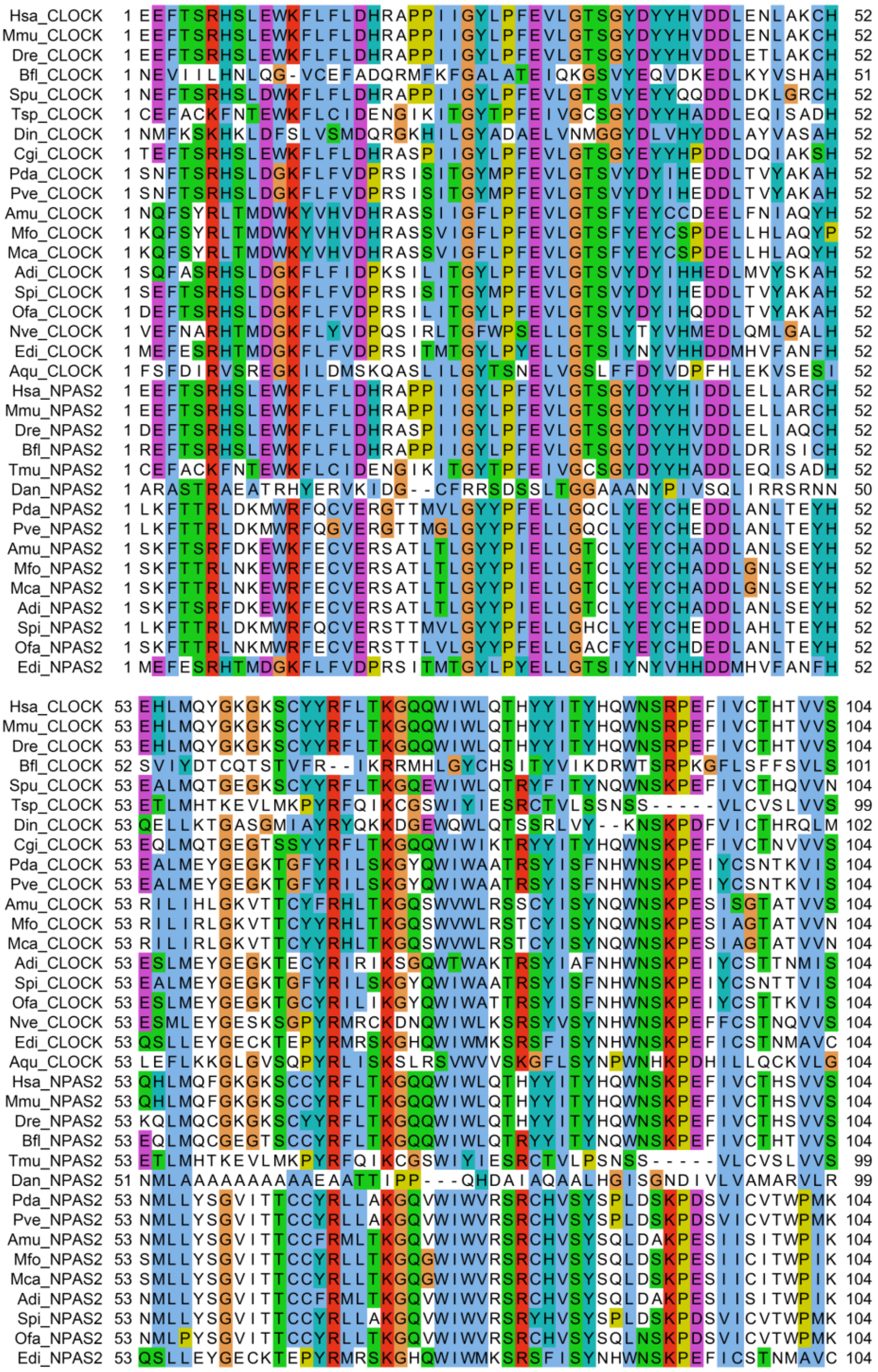

**Fig. S32** Conserved domains of *Clock* or *Npas2* genes among different species. Hsa: *Homo sapiens*; Mmu: *Mus musculus*; Dre: *Danio rerio*; Bfl: *Branchiostoma floridae*; Spu: *Strongylocentrotus purpuratus*; Tsp: *Trichinella spiralis*; Din: *Drosophila innubila*; Cgi: *Crassostrea gigas*; Pda: *Pocillopora damicornis*; Pve: *Pocillopora verrucosa*; Amu: *Acropora muricata*; Mfo: *Montipora foliosa*; Mca: *Montipora capricornis*; Adi: *Acropora digitifera*; Spi: *Stylophora pistillata*; Ofa: *Orbicella faveolata*; Nve: *Nematostella vectensis*; Edi: *Exaiptasia diaphana*; Aqu: *Amphimedon queenslandica*; Tmu: *Trichinella murrelli*; and Dan: *Drosophila ananassae*.

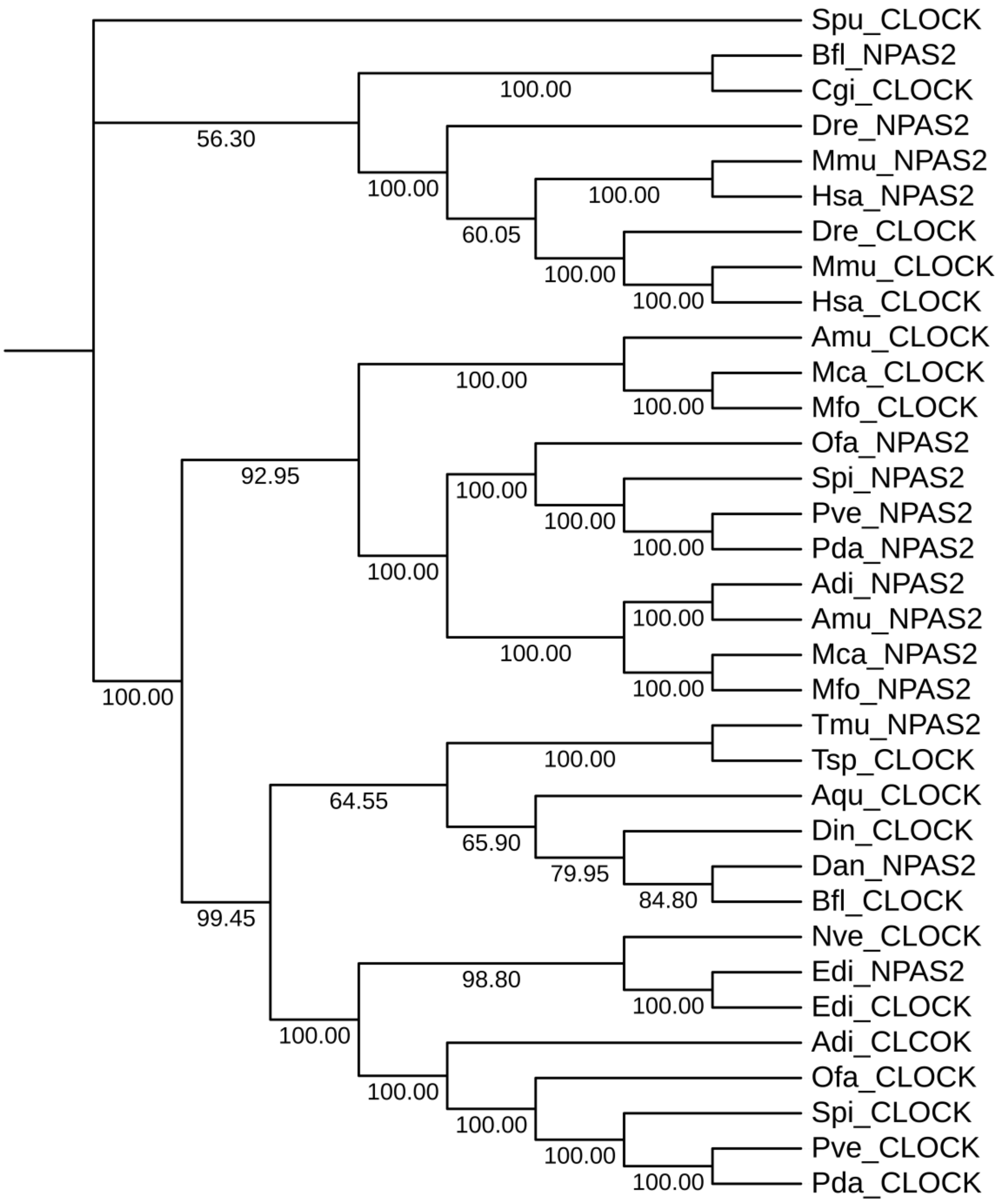


**Fig. S33** Phylogenetic tree of *Clock* or *Npas2* genes based on the Neighbor-Joining method. Hsa: *Homo sapiens*; Mmu: *Mus musculus*; Dre: *Danio rerio*; Bfl: *Branchiostoma floridae*; Spu: *Strongylocentrotus purpuratus*; Tsp: *Trichinella spiralis*; Din: *Drosophila innubila*; Cgi: *Crassostrea gigas*; Pda: *Pocillopora damicornis*; Pve: *Pocillopora verrucosa*; Amu: *Acropora muricata*; Mfo: *Montipora foliosa*; Mca: *Montipora capricornis*; Adi: *Acropora digitifera*; Spi: *Stylophora pistillata*; Ofa: *Orbicella faveolata*; Nve: *Nematostella vectensis*; Edi: *Exaiptasia diaphana*; Aqu: *Amphimedon queenslandica*; Tmu: *Trichinella murrelli*; and Dan: *Drosophila ananassae*.

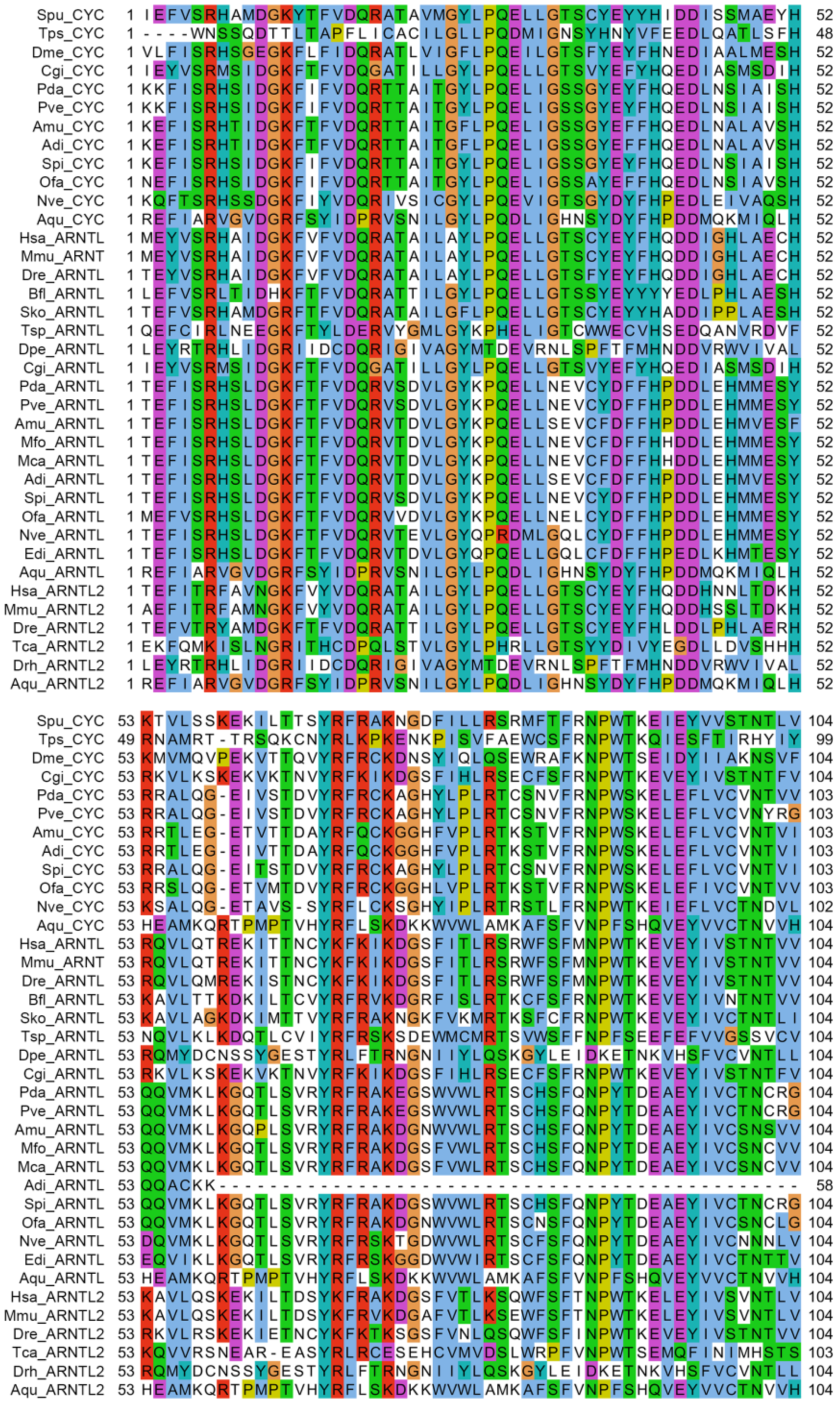

**Fig. S34** Conserved domains of *cyc* or *Arntl* genes among different species. Spu: *Strongylocentrotus purpuratus*; Tps: *Trichinella pseudospiralis*; Dme: *Drosophila melanogaster*; Cgi: *Crassostrea gigas*; Pda: *Pocillopora damicornis*; Pve: *Pocillopora verrucosa*; Amu: *Acropora muricata*; Adi: *Acropora digitifera*; Spi: *Stylophora pistillata*; Ofa: *Orbicella faveolata*; Nve: *Nematostella vectensis*; Aqu: *Amphimedon queenslandica*; Hsa: *Homo sapiens*; Mmu: *Mus musculus*; Dre: *Danio rerio*; Bfl: *Branchiostoma floridae*; Sko: *Saccoglossus kowalevskii*; Tsp: *Trichinella spiralis*; Dpe: *Drosophila persimilis*; Mfo: *Montipora foliosa*; Mca: *Montipora capricornis*; and Edi: *Exaiptasia diaphana*.

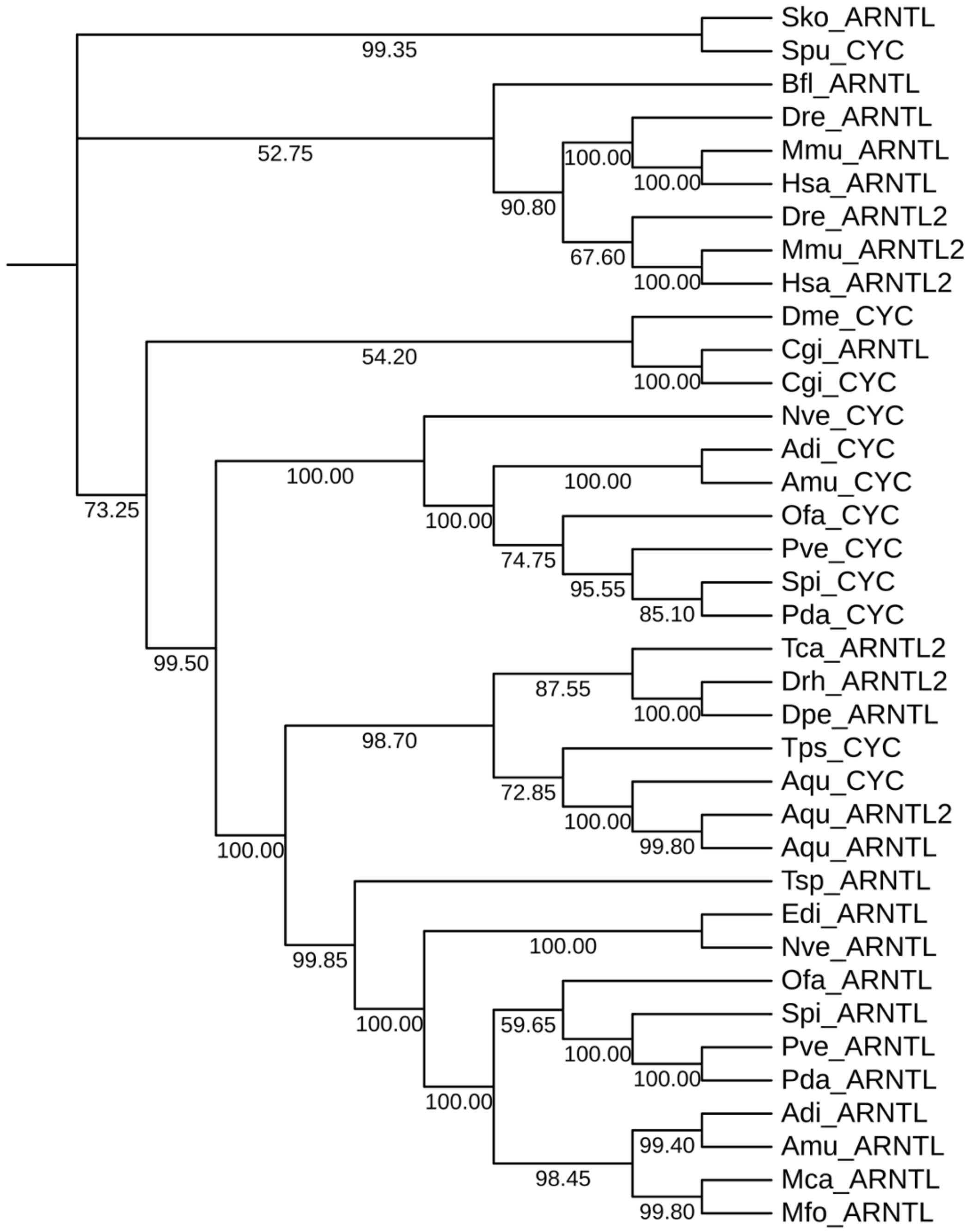


**Fig. S35** Phylogenetic tree of *cyc* or *Arntl* genes based on the Neighbor-Joining method. Spu: *Strongylocentrotus purpuratus*; Tps: *Trichinella pseudospiralis*; Dme: *Drosophila melanogaster*; Cgi: *Crassostrea gigas*; Pda: *Pocillopora damicornis*; Pve: *Pocillopora verrucosa*; Amu: *Acropora muricata*; Adi: *Acropora digitifera*; Spi: *Stylophora pistillata*; Ofa: *Orbicella faveolata*; Nve: *Nematostella vectensis*; Aqu: *Amphimedon queenslandica*; Hsa: *Homo sapiens*; Mmu: *Mus musculus*; Dre: *Danio rerio*; Bfl: *Branchiostoma floridae*; Sko: *Saccoglossus kowalevskii*; Tsp: *Trichinella spiralis*; Dpe: *Drosophila persimilis*; Mfo: *Montipora foliosa*; Mca: *Montipora capricornis*; and Edi: *Exaiptasia diaphana*.

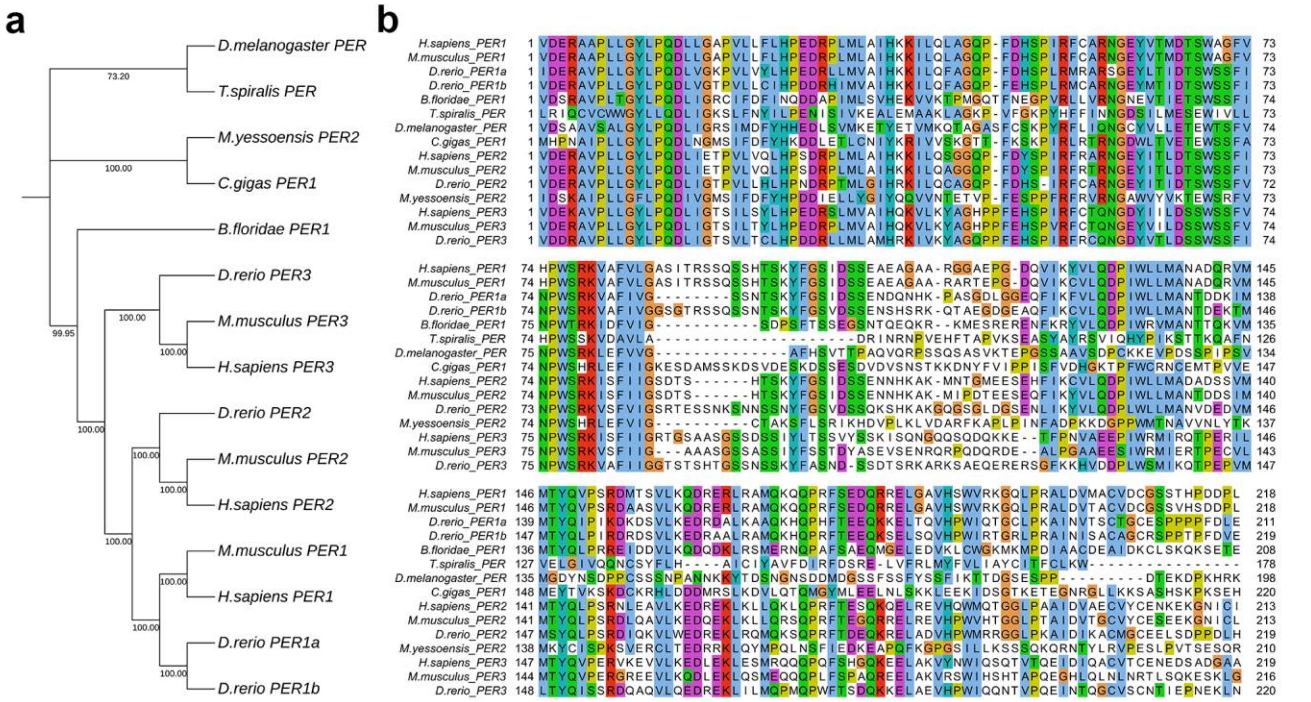


**Fig. S36** *Per1* or *2* or *3* gene phylogenetic analysis. **a.** Phylogenetic tree of *per* genes based on the Neighbor-Joining method. **b.** Conserved domains of *per* genes among different species. *H.sapiens*: *Homo sapiens*; *M.musculus*: *Mus musculus*; *D.rerio*: *Danio rerio*; *B.floridae*: *Branchiostoma floridae*; *T.spiralis*: *Trichinella spiralis*; *D.melanogaster*: *Drosophila melanogaster*; *C.gigas*: *Crassostrea gigas*; and *M.yessoensis*: *Mizuhopecten yessoensis*.
